# A breast cancer patient-derived xenograft and organoid platform for drug discovery and precision oncology

**DOI:** 10.1101/2021.02.28.433268

**Authors:** Katrin P. Guillen, Maihi Fujita, Andrew J. Butterfield, Sandra D. Scherer, Matthew H. Bailey, Zhengtao Chu, Yoko S. DeRose, Ling Zhao, Emilio Cortes-Sanchez, Chieh-Hsiang Yang, Jennifer Toner, Guoying Wang, Yi Qiao, Xiaomeng Huang, Jeffery A. Greenland, Jeffery M. Vahrenkamp, David H. Lum, Rachel E. Factor, Edward W. Nelson, Cindy B. Matsen, Jane M. Poretta, Regina Rosenthal, Anna C. Beck, Saundra S. Buys, Christos Vaklavas, John H. Ward, Randy L. Jensen, Kevin B. Jones, Zheqi Li, Steffi Oesterreich, Lacey E. Dobrolecki, Satya S. Pathi, Xing Yi Woo, Kristofer C. Berrett, Mark E. Wadsworth, Jeffrey H. Chuang, Michael T. Lewis, Gabor T. Marth, Jason Gertz, Katherine E. Varley, Bryan E. Welm, Alana L. Welm

**Author notes:** equal contribution (Katrin P. Guillen, Maihi Fujita, Andrew J. Butterfield). equal contribution and corresponding authors.

## Abstract

Model systems that recapitulate the complexity of human tumors and the reality of variable treatment responses are urgently needed to better understand cancer biology and to develop more effective cancer therapies. Here we report development and characterization of a large bank of patient-derived xenografts (PDX) and matched organoid cultures from tumors that represent some of the greatest unmet needs in breast cancer research and treatment. These include endocrine-resistant, treatment-refractory, and metastatic breast cancers and, in some cases, multiple tumor collections from the same patients. The models can be grown long-term with high fidelity to the original tumors. We show that development of matched PDX and PDX-derived organoid (PDxO) models facilitates high-throughput drug screening that is feasible and cost-effective, while also allowing *in vivo* validation of results. Our data reveal consistency between drug screening results in organoids and drug responses in breast cancer PDX. Moreover, we demonstrate the feasibility of using these patient-derived models for precision oncology in real time with patient care, using a case of a triple negative breast cancer with early metastatic recurrence as an example. Our results uncovered an FDA-approved drug with high efficacy against the models. Treatment with the PDxO-directed therapy resulted in a complete response for the patient and a progression-free survival period more than three times longer than her previous therapies. This work provides valuable new methods and resources for functional precision medicine and drug development for human breast cancer.

**Graphical Abstract:** 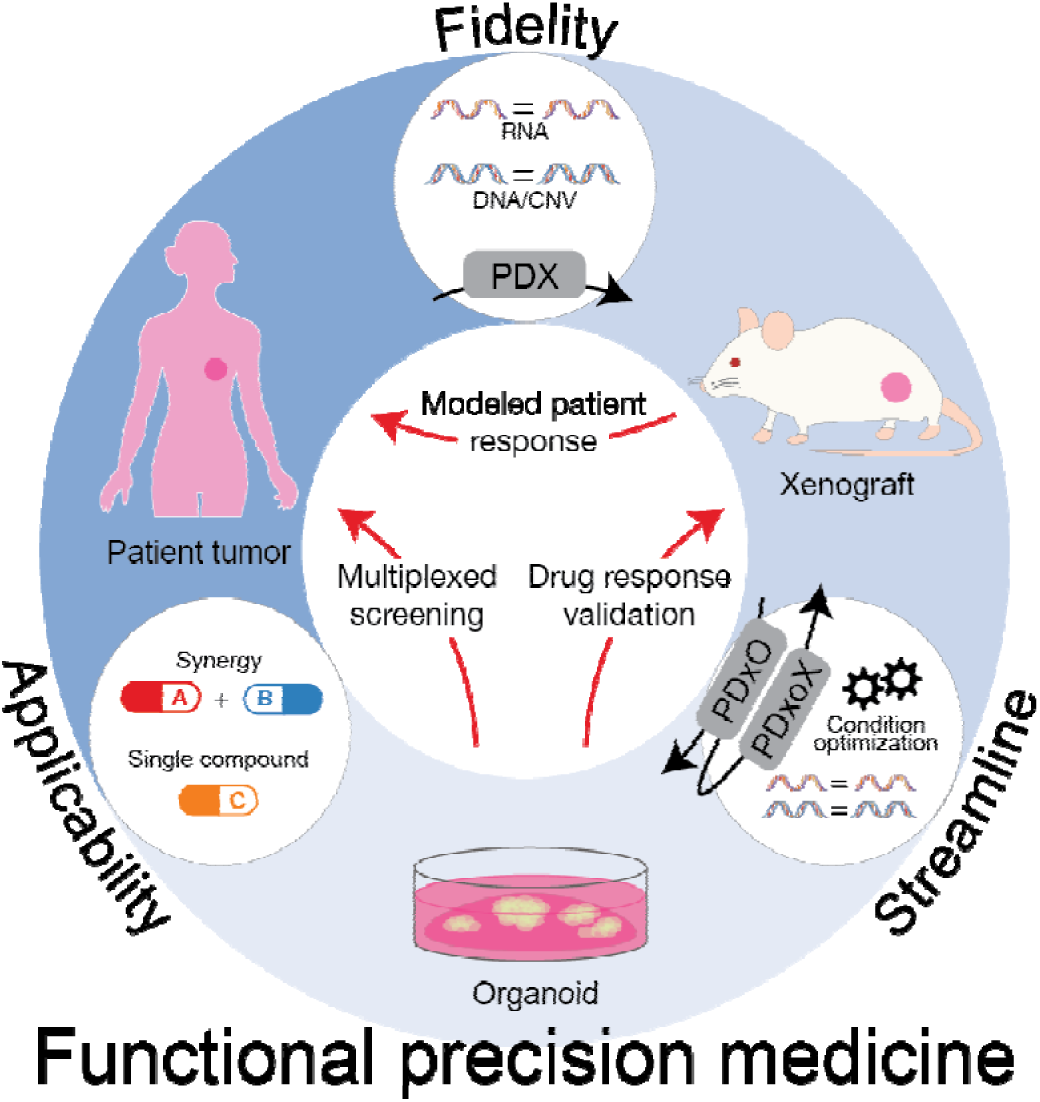

## Introduction

The immense heterogeneity of human cancers has hampered the development of cancer therapies and contributes to the limited success of drug treatment. Model systems that recapitulate the reality of variable treatment responses need to be utilized for more precise drug development and testing. In fact, the next large strides in cancer treatment success may depend on precision approaches that take into account the diverse nature of individual tumors when choosing treatments.

To model solid tumors from cancer patients more accurately, our group and others have developed and exploited patient-derived xenografts (PDX), whereby fragments of human tumors are implanted directly into immune-deficient mice and grown in a serially-transplantable manner. PDX models recapitulate human tumors with relatively high fidelity^1^ and exhibit treatment responses that are concordant with those observed in the patients from which they are derived (recently reviewed^2^). While imperfect, PDX models are currently the most robust way to model diverse human tumors in the laboratory *in vivo*.

PDX models are used for preclinical, co-clinical, and clinical research studies. The use of PDX models for precision medicine has been pioneered in programs such as The Mouse Hospital and Co-Clinical Trial Project, using PDX models to tailor patient-specific therapies for various cancer types^3–4^. As a research tool, PDX models can also be used to interrogate drug responses and mechanisms of resistance; study tumor heterogeneity and evolution; and model metastatic disease^2^. Whether used for precision oncology or as key research tools, one major shortcoming of PDX-based studies, however, is that they are limited by high cost and low throughput.

For several solid tumor types, development of three-dimensional (3D) organoid models from patient tumors and PDX models is now feasible, and may be more representative of human cancer than two-dimensional (2D) cultures^5^. Evidence is accumulating to indicate that patient-derived organoids (PDO) show strong biological concordance with the tumors from which they are derived. Pancreatic and colorectal tumors, in particular, have been extensively modeled using patient-derived organoids^6–7^. Using whole genome analysis, Gendoo et al. reported good concordance between patient tumors, PDX, and PDO in pancreatic cancer^8^. Another study in pancreatic cancer supported this notion and showed that organoids, although primarily clonal, maintain distinct patient phenotypes and respond differently to drug combinations^9^. Patient-derived pancreatic cancer or colorectal cancer organoids are now being used to predict therapeutic responses, and facilitate precision medicine for patients^10, 11^. PDO or PDX-derived organoids (PDxO) have also been described for hepatocellular carcinoma^12^, hepatoblastoma^13^ and glioblastoma^14^, as well as prostate^15, 16^; bladder^17^; ovarian^18–19^; breast^20^; gastric^21^; lung^22, 23^; esophageal^24^; kidney^25^; and head and neck^26, 27^ cancers.

Genomic testing for precision oncology is becoming more mainstream, and can personalize cancer therapy for better outcomes. In a recent study of 429 evaluable patients with diverse malignancies, molecular analysis (generally next-generation DNA sequencing, although some cases also included mRNA analysis, immunohistochemistry, and/or immunotherapy-associated markers) revealed that 62% of patients had mutations that matched to at least one drug, and 20% had mutations that matched to multiple drugs including combination therapies. When compared to the remaining 38% of patients who received physician’s choice of drug (unmatched or low-match cases), patients who received the entire regimen recommended by the molecular tumor board (all matching drugs) had longer progression-free survival times^28^. However, accumulating data suggests that functional drug testing using patient-derived models may hold distinct advantages over use of genomics alone to personalize therapy. In a larger study of 769 patients suffering from various cancer types, genomics alone identified viable therapeutic options for <10% of patients with advanced disease, with <1% success rate for a match to an approved therapy^29^. However, organoids or PDX could be grown from 38% of these cases, even with suboptimal tissue collection procedures. As a proof-of-concept, models from four of the cases were tested with combined genomic testing and functional drug testing, and in all four cases effective targeted agents and combinations were identified. Although drug responses could often be related back to genomic findings, in half of the cases functional screening clearly identified different drug responses despite similar driver mutations.

Breast cancer is particularly challenging when it comes to identifying successful therapies based on genomic alterations. An ever-increasing number of genetic and epigenetic drivers are being uncovered^30, 31^ but in metastatic breast cancer, which represents the major unmet medical need in this disease, the molecular heterogeneity is vast and represents an impediment to identification and development of successful targeted therapies^32^. Although clinically-actionable mutations can be identified in 40-46% of cases^31, 33^, no clinical benefit was realized from matching therapies to these variants in a recent clinical trial in metastatic breast cancer^33^. These data suggest that parallel functional modeling of candidate driver gene dependence and drug response may be required to achieve improved outcomes for breast cancer patients.

Hundreds of PDX models have been developed for breast cancer that retain much of the molecular diversity of this disease^34^. However, there remains a shortage of models representing some of the deadliest breast cancers. These include drug-resistant tumors, especially those derived from metastases, endocrine-resistant estrogen receptor-positive (ER+), and HER2+ tumors. A larger biobank of advanced breast cancer models, as well as *in vitro* methods to propagate these tumors for more feasible experimental manipulation, is necessary to accelerate progress toward understanding sensitivity and resistance to all types of therapies, across diverse breast cancer subtypes.

Short term 2D cultures of human breast cancer cells derived from PDX models have been shown to have reproducible responses to therapies that recapitulate tumor responses *in vivo*^35, 36^; however, long term cultures derived from PDX are desirable for mechanistic studies of tumor cell biology and drug response or resistance. The ability to run companion *in vivo* studies is also ideal. Here, we established a large collection of paired PDX and PDX-derived organoid (PDxO) models with high fidelity to their original tumor samples, and developed medium-throughput 3D drug screening techniques across a range of breast cancer subtypes. We also demonstrated feasibility for combined genomic and functional precision drug selection approaches in the clinical setting, in a case of triple-negative breast cancer with early metastatic recurrence, where we found that drug selection on the basis of functional drug screening yielded meaningful clinical benefit.

## Results

### Generation of new breast cancer PDX models representing the deadliest forms of the disease

We previously reported that engraftment of human breast cancer specimens into immune-deficient mice as PDX recapitulated human breast tumor characteristics, including metastatic behavior and clinical outcomes^37^. We continue to generate breast cancer PDX models, now with an emphasis on those representing the greatest unmet medical and research needs: ER+ endocrine-resistant tumors; ER+ and HER2+ co-expressing tumors; unusually aggressive tumor types (e.g. metaplastic breast cancer); drug-resistant tumors; and PDXs representing primary-metastatic pairs or longitudinal collections from the same patient. A summary of our current collection is shown in **Table 1** with more detail, including de-identified clinical information, in **Table S1.** As an extension of our previously published models (HCI-001 through HCI-012^37^), models include fourteen metastatic, drug-resistant, triple-negative breast cancers (TNBCs); ten metastatic, endocrine-resistant ER+ breast cancers (two of lobular subtype and four with matched estrogen-independent sublines); four rare TNBC types (a malignant phyllodes tumor and three metaplastic breast cancers); three drug-resistant primary TNBC tumors and five drug-resistant locally-recurrent breast tumors (four TNBC and one triple positive breast cancer); and sixteen untreated primary tumors (three ER+, one HER2+, and twelve TNBC). Importantly, three of the TNBC and one of the ER+ untreated primary tumors are matched to metastatic samples from the same patients, also grown as a PDX. Several of the latter TNBC patients are participants in our active “TOWARDS Precision Medicine In Breast Cancer” clinical study to use PDX to predict recurrence and inform patient care. Sixteen lines in our collection are primary-metastatic pairs or longitudinal collections from the same patient over time. Finally, we report two sub-lines derived from spontaneously metastatic lesions in the mouse.

**Table 1.**
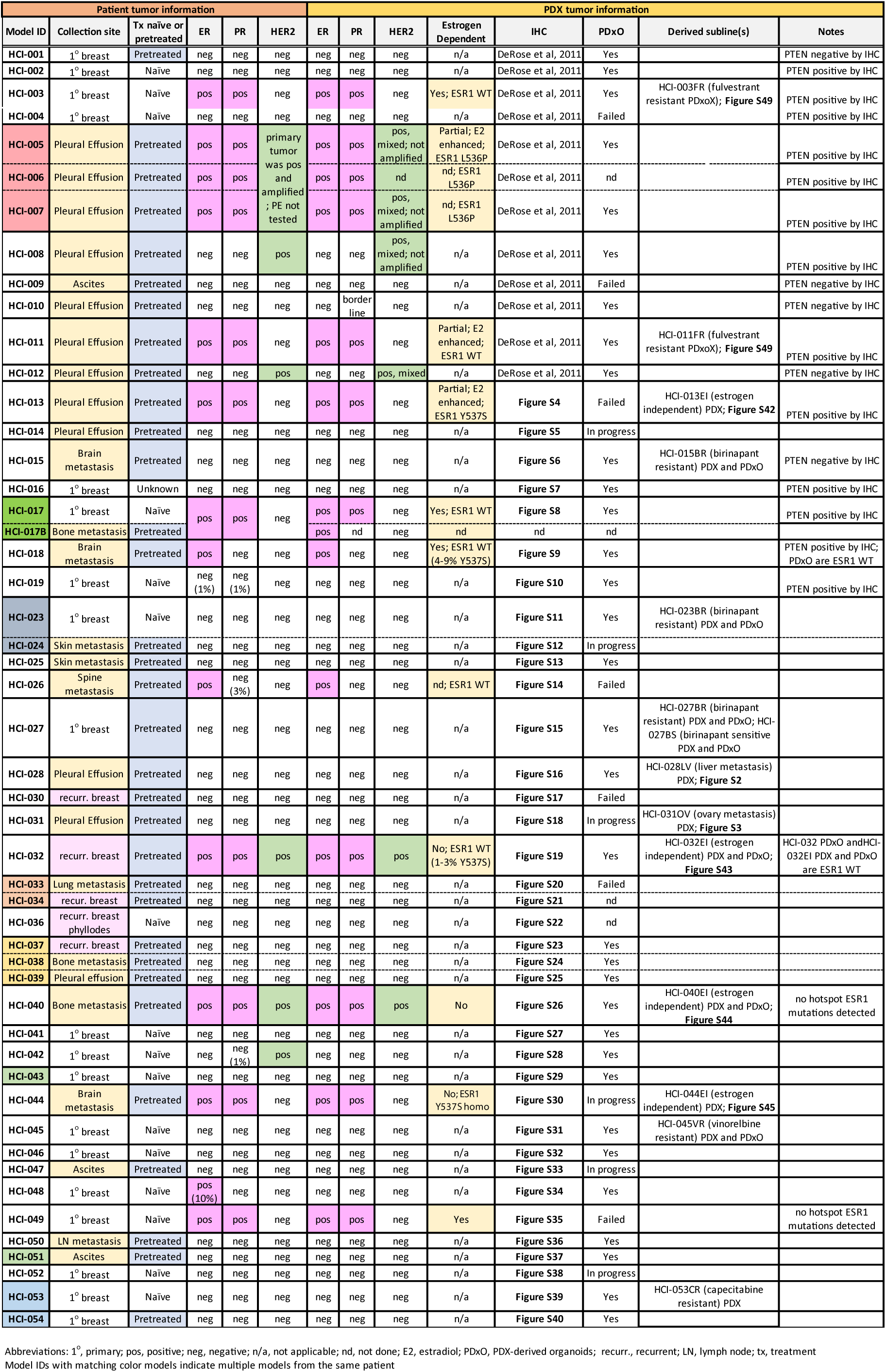

Our overall “take rate,” defined as successful growth of PDX for at least two serially-transplanted generations, was 29%. We found primary tumors to be more difficult to engraft (25% take rate for 102 attempts) compared to tumors derived from metastatic sites (36% take rate for 50 attempts). ER+ PDX were the most difficult to develop, with a take rate of only 9% for primary ER+ tumors (n=32 attempts) and 16% for metastatic ER+ tumors (n=32 attempts). Take rates for HER2+ primary tumors were 25% (n=8 attempts), as compared to 33% for HER2+ metastatic tumors (n=6 attempts). ER+ and HER2+ tumor numbers and take rates were calculated independent of coexpression of HER2 and ER. TNBC tumors were the easiest to establish, with a take rate of 58% for TNBC primary tumors (n=12 attempts) and 85% for TNBC metastases (n=13 attempts). These rates are within the historical range reported by other groups^34^. While all of the above attempts were from primary or metastatic surgical resections or body fluids such as pleural effusions or ascites, we also established PDX from several TNBC primary tumor biopsies for the TOWARDS study. We found take rates for these TNBC primary tumor biopsy samples to be approximately half that of the TNBC primary tumor surgical samples described above (29%; n=56 attempts from primary tumor biopsies). We and others previously reported that positive engraftment of a primary breast tumor as a PDX is an independent predictor of metastatic relapse and poor outcome of the patient^37, 38^, suggesting that PDXs represent the most aggressive breast cancers, and the ones for which more research is needed. Indeed, to date, at least 30/42 (71%) of the patients from which we successfully generated a PDX have died of breast cancer.

Each PDX line was systematically “credentialed” through a rigorous process (**Figure S1**). Each line was tested for human and mouse pathogens (see Methods), including *Corynebacterium bovis* (*C. bovis*) and lactate dehydrogenase elevating virus (LDEV). LDEV specifically infects mouse macrophages and is a common contaminant in PDX lines. Some of our early lines (HCI-010, HCI-013, HCI-013EI) tested positive for LDEV, and we removed this pathogen by flow cytometry sorting for human epithelial cells and/or passaging the PDX through immune-deficient rats for one passage (see Methods). We did not find obvious differences in gene expression following sorting to remove LDEV (not shown). Once confirmed to be negative for known pathogens, all PDX models were validated further by immunohistochemistry staining to be positive for breast epithelial markers and human mitochondria, and negative for the mouse and human lymphoma marker CD45 (**Figures S2-S40**). ER, PR, and HER2 staining was conducted and compared to the original tumor material and/or clinical pathology reports with concordant results (**Tables 1 and S1**).

PDX models were also characterized through DNA mutation and copy number variant (CNV) analysis, as well as RNA sequencing. Genomics analysis revealed that PDX models reflected the heterogeneity of human breast cancer with respect to both driver mutations and intrinsic subtypes. For mutation analysis, we selected known driver genes from previous TCGA analyses^39, 40^. Mutations in selected genes were restricted to missense, nonsense, nonstop, frameshift, and splice mutations (see Methods). As expected, the most common variants were in *TP53* and *PIK3CA* (**Figure 1a**, top panel). Although no sequencing data was reported in our previous manuscript with the 12 initial PDX lines, we had reported CNV data^37^. Those CNV data were not analyzed using the new PDXNet standards^1^ and reflect older technology. We have therefore repeated CNV analysis using newer technology. We noted other well-characterized lesions such as loss of *PTEN* and *RB1,* and amplification of the *MYC* locus (**Figure 1a**, lower panel). RNA-seq analysis of PAM50 genes^41^ revealed that our collection comprises a variety of breast cancer subtypes (**Figure 1b**). Analysis of the genomic relationship between PDX lines and their metastatic sublines, when available, was also examined. HCI-028LV (**Figure S2**) was derived from a metastasis to the mouse liver that arose spontaneously from the HCI-028 TNBC PDX growing in the mammary fat pad. HCI-028 was derived from the pleural fluid of a patient who developed liver, bone, ovary, and brain metastases. HCI-031OV (**Figure S3**) was derived from a spontaneous metastasis to the mouse ovary from the HCI-031 lobular TNBC PDX. HCI-031 was derived from the pleural fluid of a patient that developed metastases in fallopian tubes, bones, pleura, liver, and brain. We found that in both cases the metastatic sublines retained the same genomic driver mutations and have similar gene expression profiles to their parental PDX lines (**Figure 1, Figures S1 and S2**), and both metastatic sublines spontaneously metastasize back to the organ from which they were derived (liver and ovary, respectively). The metastatic profiles of each PDX model, and the corresponding patient, when known, are shown in **Table S1**.

**Figure 1.**
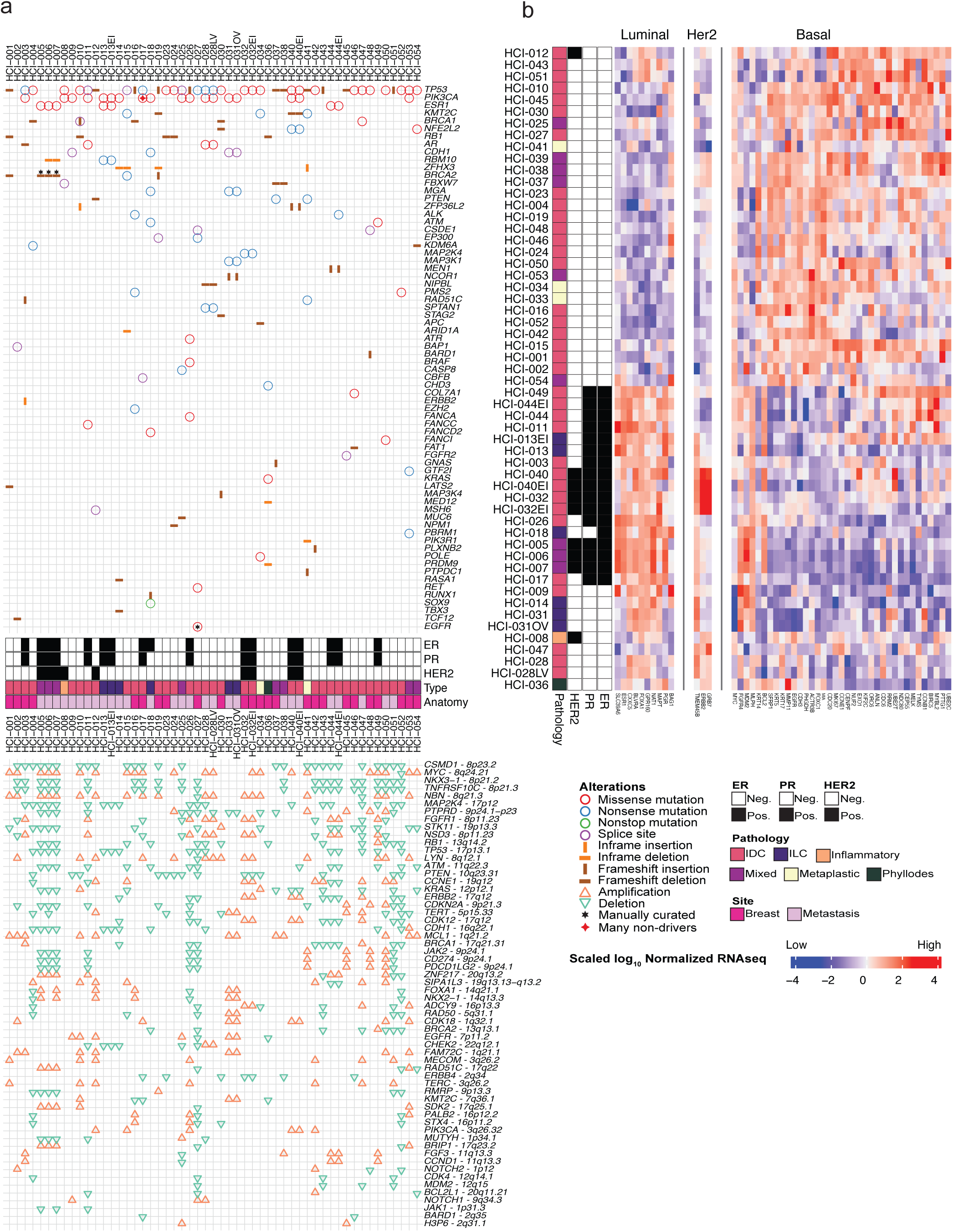
Genomic characterization of breast cancer PDX models. **(a) Top:** Oncoprint plot showing single nucleotide variants and indels for commonly mutated genes in cancer (top). Annotations for each model include hormone receptor status for ER and PR; HER2 status; pathology: IDC (invasive ductal carcinoma), mixed, phyllodes, ILC (invasive lobular carcinoma), inflammatory, or metaplastic; and whether the sample was from the primary breast tissue or metastatic site. **Bottom:** Specific gene level copy number alterations are displayed. **(b)** Unsupervised clustering of the PDX models was performed using root-mean-squared scaling of transcript abundance in the PAM50 geneset.

### New models of ER+ breast cancer with features of endocrine resistance

As we and others have reported^1, 34, 37, 42, 43^, the molecular and pathophysiological features of breast PDX are representative of the human breast cancers from which they arise. Growth of ER+ breast cancers is very challenging, due in part to their relatively slow growth rate and to their dependence on estrogen, which is present at lower levels in mice than in humans^44^. It is clear from prior work that some ER+ tumors require supplementation with estradiol^37, 45^. To avoid urine retention and cystitis caused by supraphysiological doses of estrogen in some strains of mice^46^, we lowered the 0.8 mg dose of supplemental estradiol (E2) given in our beeswax E2 supplementation pellets^47^, and still retained estrogen-dependent growth of ER+ PDXs. For extremely estrogen-dependent lines such as HCI-018 (ER+ metastatic lobular carcinoma), the combination of short-term E2 pellet and low dose E2 maintenance in drinking water was the only way to reproducibly sustain tumor growth (**Figure S41**). Thus, we establish and maintain all standard ER+ PDX lines with subcutaneous 0.4 mg E2 pellets placed at the time of tumor implantation, followed by E2 maintenance in the drinking water from four weeks after tumor implantation to the end of the experiment (see Methods).

The dependence of each ER+ PDX model on E2 was tested by attempting to grow established ER+ PDX tumors in the cleared mammary fat pads of ovariectomized mice with no E2 supplementation. Tumors that grew in estrogen-deprived conditions were considered E2-independent, and sublines were generated with the “-EI” designation (**Table 1 and Figures S42-45**). Estrogen-independence or endocrine resistance in metastatic breast cancer has been attributed to mutations in *ESR1*, which encodes ERα. We examined the mutation status of *ESR1* in all of our ER+ models by droplet digital PCR for hotspot mutations (Y537S/C/N and D538G). Five out of nine (56%) of metastatic ER+ breast PDX models from seven different patients contained an *ESR1* mutation (**Table 1**). One model, HCI-044, carries a homozygous Y537S mutation, as does the matching patient tumor (**Figure S46A**). We found two cases where no *ESR1* mutations were detected in the patient sample, but the Y537S mutation was detected at low frequency (<10%) in the PDX, in both DNA and RNA samples. These include HCI-018, from a brain metastasis (allele frequency of 4.3% in DNA and 9.3% in RNA; **Figure S46B**) and HCI-032, from a locally recurrent breast tumor (allele frequency of 1% in DNA and 3% in RNA; **Figure S46C**). The discrepancy between the patient sample and PDX is likely due to heterogeneity between the patient tumor fragment that was used to make PDX versus the fragment used for DNA/RNA isolation. Heterogeneity of *ESR1* mutations in patient samples have previously been reported, along with retention of the heterogeneity in the matching PDX^43^. Alternatively, the low frequency mutation in the PDX models could reflect evolution of the tumor upon engraftment in mice, but the low allele frequency does not suggest strong selective pressure. To this point, selection of the estrogen-independent subline of HCI-032 (HCI-032EI; **Figure S43**) in ovariectomized mice did not result in any Y537S mutations detected by ddPCR (not shown).

### Generation of paired PDX and PDX-derived organoid models for multiple breast cancer subtypes

In order to generate matched *in vivo* and *in vitro* patient-derived models of breast cancer, we sought to grow long-term organoid cultures from each of our PDX models. To theoretically minimize alterations in the biology of individual tumors, we optimized *in vitro* conditions that could promote long-term growth of PDX tumor-derived organoids (PDxO) in 3D culture with the fewest medium supplements possible. We examined various medium conditions, and assessed PDxO growth using a variety of methods: the area occupied by live cells in the organoids in an entire well, morphology of individual organoids, and metabolic activity (intracellular ATP content) of organoids. We used HCI-002, a treatment-naive TNBC PDX, as an initial model for experimentation. By testing various additives often used in organoid cultures, we found that the addition of a Rho kinase inhibitor (Y-27532) was sufficient to support growth of freshly isolated HCI-002 PDxO over 15 days of initial culture (**Figure 2a-b; Figure S47a**). Addition of other common organoid culture supplements did not enhance the beneficial effect of Y-27632. In fact, some of the supplements decreased the live cell content of organoids, indicating a detrimental effect on survival and outgrowth (**Figure 2a-b; Figure S47b**). Culturing other freshly isolated TNBC PDxO lines in the conditions optimized for HCI-002 PDxO and with other supplements also showed that the critical medium supplement was Y-27632 (**Figure S47b,d**). While some TNBC organoid lines grew slower in our basal conditions compared to a growth medium previously described by Sachs et al.^20^, others grew at similar rates (**Figure S47e**), indicating that both methods appear to be sufficient to support the PDX-derived organoid cultures. Since the medium described by Sachs et al. contains many additives to stimulate or inhibit various signaling pathways, we chose to move forward with the simpler, Y-27632-supplemented medium as our “standard” basal condition for establishment and maintenance of PDxO lines, with the rationale to try to maintain endogenous signaling within the tumors as much as possible.

**Figure 2.**
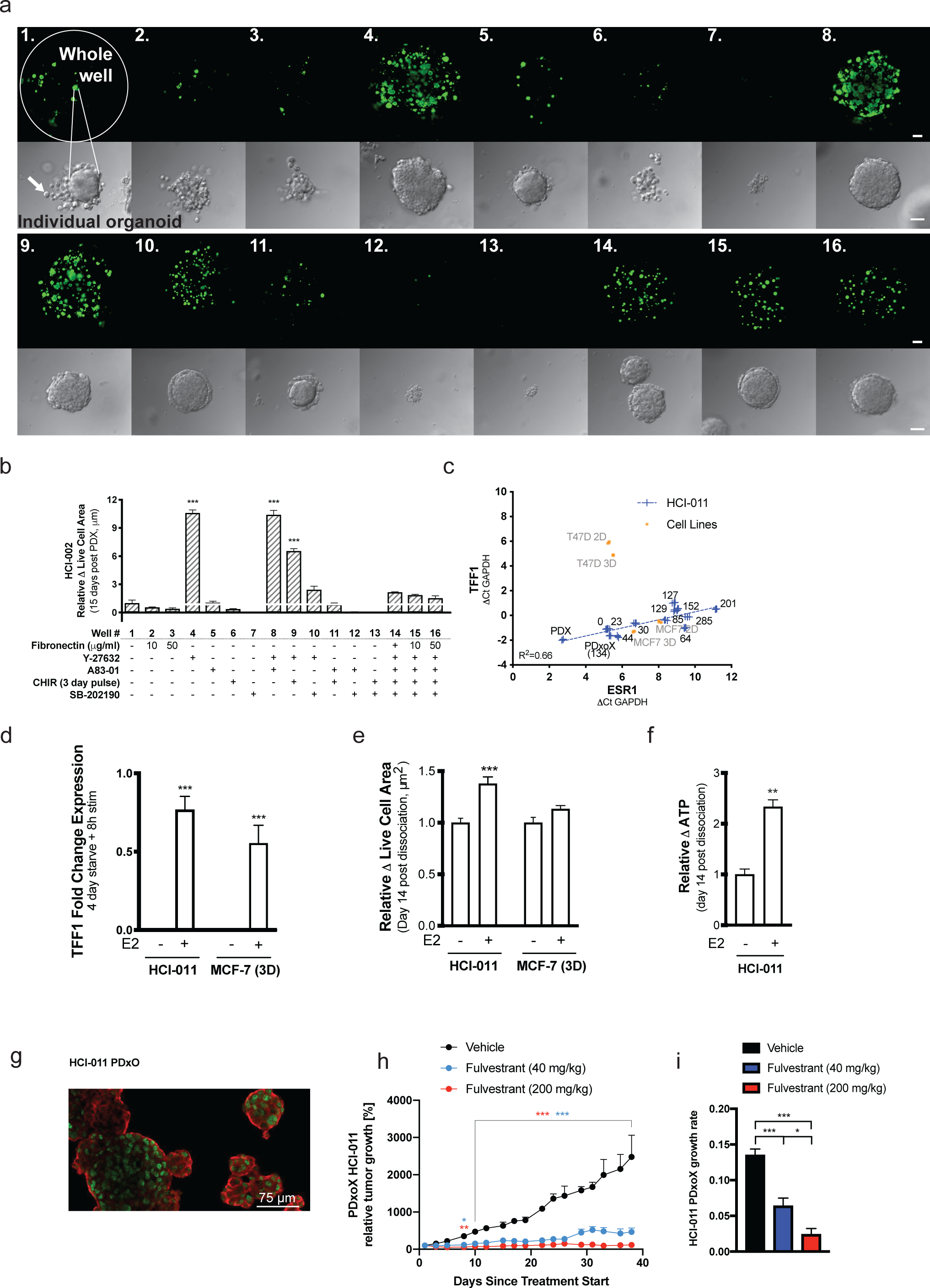
Optimization of PDxO culture conditions. **(a)** Live cell area of entire wells (upper row) and brightfield images of individual organoids (lower row) of HCI-002 PDxOs grown in 16 different conditions, 14 days post-organoid preparation. Scale bars indicate 500 um (upper row) and 50 um (lower row). **(b)** Quantified live cell area of HCI-002 PDxOs grown in 16 different culture conditions. Data are normalized to the control condition. **(c)** Expression level of *ESR1* and its downstream target *TFF1* in the ER^+^ PDxO line HCI-011 over time in culture compared to HCI-011 PDX, PDxoX or cell lines MCF7 or T47D cultured in 2D or 3D for 6 days. Numbers associated with each data point represent the sample’s total number of days in culture. Data are normalized to human GAPDH Ct. **(d)** TFF1 expression in ER^+^ HCI-011 PDxOs and 3D cultures of MCF7 stimulated with E2 for 8h after 4 days of culturing in phenol red-free medium with charcoal-stripped FBS. Data are normalized to human GAPDH Ct and control condition. **(e)** Quantified live cell area in the same conditions as panel d after 14 days in culture. Data are normalized to control condition (no added E2). **(f)** Quantified ATP of HCI-011 in panel e after 14 days in culture. Data are normalized to the control condition (no E2). **(g)** Immunofluorescence staining of ER (green) and EpCAM (red) on cytospin slides of HCI-011 PDxOs without E2 stimulation. Scale bar indicates 75 um. **(h)** HCI-011 PDxO xenograft (PDxoX) tumor response to 40 mg/kg or 200 mg/kg fulvestrant treatment. Data are shown relative to tumor volume at treatment start (n=4 for vehicle, n=5 for treatment). **(i)** Tumor growth rate calculated from data in panel g. All data are shown as mean ± SEM with technical replicates of n=3 for live cell area and n=4 for qRT-PCR. Statistical tests: (b), (e), ordinary two-way ANOVA, uncorrected Fisher’s LSD with single pooled variance; (d), (h) ordinary two-way ANOVA, uncorrected Fisher’s LSD with individual variances computed from each comparison; (f), two-tailed unpaired t-test; (i), ordinary one-way ANOVA, Tukey’s multiple comparison with single pooled variance. Statistically significant differences are *p < 0.05, **p < 0.01,***p < 0.001, unless indicated otherwise.

We found that this new standard culture condition consistently supported growth of TNBC PDxO lines, but was not optimal for growth of all subtypes of breast cancer. ER+ PDxOs displayed good viability when organoids were freshly grown from PDX tumors, but failed to regrow after dissociation for passaging, a necessary step for expanding PDxO cultures long term. To determine additional supplementation needed for ER+ PDxOs to thrive long term, we first used HCI-011, a pretreated ER+ metastatic breast cancer PDX, as a test case. HCI-011 PDxO was dissociated after 14 days in initial culture in standard medium, and then tested with another set of common organoid medium supplements (**Figure S48a-d**). Y-27632 was determined to be the most effective Rho kinase inhibitor of the three tested, and most supplements did not outperform Y-27632 alone (**Figure S48c, left panel**). In the presence of Y-27632, the addition of N-acetylcysteine (NAC), oleic acid, or basic fibroblast growth factor (bFGF) proved beneficial in promoting growth of HCI-011 PDxO post-dissociation (**Figure S48c-d**). Oleic acid was not pursued further, because we noted that it caused accumulation of lipid-filled vacuoles in PDxO lines (not shown). Thus, our optimal medium for ER+ PDxO comprised our basal conditions plus bFGF and NAC. Again, both our optimized medium and the growth medium described by Sachs et al^20^ supported long term growth, although differences in growth rate were observed with some lines (**Figure S48e**). The significance of *in vitro* growth rates in terms of pathophysiological relevance to human tumors is unclear, so we again chose the simpler, more cost-effective medium and next looked for evidence of ER functionality in growth of the organoids.

ER+ breast cancer causes the most deaths compared to any other subtype, yet there is a paucity of models that faithfully retain ER expression and function. We examined the functionality of ER signaling in ER+ PDxO lines in culture. We noted that with increasing time in PDxO culture the mRNA expression of *ESR1*, as well as its classical transcriptional target *TFF1*^48^, decreased significantly. However, the levels of these genes in organoids remained the same or higher than that observed in the ER+ cell lines MCF7 and T47D grown in 2D or 3D (**Figure 2c**). We next investigated the estrogen-responsiveness of long-term ER+ PDxO cultures. Our culture conditions contain 5% FBS, which contains β estradiol (E2); we tested whether removing estrogen, or supplementing additional saturating levels of E2, would affect the estrogen receptor function in PDxO lines. We cultured ER+ PDxO in our standard ER+ PDxO medium and compared these conditions to standard ER+ PDxO medium containing charcoal-stripped serum to remove E2. Stripped serum reduced mRNA expression of the ER target gene *TFF1* to undetectable levels, and *TFF1* expression was acutely rescued 8 hours after adding back 10 nM E2 (**Figure 2d**). Survival and growth of HCI-011 PDxO was also stimulated by E2 (**Figure 2e-f**). Similar results were seen with other ER+ PDxO lines (**Figures S49a-e)**, and the magnitude of responses was equal to or better than those of the MCF7 cell line in 3D culture (**Figures 2d-e**).

Examination of ER protein levels in standard ER+ PDxO culture conditions (with the only E2 contributed by serum) confirmed positivity for ER by immunofluorescence (**Figures 2g and S49f**). Three representative ER+ PDxO lines cultured long term under our conditions also demonstrated a dose-dependent response to the selective ER degrader fulvestrant after being engrafted into NSG mice *in vivo* as PDxO xenografts (PDxoX; **Figures 2h-i and S49g-h**). These data indicate that our optimized conditions for long term culture of ER+ PDxO effectively maintain functional ER signaling, and retain estrogen dependence *in vitro* and *in vivo*. Of note, we were able to select for resistance to fulvestrant in the HCI-003 and HCI-011 PDxoX models *in vivo* by allowing responsive tumors to recur after stopping treatment, and then retreating with fulvestrant (**Figures S49i-j and Table 1**). In contrast, we were not able to select for fulvestrant-resistance in the HCI-017 PDxoX model, where tumors did not recur following the first round of fulvestrant treatment (not shown). Together, these models, which demonstrate different degrees of estrogen dependence and response to endocrine therapy, will be useful for future studies to understand mechanisms of endocrine resistance.

Although we have few HER2+ breast cancers in our PDX collection, we have models from metastatic breast cancer patients who had HER2+ primary tumors, but whose tumors lacked HER2 amplification upon progression after HER2-targeted therapy (**Table 1**). These models express variable levels of HER2 in the PDX^37^ and in PDxO culture (**Figure S50a**). We optimized long term PDxO conditions for these lines as well, and found that addition of neuregulin 1 (NRG1) was superior to any other supplement tested; NRG1 significantly boosted the growth and metabolic activity of HCI-005, an example of a PDxO with a HER2+ history (**Figure S50b**).

Even with optimized growth conditions for each subtype of breast cancer, some organoid cultures were difficult to establish due to stromal cell takeover. Certain tumor lines had more aggressive stroma, and in these cases stromal cells were removed using fluorescence-activated cell sorting (FACS) during passaging (see Methods). In summary, we optimized conditions for long term culture of several breast cancer subtypes: TNBC, ER+, and HER2+ PDxO lines, including those originally HER2+, and considered each line to be “established” once it was confirmed to be human breast cancer, free of mouse stroma, and reliably passageable. We next set out to characterize their biology and, most importantly, their ability to represent the tumors from which they were derived.

### PDxO cultures are heterogeneous and retain their phenotypes over long-term culture

Using our optimized conditions, we derived 40 PDxO lines from 47 attempts (85% take rate) using PDX tumor material from the current HCI PDX collection (**Table 1**), with additional lines in progress, including those from Baylor College of Medicine PDX models^45^ (https://pdxportal.research.bcm.edu/). These lines have now been cultured for 200-500 days. PDxO lines from different tumors are diverse with respect to morphology and behavior (**Figure 3a**). For example, despite the fact that both are TNBC, HCI-001 forms cohesive organoids while HCI-002 grows as dispersive cell clusters. Some PDxO lines, like HCI-010 and HCI-019, are morphologically heterogeneous even within the same culture. Generally, PDxOs are solid spheres of cells, with few extruding cells. Doubling times of PDxO cultures are also diverse, ranging between three and eight days, with no notable trends across breast cancer subtypes (**Figure 3b**), although doubling times below four days were only observed in TNBC PDxO lines. PDxOs are also diverse in size and density (**Figure 3a**).

**Figure 3.**
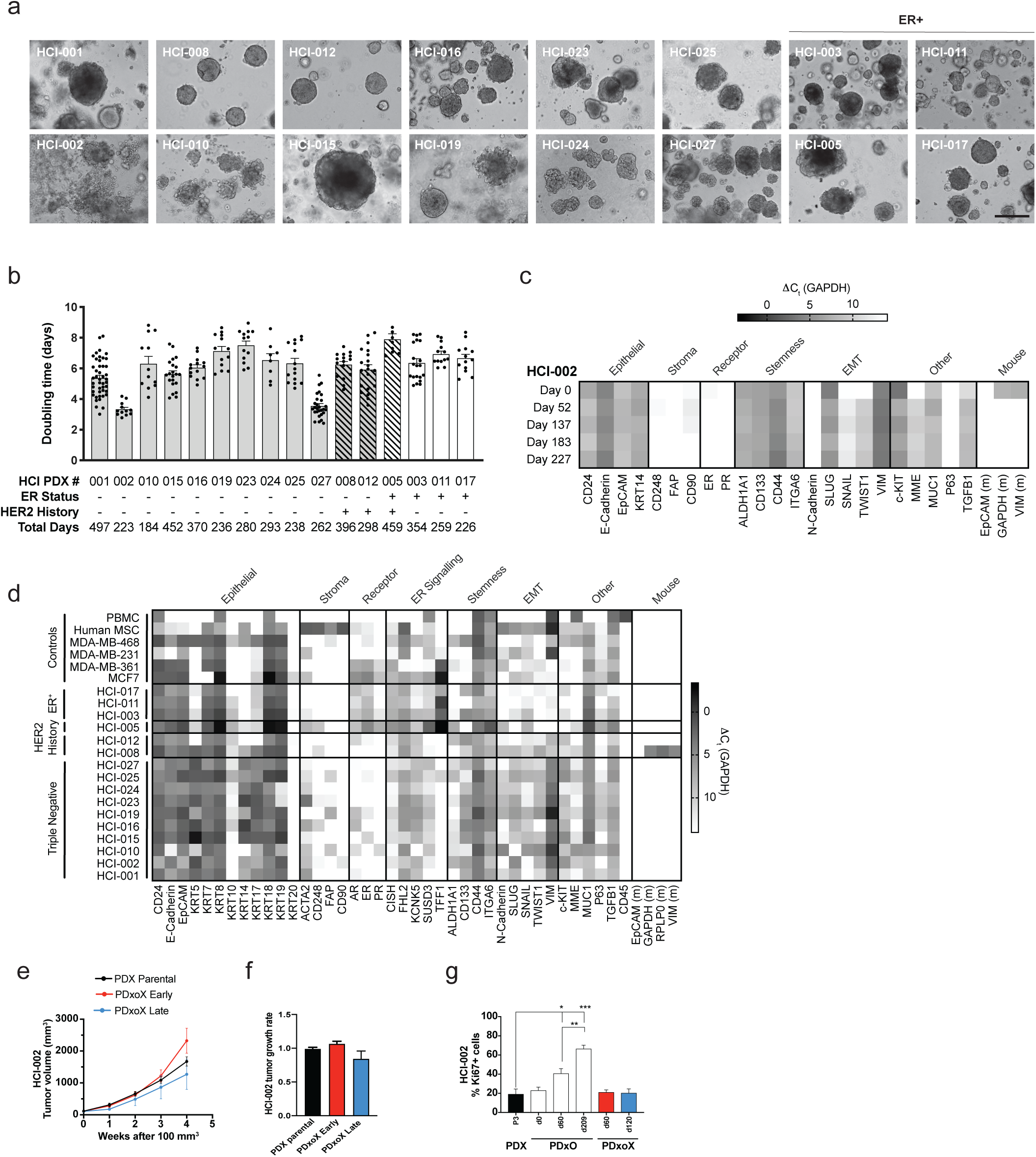
Characterization of established PDxOs. **(a)** Brightfield images of representative PDxO lines. Scale bar indicates 50 um. **(b)** Culture doubling times for each established PDxO line over long-term culture. Each dot indicates a culture passage. **(c)** Expression of PDxO characterization genes on HCI-002 PDxO at different culture time points. Data are normalized to human GAPDH Ct and represent technical replicates of n=4. **(d)** Expression of gene expression panel at single timepoints for representative PDxOs validating their human epithelial nature and subtype status, with additional markers to highlight diversity across different lines. Data are normalized to human GAPDH Ct, and technical replicates of n=4. **(e)** HCI-002 PDxoX tumor growth and **(f)** tumor growth rates compared to the parental HCI-002 PDX line (n=5 mice per group). PDxOs were in culture for 52 days (PDxoX early, n=5) or 121 days (PDxoX late, n=4) prior to xenografting. **(g)** Quantification of Ki67+ nuclei in parental PDX tumors, early (56 days) and late (125 days) PDxO cultures, and early and late PDxoX tumors. Data are averaged from 3 individual sets of IHC stains, and normalized to total hematoxylin^+^nuclei count for each image. All data are shown as mean ± SEM. Statistical tests: (f) ns by ordinary one-way ANOVA, Tukey’s multiple comparison with single pooled variance; (g) ordinary one-way ANOVA, Tukey’s multiple comparison with single pooled variance Statistically significant differences for Ki67 quantification are indicated by *p < 0.05, **p < 0.01,***p < 0.001.

All PDxO were validated to be human epithelial cells, and were diverse with respect to the expression of other selected genes by qPCR (**Figure 3c-d**). For each individual PDxO line, gene expression across selected panels was quite consistent over time in culture (**Figures 3c and S51**). Exceptions include changes noted concurrently with the loss of PDX mouse stromal cells over time, assessed with mouse-specific qRT-PCR assays for GAPDH, vimentin, and EpCAM (**Figures 3c-d and S51**) and the gradual loss of E-Cadherin in HCI-010 (**Figure S51**). HCI-008, an inflammatory breast cancer which was originally HER2+, was also notable in its recruitment and retention of stromal cells (**Figure 3d**), which eventually had to be removed by FACS sorting.

We characterized several PDxO lines in depth, and asked whether they retained their original tumor characteristics after being propagated long term as organoids. When re-implanted into mice as xenografts (PDxoX), tumor growth rates were generally not statistically different from parental PDX tumor growth rates, even when implanted after different time points in culture (**Figures 3e-f and S52**). Exceptions were HCI-001, which showed a consistent decline in PDxoX tumor growth rate with increasing time in culture (**Figure S52a**), and HCI-010, where PDxoX derived from an early culture time point grew significantly faster than the parental PDX counterpart (**Figure S52b**). Interestingly, the late culture time point of HCI-010 failed to grow PDxoX at all, coincident with its phenotypic switch to loss of E-cadherin and gain of Slug expression (**Figure S51**). The same result was obtained with an independently-generated PDxO line from a different HCI-010 PDX tumor (not shown), suggesting this phenomenon is intrinsic to the biology of this model.

Although some PDxO cultures had more Ki67-positive cells than what was observed in tumors growing in mice, Ki67 staining showed that the percent of proliferating cells was not different between PDxoX and PDX, across five PDxO lines examined (**Figures 3g and S52**). Ki67 staining also revealed that for PDX tumors with <20% of Ki67-positive cells, PDxO culture prompted a significant increase in proliferation; however, for PDX tumors with >50% Ki67+ cells, Ki67+ levels were not significantly altered in culture (**Figures 3g and S52**). Organoid morphology was assessed by hematoxylin and eosin (H&E) staining and immunohistochemistry (IHC) with antibodies specific for human specific-vimentin, Ki67, and human-specific cytokeratin 8 (CAM5.2). PDxO cultures again resembled their originating PDXs, and PDxoXs generated after different lengths of time in culture also resembled their PDX counterparts (**Figures S53-56**).

### PDX and PDxO retain genomic characteristics of their originating human tumors

Breast tumors have specific patterns of DNA methylation, which distinguish cancers from normal tissue or benign tumors, and reflect breast cancer subtype and biological features such as metastatic potential^49–50^. As an initial comparison between our models and the original tumors from which they were derived, we performed genome-wide DNA methylation analysis on nine sets of matched patient tumors, PDX and PDxO models, and an additional two PDX-PDxO pairs for which primary tumor material was not available. These data revealed that the patient-derived models are more similar to their originating human tumors than are commonly used cell lines (**Figure 4a**).

**Figure 4.**
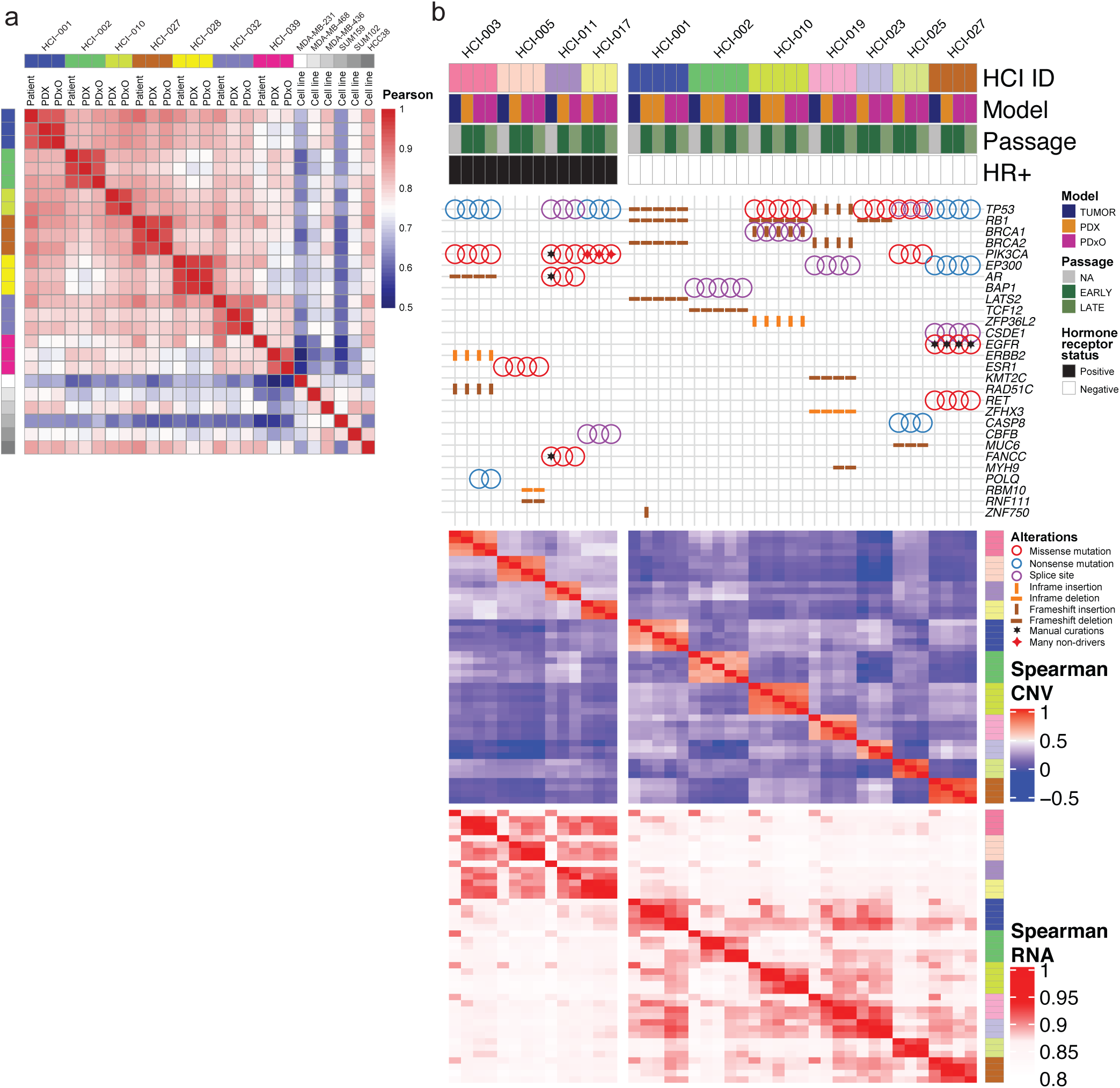
Genomic landscape of PDxOs compared to PDXs and patient tumors. **(a)** Correlation heat map illustrating genome-wide DNA methylation analysis for seven sets of patient-derived models, compared to commonly used breast cancer cell lines. Color scale indicates Pearson Correlation Coefficient **(b)** Eleven sets of models were characterized at different time points (early and late) to assess molecular fidelity with the patient tumors. The heatmap is divided into four sections from top to bottom: annotations, exome sequencing variant detection, copy number correlations from SNParray data, and RNA-seq gene expression correlations. Mutation variants are shown with an oncoprint plot highlighting single nucleotide variants and indels for significantly mutated genes in breast cancer. Quantitative CNV correlations are shown using a heatmap of Spearman correlations for gene-level log2 CN ratios. Quantitative transcriptome correlations are shown using a heatmap of Spearman correlations for gene-level log10 transformed RSEM count estimates.

We next performed in-depth genomics characterization to assess the molecular fidelity of a subset of our patient-derived tumor models over time. We chose models for which we had both early- and late-passage matched PDX and PDxO models (11 sets of models comprising both ER+ and TNBC), and conducted in-depth genomic and transcriptomic analysis. We systematically assessed mutations, copy number variation, and transcript abundance correlations between samples from patient tumors (when available), PDX, and PDxO (**Figure 4b**). Single nucleotide variants and small indels were predicted using the PDXNet somatic mutation calling pipeline available on Seven Bridges (see Methods). In-depth characterization showed high genomic concordance between patient samples, PDX and PDxO models. The patient-derived models maintained genetic mutations in known cancer genes when comparing PDX to PDxO and patient tumor material (**Figure 4b, top panel**). While common driver mutations (e.g. those in *TP53*, *RB1*, *BRCA1*, *PIK3CA* genes) were retained in all early- and late-passage models, unique variations in five cancer-associated genes were observed in four of the 11 sets of models. The model-specific variations include a *POLQ* nonsense mutation only in PDxO derived from HCI-003, and *RBM10* and *RNF111* deletions only in PDxO derived from HCI-005. We also found a *ZNF750* in-frame insertion in early-passage HCI-001 PDX, but not in later-passage PDX nor early- or late-passage PDxO; and a frameshift deletion of *MYH9* that was observed in HCI-019 PDxOs, but not in the PDX or patient tumor. In all cases, the organoid lines were made from a different passage of the PDX than that which was sequenced. Thus, mutational differences could be a result of subclone selection bias from passaging heterogeneous tumors, or could be a result of a small degree of tumor evolution. However, these data show that, overall, the major cancer mutational drivers observed in patient tumors were largely maintained in models at various passages.

Systematic, engraftment-specific copy number changes have been reported to occur in PDX models^51^; however, a recent comprehensive study found that copy number changes are minimal and largely attributed to spatial heterogeneity of samples, rather than evolution upon PDX engraftment^1^. To examine whether gross copy number changes occurred in our models following *in vivo* and *ex vivo* passaging, we assessed copy number differences in the early- and late-passage models versus their matching patient tumors using SNP arrays. Overall, we observed a high correlation between patient samples and their derived models compared to models from different patients (p=9.14e-42, 95% CI = 0.57 to 0.63, Mann-Whitney U of Spearman correlation values; **Figure 4b** and **Figure S57**). This result supports the recent PDXNet finding of gross CNV conservation from patient samples to their derived models^1^. At the gene level, we only observed a few genes that transitioned from amplification to deletion following model passaging. One such example is *SNCG* (10q23.2) in HCI-001. *SNCG* encodes synuclein gamma, which is a protein shown to affect the mitotic checkpoint function of breast cancer cells and is associated with a higher percentage of invasive and metastatic breast carcinomas^52^. Interestingly, *SNCG* is the only gene of 30 in the region that transitions from amplified to deleted across passages (CN ratios = 0.99, −0.56, −0.66, −1.47, −1.51 for patient tumor, PDX-early, PDX-late, PDxO-early, PDxO-late, respectively). Another five genes exhibiting large genomic changes (*PSMB1*, *PDCD2*, *TBP*, *OR4F7P*, *WBP1LP8*) were found in HCI-010 as the region transitioned from amplified in the patient tumor to deleted in the early passage of the PDX and PDxO (**Figure S57**). Thus, while some copy number alterations may exist at specific segments between models and may reflect some selection in the PDX, our data suggest that xenograft and organoid models generally retain the genomic structure of the original patient tumor.

In addition to sequence variant and CNV analysis, we sought to correlate specific transcript abundance across multiple time points in these same models. RNA-seq reads were processed using the PDXNet workflows on Seven Bridges (see Methods). Analysis of the whole transcriptome showed a high correlation of transcript abundance between samples (0.86 SD +/− 0.023, Spearman correlation coefficient). We observed that intra-model correlation was significantly greater than inter-model correlations for each model (p=7.75e-29, Mann-Whitney U of Spearman correlation values; **Figure 4b**). We noted less similarity between patient tumors and models in ER+ cases, suggesting that more selective pressure may be applied to the ER+ models (p=0.0008, 95% CI −0.03 to −0.01, Mann-Whitney U of Spearman correlation values). This is consistent with the lower “take rates” of ER+ PDX compared to TNBC. Together, our data confirm the genomic heterogeneity of patient samples, and provide evidence that intra-model derivatives of the patient tumor are collectively more similar than different models are to each other.

### PDxO models are amenable to drug screening and results are concordant with drug responses *in vivo*

Following optimization of PDxO long term culture conditions and validation of phenotypic and genomic fidelity to their originating tumors, we developed drug testing and screening protocols for PDxOs. PDxO cultures were dissociated and plated into 384-well format for drug testing in a four-day drug response assay. We used a large dose range of each compound, so that the entire range of effects (growth, cytostasis, or cytotoxicity) could be observed in each PDxO line with each drug in a relatively high-throughput manner. Sixteen PDxO lines were screened against a panel of 50 compounds in an eight-point dose response assay, with technical quadruplicates, and in biological triplicate, to determine therapeutic response. Due to variable doubling times between models, we used the growth rate correcting strategy proposed by Hafner et al. to calculate and score our dose-response curves^53^. This method adjusts cell viability using a log2 fold change estimator and allows us to compare drug responses between models from different patients even when growth rates were different. We found that GR50, the concentration at which we model half-maximal adjusted growth, and GRaoc, the area over the dose-response curve (see Methods), are suitable estimations for organoid drug sensitivity and cytotoxicity. The landscape of treatment responses across different PDxO lines and different drugs is shown in **Figure 5a** and **Figure S58**.

**Figure 5.**
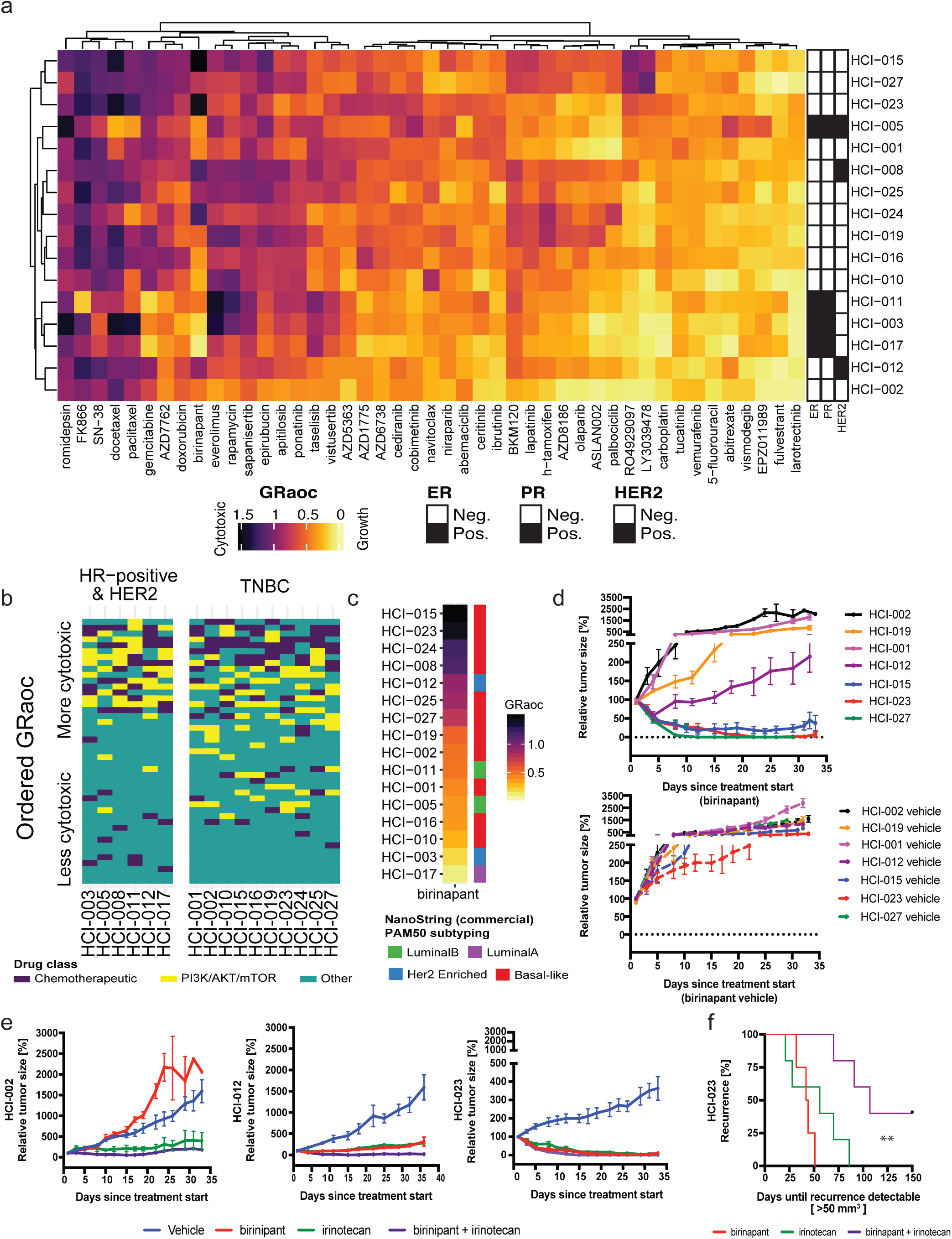
PDxO drug screening shows concordance with *in vivo* data and identifies birinapant as a potential therapy for some TNBC tumors. **(a)** Unsupervised clustering of 16 PDxO models and 45 screened compounds. Color indicates GRaoc statistics (darker colors indicate cytotoxicity; lighter colors indicate growth). Annotations for each model indicate hormone status for ER, PR, and HER2. **(b)** A tile plot displays sample-specific drug ranks colored by drug class: chemotherapeutic agents (dark purple), PI3K/AKT/mTOR targeted agents (yellow), and all other drugs (teal). Samples are separated by hormone receptor (HR)-positive and HER2+ tumors or TNBC. **(c)** Stacked heatmap rank of PDxO models for birinapant drug responses. Samples are sorted by GRaoc, with best responses on top. The PAM50 breast cancer subtype for each model are displayed to the right. (d) *In vivo* drug treatment response to birinapant in various PDX models (top) with matching vehicle controls (bottom). (e) *In vivo* drug treatment response to birinapant, irinotecan, or combination in HCI-002 (left), HCI-012 (middle) and HCI-023 PDX models (right). **(f)** Time to recurrence of HCI-023 PDX tumors following cessation of birinapant, irinotecan, or combination treatment (**p < 0.0046, Log-rank Mantel-Cox test).

As expected, certain drugs showed selective effects on particular breast cancer subtypes. For example, cytotoxic chemotherapies showed more activity against TNBC lines compared to non-TNBC lines (p=6.42e-3, Mann-Whitney U, comparison of GRaoc scores, 95% CI 0.05 to 0.29), while PI3K, AKT, and mTOR inhibitors showed more activity in ER+ and/or HER2+ lines compared to TNBC lines (p=9.56e-5, two-sided Mann-Whitney U comparison of ranks, 95% CI 3 to 9 rank positions; **Figure 5b**). Importantly, PDxO drug responses were reproducible when the screens were repeated multiple times over up to one year in culture (**Figure S59a-b**), and similar responses were achieved with compounds made by different companies against the same target (**Figure S59c**). These data indicate that breast cancer PDxOs can be efficiently screened for drug responses, and that the responses are reproducible over time in culture and across multiple replicate assays. Using GRaoc scores (**Figure S60a**), each model can be ranked for sensitivity to each drug. Examples are shown for the relative sensitivity of HCI-010, a TNBC model, to the BCL2 antagonist navitoclax, where HCI-010 was the most responsive PDxO line (**Figure S60b-c**). Likewise, HCI-010 was also the most responsive PDX when navitoclax was tested in a panel of five different TNBC PDX models *in vivo* (**Figure S60d-e**). Similar results comparing organoid and PDX responses to docetaxel are shown in **Figure S61**, also with good concordance.

To investigate further how well drug responses in PDxO mirrored drug responses in PDX, we selected drugs that showed very distinctive response or resistance patterns in PDxO lines. For example, six of the twelve TNBC PDxO lines responded with remarkable sensitivity to birinapant, a SMAC mimetic and inhibitor of apoptosis protein antagonist^54^, while the remaining six TNBC lines were resistant to high doses of birinapant (**Figure 5c**). We tested birinapant *in vivo* on seven PDX lines that were predicted to span a range of birinapant sensitivity according to PDxO results. TNBC PDXs predicted to be resistant to birinapant (HCI-001, HCI-002, and HCI-019) resulted in progressive disease similar to controls, whereas TBNC PDX lines predicted to be sensitive (HCI-015, HCI-023, and HCI-027) resulted in tumor shrinkage with birinapant treatment (**Figure 5d**). HCI-012, a HER2+ model, had an intermediate response, showing initial shrinkage followed by growth (**Figure 5d**).

As another example of concordant drug responses, we unexpectedly found that the TNBC PDxO lines HCI-015 and HCI-027 showed exceptional responses to two different gamma-secretase inhibitors, RO4929097 and LY3039478 (**Figure 5a, Figure S59c**). We were unable to obtain LY3039478 for *in vivo* studies, but found that testing of RO492097 in PDX was concordant with PDxO results for the two responsive and two unresponsive lines (**Figure S62**). Interestingly, we observed a *NOTCH1* copy number amplification (**Figure 1**) that may explain the favorable response of HCI-027 to RO4929097.

We next investigated whether drug combinations could be efficiently tested in PDxO culture using synergy matrices. From the literature, we identified a potential synergistic interaction between birinapant and SN-38 (the active metabolite of the prodrug irinotecan) in ovarian cancer cells *in vitro*^55^. Using dose responses of both drugs separately and in combination, we determined that this synergy was also apparent in breast cancer PDxOs, especially for birinapant-sensitive tumors like HCI-023 (**Figure S63**). Similar to the PDxO synergy results, *in vivo* treatment of PDX HCI-002 (birinapant resistant) showed little or no benefit to the irinotecan combination versus irinotecan alone, while the combination treatment improved response in the partially birinapant sensitive line HCI-012 (**Figure 5e**). In the birinapant-sensitive line HCI-023, either birinapant or irinotecan was able to completely eliminate tumors, but the combination treatment resulted in a more durable response following treatment cessation compared to either of the single agents (**Figure 5e-f**). Together, these results suggest that PDxOs are able to predict drug responses in PDX models *in vivo* accurately, and illustrate the potential power of obtaining functional drug response data to identify new, effective treatment options for breast cancer.

### PDxO drug screening is feasible to inform patient care for aggressive breast cancer

To determine if establishment of PDxO cultures and subsequent drug screening can be feasibly performed to inform patient care, we present the case of a 43 year-old woman with mammographically-detected stage IIA TNBC (**Figure 6a**). She was enrolled in our TOWARDS study, for which a treatment-naïve biopsy was taken to make PDX. She then received preoperative chemotherapy with doxorubicin and cyclophosphamide given every two weeks for four cycles, followed by weekly paclitaxel for 12 weeks (AC-T therapy, as per standard of care). Surgical pathology (lumpectomy and sentinel lymph node biopsy) was consistent with complete pathologic remission (pCR), and adjuvant radiation therapy per standard of care ensued. However, rapid growth of the PDX (HCI-043) in the interim suggested a high risk of recurrence based on our published data^37^.

**Figure 6.**
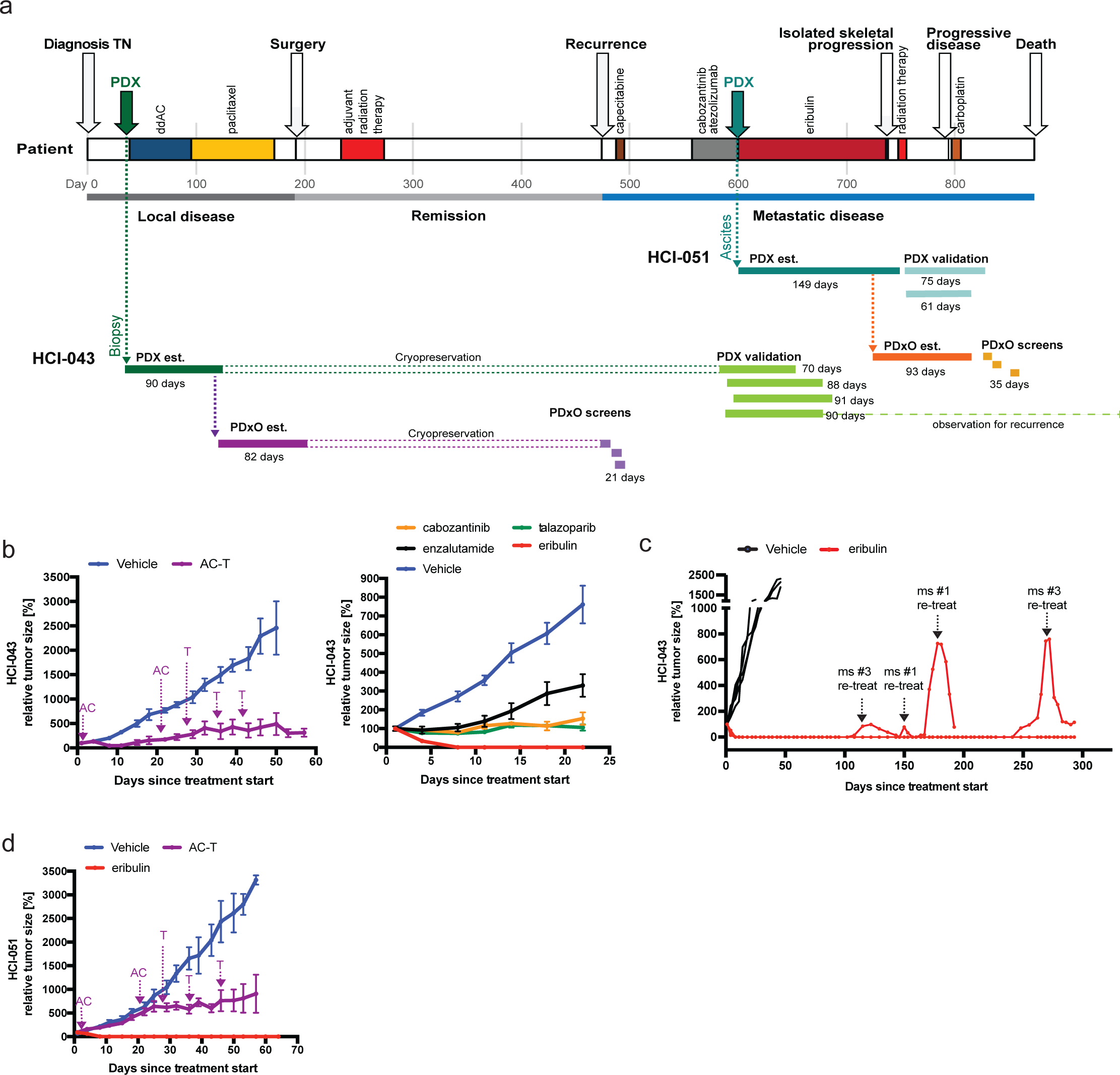
PDxO screening can be performed in real-time with patient care. **(a)** Timeline of the patient HCI-043 including clinical history, patient-derived model establishment (PDX and PDxO), PDxO drug screens, and *in vivo* validation of responses in PDX. Model development and drug screening/testing was done with both the pre-treatment biopsy sample (HCI-043) and the metastatic ascites sample (HCI-051), with similar results. **(b)** Treatment of HCI-043 PDX with patient-matched neoadjuvant therapy (adriamycin+cyclophosphamide followed by paclitaxel, AC-T; left) or with drugs selected from the PDxO screen (right). Arrows indicate the sequencing of AC-T drug treatment. **(c)** Follow up of HCI-043 PDX mice after stopping treatment with eribulin following 3 doses. Three different mice exhibited recurrence off-treatment, but the tumors regressed after treatment was restarted. No resistant tumors were detected over the lifespan of the mice (293 days after initial treatment began). **(d)** Treatment of HCI-051 PDX with AC-T, as in panel b. As expected, the metastatic sample was more resistant to this neoadjuvant therapy.

Despite the achievement of pCR, which portends a favorable prognosis^56, 57^, less than seven months after completion of her treatments the patient experienced an early metastatic recurrence in the liver (**Figure 6a** and **Figure S64a**). Her first line of therapy in the metastatic setting (to which we were blinded) consisted of capecitabine, but the disease did not respond. She experienced progression with new onset skeletal metastases (**Figure 6a**). Her tumor tested weakly positive for PD-L1 (**Figure S64b**), which can portend a favorable response to atezolizumab^58, 59^, so her next line of therapy consisted of cabozantinib and atezolizumab in the context of a clinical trial. Cycle two of this regimen was delayed due to adverse events, but eventually the investigational therapy was stopped due to progressive disease (**Figure 6a**).

In the meantime, we had already developed the HCI-043 PDxO line corresponding to this patient’s untreated primary tumor, and had cryopreserved stocks. Upon learning of her initial metastatic recurrence, we thawed the organoids and screened a library of FDA-approved and experimental drugs. We also performed genomic profiling, consisting of whole genome sequencing and bulk- and single cell-RNA sequencing on the PDX model. Commercial genomics analysis did not reveal any clinically-actionable mutations, but did show that the tumor belonged to the luminal androgen receptor (LAR) breast cancer subtype^60^, with AR mRNA being expressed in all tumor clones and AR protein detected in a subset of cells by clinical immunohistochemistry staining (**Figure S64c**). In the organoid drug screen, two FDA-approved breast cancer drugs, eribulin and talazoparib, emerged as promising candidates, while several of the chosen clinical therapies, including 5-fluorouracil (used in our organoid screen as the active metabolite of capecitabine) and cabozantinib, did not appear to be effective in the organoid assays (**Figure S64d**). We also noted genomic deletions in *BRCA1*, *BRCA2*, and *RAD50* (**Figure 1**), which may explain the enhanced sensitivity to talazoparib.

*In vivo* testing on the HCI-043 PDX model verified a non-complete response to the patient-matched AC-T therapy, and confirmed PDxO data that eribulin was the best drug tested (**Figure 6b**). Enzalutamide (tested as an androgen receptor antagonist due to the LAR phenotype), cabozantinib, and talazoparib each reduced the growth rate of the tumor compared to vehicle-treated controls, but did not cause tumor regression. The durability of the apparent complete response to eribulin was tested in mice by stopping treatment after three doses and following the animals for tumor recurrence. Tumors that recurred off treatment were then tested for resistance by restarting treatment. In all mice, there was either no recurrence or complete regression on retreatment over the course of 293 days, when we had to euthanize the mice for old age (**Figure 6c**).

With IRB approval, we returned our drug testing results to the clinic. The patient was put on eribulin, and the hepatic metastases sustained a complete remission for a period of nearly five months (**Figure 6a and S64a**). In the meantime, we also generated PDX and PDxO (HCI-051) from an ascites sample from the patient, collected just prior to starting eribulin treatment (**Figure 6a**), representing the AC-T resistant metastatic recurrence. The metastatic models were indeed more resistant to AC-T; however, they were still completely responsive to eribulin (**Figure 6d and S64e**).

While the patient’s liver metastases and ascites regressed completely in response to eribulin (**Figure S64a**), the patient eventually had an isolated progression in bone, for which she received radiation therapy; eribulin was withheld. While off systemic therapy, her liver metastases remained in remission for two additional months but eventually returned. It is unknown whether the recurrent liver metastases were sensitive to eribulin because a different systemic therapy was given at that point (carboplatin; **Figure 6a**). Carboplatin was not effective and, unfortunately, the patient died. In an analysis of all therapies given, the progression-free survival (PFS) and time to next systemic therapy (TTNT) that was achieved with the PDxO-informed therapy (eribulin) was 3.5 and 4.8 times longer compared to the PFS and TTNT achieved with the prior therapy (138 days vs. 41 days, and 197 days vs. 41 days, respectively). Of note, a recent clinical trial using genomically-informed therapies in patients with metastatic cancer (MOSCATO-01) used a matched therapy to prior therapy PFS ratio of only 1.3 as a benchmark of success^61^. These results suggest that functional drug testing with PDxOs is feasible and can give beneficial results in real-time with patient care.

## Discussion

We report development and characterization of a large bank of >100 patient-derived xenografts and matched long-term organoid cultures. Although a large bank of human breast tumor organoids has been previously described by Sachs et al^20^, it is important to note how our collection is different. First, the organoid collection described by Sachs et al was primarily developed from primary, untreated breast tumors. While primary breast tumors are easy to obtain during surgery, these tumors are curable 70-80% of the time with standard therapy. The greatest unmet medical needs in breast cancer are in treatment-resistant and recurrent/metastatic tumors. 65% of our collection comes from treatment-resistant tumors and represents recurrences from 8 different metastatic sites. We also made a concerted effort to develop models of primary-metastatic pairs or longitudinal collections over time from the same patients, and each model is accompanied by comprehensive, de-identified clinical data. Second, our collection of matched PDX and PDxO models facilitates high-throughput analysis and drug screening on models of advanced breast cancer representing the population of patients likely to be in clinical trials. We find that PDxO-based drug screening is feasible and cost-effective, and allows *in vivo* validation of results in the matching PDX models. Our data revealed consistency between drug screening results in organoids and drug responses in PDX. Moreover, we demonstrated the feasibility of using these patient-derived models for precision oncology. In a case study of a triple negative breast cancer with early metastatic recurrence, we found that drug responses in the models, to which the treating oncologist was initially blinded, aligned with observed clinical responses. Although our study was not originally designed to prospectively inform patient care, compelling results uncovered a potentially effective, FDA-approved drug and led to IRB approval to return those results to the clinic. Treatment with the PDxO-directed therapy resulted in a complete response for the patient and a PFS and TTNT periods that were 3.5 and 4.8 times longer, respectively, than her previous therapies. These data suggest that this new PDxO screening methodology can be leveraged in the clinical arena to identify therapeutic vulnerabilities to FDA-approved or investigational agents to inform medical decision making. Future trials will be required to determine if this strategy of functional precision oncology using patient-derived models can improve outcomes in patients with metastatic breast cancer.

Our data also showed that PDxO screening can uncover experimental drugs that have therapeutic potential. We found that birinapant, a SMAC mimetic, showed potent activity in certain TNBC organoid models, and this was validated in PDX models. Indeed, others have recently shown that birinapant has selective activity for TNBC as compared to ER+ breast cancers^62, 63^, and a similar SMAC mimetic (LCL161) combined with paclitaxel showed clinical activity in the neoadjuvant setting, but only in TNBCs expressing a TNFalpha signature^64^. These data demonstrate that screening our human breast cancer PDxO collection can identify new experimental agents that can have clinical activity in particular breast cancer subtypes.

PDxO drug response assays are not without limitations, however. Although we were able to discern cytotoxic effects in our assays, we were unable to reliably detect activity of drugs that convey cytostatic or less potent activity. For example, response of several ER+ PDxoX lines to fulvestrant is shown in **Figure 2** and **Figure S49**; however, response to fulvestrant was difficult to discern in PDxO models (**Figure 5**). Similarly, 4-hydroxytamoxifen responses in PDxO did not correlate with ER status (**Figure 5**). It is likely that four days of drug exposure on models that are relatively slow growing, such as ER+ breast cancer, is not enough to reveal susceptibilities, especially when the main biological outcome is to maintain stable disease (as for fulvestrant). Thus, four-day PDxO drug responses are best for identifying drugs with cytotoxic activity. Future work will determine whether longer-term drug exposure, possibly with passaging, will be a better read-out for less potent, yet clinically relevant, drug activity.

Human cancers are naturally heterogeneous, and heterogeneity of tumors is also considered to be a barrier to successful therapy. Models of human cancer that retain tumor heterogeneity are important to assess treatment efficacy and to investigate inter-clonal interactions that facilitate treatment response or resistance. Heterogeneity of PDX models has been investigated extensively and, although some “bottlenecking” can occur especially in the first passage^65^, PDXs can effectively recapitulate the heterogeneity of human cancer (recently reviewed^2^). Prostate cancer PDX-derived organoids have been shown to retain the genomic features of PDX models, at least in the short term^15^, although this has not yet been thoroughly investigated for PDxO models of breast cancer. Our data indicate that critical cancer driver alterations and copy number changes are largely maintained throughout the derivatives of patient tumor modeling from the acquired patient sample to PDX to PDxO. Transcript abundance is also highly correlated within patient sample derivatives.

It is important to consider inter-tumor and intra-tumor heterogeneity when models derive from a particular tumor site(s), however. In the case of HCI-043/HCI-051, models were made from the primary untreated breast tumor and from ascites following recurrence in the liver. Both models showed strong response to eribulin and, while the patient’s liver metastases regressed completely and did not return while on eribulin, a metastasis in the bone appeared during this treatment. It is unknown whether the eribulin-resistant bone metastasis in the patient was a different tumor clone that was perhaps not represented in our models. Attempts to make models from as many anatomical locations as possible should help inform these tests. Models derived from circulating tumor cells (CTCs) may better represent the heterogeneity of the disease in the patient^66^, but it is difficult to derive models from CTCs for certain cancers, including breast cancer, due to low CTC numbers. Integration of functional data from patient-derived models such as PDX and organoids with genomic data from circulating tumor DNA (ctDNA) may allow prioritization of drugs that might be effective in multiple tumors.

One limitation of many models of human cancer, whether *in vitro* or *in vivo*, is lack of human stroma, including human immune cells. Human stromal cells are replaced by mouse stroma during PDX growth in immunodeficient mice^37, 67^. Human immune cells can be engrafted to allow “humanized” immune systems in mice with PDX^68, 69^, but determining the full functionality of the engrafted immune system is plagued by variability of immune engraftment, and the recapitulation of tumor-immune interactions have not been well established in this reconstituted context. In our PDxO system, the mouse stroma is purposefully removed during organoid propagation because some tumors recruit an aggressive stroma that compromises the ability of the organoids to thrive. As a result, the organoid platform, like most other *in vitro* systems, does not fully recapitulate the natural tumor microenvironment including immune cells, which limits the classes of drugs that can be tested in this system.

In summary, this work provides a large, clinically-relevant resource of paired *in vivo* and *in vitro* patient-derived models of breast cancer, with an emphasis on the most difficult cases for which research advances are urgently needed. We show that these models can be used for drug screening and discovery, and our methods are also conducive to conducting functional precision medicine in real time with patient care.

## Methods

### Development of breast cancer PDX lines

Human samples were collected and studied under approved IRB protocols #89989 and #10924 at Huntsman Cancer Institute, University of Utah. All procedures using live animals were reviewed and approved by the University of Utah Institutional Animal Care and Use Committee (IACUC). Tissues were immediately processed under sterile conditions. The detailed tissue processing and implantation protocol has been previously described^47^. Briefly, female immune-compromised mice (strains NSG (Jax stock #5557), NOD/SCID (Jax stock #1303), or NRG (Jax stock #7799) were used to generate PDX, typically at the age of 3-4 weeks. In rare cases, for urgent situations with younger mice not available, we used mice up to 10 weeks of age. Fresh or thawed human breast tumor fragments were implanted into the cleared inguinal mammary fat pad. In some cases (bone metastasis samples), bone fragments were co-implanted. For liquid specimens such as pleural effusion or ascites fluid, samples were processed as previously described^47^ and 1-2 million cells were injected into cleared mammary fat pads in 10-20 microliters of commercial Matrigel (BD Biosciences). For ER+ tumors, mice were given supplemental estradiol (see below). The mice were monitored for health and tumors were measured weekly with digital calipers once growth was observed. When tumors reached 1-2cm in diameter, tumors were aseptically harvested and re-implanted into new mice or viably banked^47^. The initial tumor was termed “passage 0” (P0), and passages continued to be tracked with each generation. Clinical information and patient demographics for PDX lines combined can be found in the BCM PDX Portal (https://pdxportal.research.bcm.edu/) and on the Welm lab research website (https://uofuhealth.utah.edu/huntsman/labs/welm-labs/research.php).

### Pathogen testing and removal of LDEV or *C. bovis*

PDX tumors were tested for select human pathogens (EBV, HCMV, Hep B, Hep C, HIV1, HIV2, and LCMV) as well as confirming *C. bovis* and LDEV negativity using commercial testing from IDEXX Laboratories (Westbrook, ME). Other common mouse pathogens were monitored using standard sentinel testing in the mouse facility. The facility is a specific pathogen free (SPF facility) and no positive results have been obtained during the course of developing these PDX.

### Fluorescence-activated cell sorting or rat passaging to remove LDEV

A common contaminant in PDX tissue is the single-stranded RNA virus LDEV. LDEV is a mouse-specific pathogen that propagates in tissue macrophages and can cause debilitating infections and even paralysis in immune-compromised host mice. LDEV can be transmitted through serial transplantation of infected tissue in mice or by use of LDEV-contaminated matrigel-like products^70, 71^. Some of our early PDX tumors (HCI-010, HCI-013, and HCI-013EI) were generated with Englebreth-Holm-Swarm (EHS)-derived matrix^47^ prior to the realization that EHS tumors, and some commercial Matrigel lots, were infected with LDEV. To decontaminate LDEV-positive tumors, including EHS tumors, we utilized various methods. In two of the three infected lines, we found it was sufficient to perform FACS sorting using positive selection with anti-human CD298 as a universal cell surface marker of human cells^72^ and negative selection for mouse-specific immune marker CD45, followed by re-transplantation into mice. For HCI-010, this method was not sufficient to remove infected cells even after adding additional negative selection with antibodies specific for mouse CD11b and F4/80, so we passaged HCI-010 once through immune-deficient rats (NIH-RNU; Charles River stock #568) to achieve LDEV-negative status. For FACS sorting, freshly harvested PDX or EHS tumors were processed to single cells^47^ and stained with PE-conjugated anti-human CD298 PE (BioLegend; 1:50 dilution) and FITC-conjugated anti-mouse CD45 FITC (BioLegend; 1:200 dilution). Stained cells were sorted with the BD FACS Aria. Human CD298-positive and mouse CD45-negative tumor cells were collected and washed in HBSS (HyClone), followed by resuspension in LDEV-free matrigel (from Corning or prepared in the lab; see below). 0.5 to 2 million cells in 10-20uL matrigel suspension were injected into the cleared mammary fat pads of 3-4 week-old NSG mice and the tumors were harvested when they reached 2 cm in diameter. Harvested tumors were pathogen tested for LDEV using IDEXX commercial testing. LDEV-free matrigel was prepared by growing EHS tumors subcutaneously in C57BL/6J mice (Jax stock #664), harvesting, and preparing single cell suspensions as previously described^47^. Cells were FACS sorted to remove macrophages as described above. After LDEV was confirmed to be negative, tumor stocks and matrigel were prepared as previously described^47^.

### C. bovis removal

During this study, many institutions began reporting problems with *C. bovis* infection of PDX models due to rapid transmission in immune-deficient mice. Immune-deficient mice infected with *C. bovis* typically show signs of skin problems: red ears, rash and alopecia around the face and the dorsal portion of the neck. Since *C. bovis* infects the skin, a handler or any fomites can transmit the disease very easily among the colony. We screen for *C. bovis* by performing fur swabs on the mice (around face and dorsal neck) and sending the swabs to IDEXX for specific testing. If a swab comes back positive, the mouse is immediately euthanized and disinfected with 2% Chlorhexidine gluconate solution by submerging the entire mouse for 5 minutes^73^. The PDX tumor is then aseptically harvested without allowing skin or fur to touch the tumor. PDX tumors were always confirmed to be *C. bovis* negative on the next passage.

### Estrogen delivery

We modified our previously published estrogen pellet protocol^47^ by reducing the dose to 0.4 mg E2 in the beeswax pellets. E2 was also delivered in drinking water using a protocol kindly shared online by the Wicha lab (http://www.med.umich.edu/wicha-lab). Briefly, a 2.7 mg/mL stock of 17-estradiol (Sigma # E2758) in ethanol was diluted to a final concentration of 8 ug/mL in sterile drinking water. Water was changed once per week, since we found no significant difference in plasma E2 concentrations when the water was changed once vs. twice per week (not shown). Since E2 is reported to be light-sensitive, we also tested the stability of the E2 in the water over time in clear or amber bottles, but found no significant differences in the mouse plasma levels (not shown), so we use clear bottles. In all standard experiments (all those except when growing estrogen-independent sublines), ER+ tumors were grown in mice with a 0.4mg E2 pellet implanted subcutaneously at the time of tumor implantation, followed by administration of E2 in the drinking water beginning four weeks after tumor implantation, until the tumor was harvested.

### Development of estrogen-independent ER+ breast PDX models

ER+ PDX tumors were harvested and transplanted into ovariectomized mice with no E2 supplementation. Tumor sublines that grew under these conditions were considered to be estrogen independent and were given the designation “-EI.” To generate “-EI” lines, 6-8 week-old mice were ovariectomized bilaterally using two separate dorsal incisions, parallel to the midline under general anesthesia using standard procedures (https://www.criver.com/sites/default/files/resource-files/ovariectomy.pdf), immediately followed by tumor implantation into the mammary fat pad. To minimize pain and distress due to the ovariectomy, mice were given buprenorphine (0.1-0.2mg/kg) once before and once after surgery, and carprofen (5mg/kg) daily for three days following surgery. In the case of HCI-013EI, a two-week culture step in phenol red-free HBEC medium^47^ supplemented with charcoal-stripped FBS occurred between the steps of tumor growth in ovariectomized mice and re-transplantation. Future passages of “-EI” lines occurred in intact mice with no E2 supplementation.

### Establishment of PDxO cultures

For PDxO preparation, PDX tissue chunks were digested in a GentleMACS dissociator in PDxO base media (Advanced DMEM, 5% FBS, HEPES, Glutamax, Gentamycin, 10 ng/ml human EGF, 1 ug/ml hydrocortisone) with human tumor dissociation enzymes and 10 uM Y-27632 added. After digestion, differential centrifugation steps were performed to enrich for organoids and deplete single (stromal) cells from the sample. Organoids were embedded in matrigel domes, which were plated in multiwell tissue culture plates onto a matrigel base layer. After a 5 min incubation period, plates were flipped and matrigel domes were allowed to solidify for 10 min before subtype specific culture media was added. For all breast cancer subtypes, 10 uM Y-27632 was added fresh to the culture medium. Additionally, for HER2+ PDxO cultures 10 nM Heregulin β-1 was added, and for ER+ PDxO cultures 100 ng/ml FGF2 and 1 mM NAC (N-Acetyl-L-cysteine) was added. Media was exchanged every 3-4 days and, once mature, cultures were passaged by incubating in 20% FBS in dispase with 10 uM Y-27632, followed by a wash step with base media and a dissociation step in TrypLE Express. Single cells were seeded at 200,000-400,000 cells per 200 ul matrigel dome in a 6 well plate onto a 50 ul matrigel base layer. To eliminate mouse stroma cells from organoid cultures, organoid cultures were either differentially centrifuged several times after dispase incubation, or sorted by FACS. For FACS, single cell suspensions of organoid cultures were incubated with human and mouse anti-FcR, followed by antibody staining (AlexaFluor 647 L mouse CD90.2, AlexaFluor 647 L mouse CD29, AlexaFluor 488 L human CD326, FITC L human CD298). Post sorting, human cells were cultured on ultra-low attachment plates in subtype-specific media overnight and embedded in matrigel domes the next day. Minimal to zero mouse content was confirmed by qRT-PCR. For cryopreservation, mature PDxO domes were frozen in PDxO base media with 20% FBS, 10% DMSO and 10 uM Y-27632.

### PDxO nucleic acid extraction and qRT-PCR

Matrigel domes from mature PDxO cultures were disrupted and washed twice in cold unsupplemented Advanced DMEM media. The pellet was lysed in RLT Plus with freshly added BME, and stored at −80 degree. For nucleic acid extraction, samples were incubated at 37 degree Celsius water bath for 5 min and vortexed for 1 min. Lysate was added onto a QiaShredder column and centrifuged. DNA and RNA was isolated from the flow-through using a Qiagen AllPrep kit following manufacturer’s instructions. RNA and DNA concentrations were determined by Qubit BR assays. RNA was cleaned of genomic DNA and reverse transcribed using a SuperScript IV VILO Master Mix with ezDNase Enzyme kit. 1 ng cDNA from each sample was run in 5 ul reactions in technical quadruplicate using PowerUp SYBR Green Master Mix against 500 nM of each forward and reverse primer (see Resource table). Reactions were run on a Roche LightCycler 480 using the PowerUp SYBR Green-recommended standard cycling mode (primer T_m_ ≥ 60°C). Ct values were normalized to the human GAPDH signal.

### Immunohistochemistry staining of PDX tumors and PDxO cultures

PDX tissues were fixed and processed as described previously^47^. For patient samples that were not solid tumors (e.g. pleural effusion or ascites fluid), cells were spun down and fixed in 2% PFA before embedding in histogel for sectioning. To prepare FFPE PDxO blocks, media was aspirated, and PDxO domes were mechanically disrupted in cold PDxO base medium. Organoids were washed once with cold PDxO base medium and resuspended in matrigel. The matrigel/organoid mixture was pipetted into a chamber slide and incubated at 37 degrees to solidify. To prepare histology blocks, a published protocol^74^ was followed, with the exception that the histogel for the bottom and top layer was warmed up to 65 degrees prior to pipetting. H&E staining of FFPE sections was performed to confirm PDxO content. H&E staining and IHC stainings were performed for ER, PR, HER2, pan-cytokeratin (CK-pan), human vimentin (hVim), E-Cadherin (E-cad), human mitochondria (hMito), mouse and human CD45, EpCAM and human specific cytokeratin (CK-CAM5.2). To quantify Ki67^+^ nuclei, three independent Ki67 staining sets were performed and the percentage of Ki67^+^ nuclei relative to hematoxylin-positive nuclei was quantified using ImageJ. For each staining set, one representative image per slide was quantified.

### Cytospin and immunofluorescence staining of PDxO cultures

For immunofluorescence staining, mature PDxO culture domes were gently disrupted and incubated in dispase solution (20% FBS in dispase with Y-27632) for 30 min at 37 degrees. PDxOs were washed twice with cold PDxO medium and fixed in PDxO fixing solution (2% PFA, 0.01% Tween20 in PBS) for 20 min at room temperature. After centrifugation, PDxOs were permeabilized in PBS with 0.5% Triton X-100 for 30 min. PDxOs were centrifuged and two aldehyde blocking steps were performed by incubating in freshly prepared aldehyde block solution (1 mg/ml NaBN_4_, 0.01% Tween20 in PBS) for 5 min. After resuspending in 0.01% Tween20 in PBS, PDxOs were centrifuged and washed once in 0.01% Tween20 in PBS. The PDxO pellet was resuspended in 1ml of PBS with 2% BSA. To prepare cytospin slides, 100 ul of PDxO solution was loaded and spun at 2000 rpm for 2 min at RT. Slides were dried overnight and ER/EpCAM double-IF staining was performed the next day. Cytospin slides were rehydrated in PBS containing 0.5% Triton X-100 for 20 min at RT, then washed in PBS and incubated in blocking buffer (50 mM NH_4_Cl, 10% goat serum, 5% BSA, 0.5% Tween20, 0.2% Triton X-100 in PBS) with mouse FcR Blocking reagent. For ER stains, PDxO cytospin slides were incubated with mouse-α human ER antibody. ER antibody was removed and α human EpCAM antibody was diluted in blocking buffer (10% goat serum, 5% BSA, 0.5% Tween20, 0.2% Triton X-100 in PBS) and incubated with M.O.M. diluent. Slides were washed in PBS with 0.05% Tween20 and incubated with secondary antibodies α mouse-AF555 and α rabbit-AF488, which were diluted 1:500 in blocking buffer with M.O.M. diluent. After washing, PDxOs were stained with DAPI and washed. Slides were incubated in 0.1% Sudan Black solution for 7 min to quench autofluorescence, then washed and covered with mounting media and cover slips.

### Quantification of live cell area and relative PDxO growth

To quantify PDxO live cell area and to assess relative PDxO growth, organoids were seeded as single cells in matrigel domes at a concentration of up to 11,300 cells per well onto matrigel base layers in a 48 well glass bottom plate (MatTek # P48G-1.5-6-F). For PDxO live cell area quantifications, 0.5 uM Calcein AM and 0.5 ug/ml Hoechst dye was added to each well and incubated for 1 hour at 37 degrees. Wells were washed once with HBSS and imaged using an inverted confocal microscope at the HCI Cell Imaging Core. The area of Calcein AM positive organoids was quantified using ImageJ. To quantify relative PDxO growth, CellTiter-Glo 3D (CTG-3D) cell viability assays were performed. Specifically, culture media was replaced with 250 ul HBSS and 100 ul CTG-3D was added per well. Matrigel domes were disrupted by pipetting and placed on a shaker in the dark for 20 min (500 rpm, room temperature). Plates sat at room temperature for an additional 10 min in the dark before samples were read on an EnVision XCite plate reader (PerkinElmer # 2105-0020).

### 3D drug screening assay of PDxOs

Mature organoids were harvested from culture using Dispase™ treatment at 37C for 20-25 min to remove residual matrigel. 5,000-10,000 cells (∼50 organoids) per well were seeded in TC-treated 384 well plates, each comprising a solidified 10 ul matrigel base layer and 30 ul of subtype-specific PDxO medium supplemented with 5% (by volume) matrigel. Seeded plates were incubated at 37 degrees C, 5% CO2, overnight. The following day, 8-dose drug plates were prepared by serial dilution and 30 ul of each condition, in technical quadruplicate, were transferred to seeded 384 well plates. Each 16-well set had 30 uL culture medium added immediately and was assayed with CTG3D to generate a seeding control value. Dosed plates were covered with Breathe-Easy seals and incubated for 96 hours (37C, 5% CO2). At the end of incubation, plates were assayed with CTG3D and read on an Envision system. Raw values from each condition were normalized to the day 0 seeding control value.

### Scoring dose-response curves for PDxO

One of the challenges we faced in this analysis was to compare drug responses across different organoids and subtypes. Each organoid model exhibited variable growth rates and responses to treatment. To overcome this challenge we calculated two scores based on a published growth rate normalization strategy^53^.This algorithm was implemented computationally using the four-parameter log-logistic function, self starter function (“LL.4”) largely based on using the “drc” R-library version (Ritz et al., 2015) which uses the following: Where b, c, d, e correspond to growth asymptote, the cytotoxic asymptote, the concentration at max response slope and the maximum slope of the mode, respectively. When screening results did not fit the sigmoidal curve, GRmetrics produced missing values for GR50 statistics. This typically occurred in organoids that did not respond to a compound and little or no cytotoxicity was observed.

### Genomic characterization of patient samples, PDX, and PDxO models

#### Whole-exome sequencing (WES)

Genomic sequencing for our models was performed at the Huntsman Cancer Institute High-Throughput Genomics and Bioinformatics Core. WES was conducted as follows: Agilent SureSelectXT Human All Exon V6+COSMIC or Agilent Human All Exon 50Mb library preparation protocols were used with inputs of 100-3000ng sheared genomic DNA (Covaris). Library construction was performed using the Agilent Technologies SureSelectXT Reagent Kit. The concentration of the amplified library was measured using a Qubit dsDNA HS Assay Kit (ThermoFisher Scientific). Amplified libraries (750 ng) were enriched for exonic regions using either the Agilent Technologies SureSelectXT Human All Exon v6+COSMIC or Agilent Human All Exon 50Mb kits and PCR amplified. Enriched libraries were qualified on an Agilent Technologies 2200 TapeStation using a High Sensitivity D1000 ScreenTape assay and the molarity of adapter-modified molecules was defined by quantitative PCR using the Kapa Biosystems Kapa Library Quant Kit. The molarity of individual libraries was normalized to 5 nM, and equal volumes were pooled in preparation for Illumina sequence analysis. Sequencing libraries (25 pM) were chemically denatured and applied to an Illumina HiSeq v4 paired-end flow cell using an Illumina cBot. Hybridized molecules were clonally amplified and annealed to sequencing primers with reagents from an Illumina HiSeq PE Cluster Kit v4-cBot (PE-401-4001). Following the transfer of the flow cell to an Illumina HiSeq 2500 instrument (HCS v2.2.38 and RTA v1.18.61), a 125-cycle paired-end sequence run was performed using HiSeq SBS Kit v4 sequencing reagents (FC-401-4003).

#### Sequence alignment and variant calling

Fastq files were uploaded to the PDXNet shared data pool for alignment and variant calling on the SevenBridges cloud interface (https://www.sevenbridges.com/). Collectively, the PDXNet community has established agreed-upon workflows for sequence alignment and variant identification. These workflows are implemented with CWL using SevenBridges under the direction and funding of the CGC^75^. When applicable, tumor-normal variants were processed using the “PDX WES tumor-normal workflow.” Briefly, this workflow consists of five st10 processed with Xenome for mouse read removal^76^. Second, tumor (PDX model) and normal FASTQ files are QC-checked using FASTQC^77^, trimmed by Trimmomatic^78^, aligned using BWA with alt-aware enabled, sorted using Picard SortSam^79^ and prepared for variant calling (Picard MarkDuplicates, GATK BaseRecalibrator, GATK ApplyBQSR). QC is also performed on the processed BAM files using Exome Coverage QC 1.0 tool and Picard CalculateHsMetrics. QC reports are aggregated with the QC Integrate script from MultiQC^80^. Third, BAM files were indexed using Samtools. Fourth, variant calling was performed using Mutect2^81^ and filtered using the GATK FilterMutectCalls function. Importantly, the ExAC 0.3 is used^82, 83^ for AF Of Alleles Not In Resource was set to 0.0000082364 (one mutation ExAC 0.3 ∼ 60,706 Exomes or 1/(2* 60706)). Finally, the workflow SnpEff/SnpSift tools (v4.3)^84, 85^ was run to acquire basic biological annotation for the VCFs produced. This identical process is also used for Tumor-only variant calls using Mutect2. We then converted VCF files into mutation annotation format (MAF, https://docs.gdc.cancer.gov/Data/File_Formats/MAF_Format/) using the software tool vcf2maf (https://github.com/mskcc/vcf2maf v1.6.17) which implements VEP release 95.3^86^.

#### Filtering WES data for cancer mutations

In order to remain consistent with data processing from available patient samples (specifically regarding availability of normal samples for germline analysis), we present two sets of mutation analysis, an in-depth set, and a PDX-only set of mutation data using tumor-only variant calls. Due to the possible misinterpretation of presenting likely passenger mutations within cancer genes, we restricted our mutations to previously identified cancer genes and mutations. First, missense mutations were only displayed if they met stringent criteria for predicted somatic driver genes, or predisposition germline mutations. Missense mutations found in predicted driver genes were only displayed if they were described in Tokheim et al. 2019 or Bailey et al. 2018^39, 40^. If a missense mutation was found in an established germline predisposition gene and associated with breast cancer samples^87^, we also required it to have a deleterious or likely-deleterious label by SIFT v5.2.2^88^; a damaging or likely-damaging label by PolyPhen v2.2.2^89^; a pathogenic or likely-pathogenic alteration by CLIN_SIG version 201810^90^; or a ‘HIGH’ impact rating by Ensembl release 95 ensembl-variation version 95.858de3e^86^. In addition to the genes reported by Huang et al., we included *ESR1* and additional genes involved in DNA damage repair including *FANCD2*, *FANCE*, *FANCF*, *FANCG*, *HMBS*, *POLD1*, *FANCI*, and *FANCL*. Lastly, missense mutations in germline predisposition genes were required to have a gnomAD population frequency less than 0.001 or be absent from gnomAD v170228^91^. We also chose to present nonsense, in-frame, frameshift, and predicted splice site mutations from previously predicted somatic cancer genes^39^ and germline predisposition genes^87^, only if their gnomAD population frequency was less than 0.001 or not reported. All mutations presented in figures were manually curated using IGV^92^ to ensure variant calling performance. When available, commercial genetic testing reports provided additional benchmarks for variant quality. If reports conflicted with our pipeline we rescued variants if we identified supporting reads in the alignment file. Finally, all mutations required a minimum variant allele fraction greater than 10% to remain included. We did not report silent mutations.

#### SNP array CNV

Each model also underwent analysis using the Illumina Infinium Omni2.5Exome-8 v1.3 or v1.4 or the Affymetrix SNP 6.0 array for profiling. These samples were also processed according to PDXnet specifications^1^. SNP array samples were genotyped with the 2.5 Exome-8 v1.3 Beadchip array. Hybridized arrays were scanned using an Illumina iScan instrument following the Illumina Infinium LCG Assay Manual Protocol and processed using GenomeStudio. Samples evaluated using the Affymetrix SNP 6.0 arrays were scanned using the Affymetrix 3000 7G Microarray scanner. The resulting images were processed using the Affymetrix GeneChip Operating Software version 1.4. When samples had both Affymetrix and Illumina chips, we deferred to Illumina intensity values for copy number calling. Tumor-only CNV calling was calculated using the tumor-only procedure as outlined by the author recommendations in ASCAT^93^. The copy number segments were then binned into 10kb windows to derive the median log2(CN ratio), which was subsequently used to re-center the copy number segments. Median-centered segments were then intersected with GRCh38 coordinates (Ensembl Genes 93) to represent copy number status for all intact genes. Where segmentation overlaps occurred, we selected the most conserved (lowest) CNV estimation. To account for median-centered differences between the Affymetrix and Illumina chip we calculated platform specific thresholds for amplifications and deletions (0.9 and −0.9 for Affymetrix CN ratios, and 0.6 and −0.6 for Illumina CN ratios; **Figure S60**). The gene set chosen for **Figure 1a** was curated from a set of four breast cancer or pan-cancer publications^94–95^, as well as sentinel genes, i.e., genes that represent a larger genomic region, from public and commercial sources including FoundationOne, cBioPortal, and Ambry Genetics Corporation. These extra genes include *FGF3, CDK12, NOTCH2, H3P6, SIPA1L3, ADCY9, FAM72C, SDK2, CDK18, STX4, TNFRSF10C, NKX3-1, LYN, JAK1, JAK2, CD274, PDCD1LG2, ERBB4, BARD1, BRCA1, BRCA2, BRIP1, CDH1, CHEK2, MRE11A, MUTYH, NBN, NP1, PALB2, RAD50, RAD51C, and RA051D*.

#### RNA sequencing

Due to the extended period of data collections, two different library preparation strategies were used for RNA-seq preparation: Illumina TruSeq RNA Library Preparation Kit v2 (RS-122-2001, RS-122-2002) and the Illumina TruSeq Stranded Total RNA Kit with Ribo-Zero Gold (RS-122-2301 and RS-122-2302). Briefly, the Poly-A capture library prep was processed as follows. Intact poly(A) RNA was purified from total RNA samples (100-500 ng) with oligo(dT) magnetic beads and sequencing libraries recommended by Illumina. Purified libraries were qualified on an Agilent Technologies 2200 TapeStation using a D1000 ScreenTape assay (cat# 5067-5582 and 5067-5583). The molarity of adapter-modified molecules was defined by quantitative PCR using the Kapa Biosystems Kapa Library Quant Kit (cat#KK4824). Individual libraries were normalized to 10 nM and equal volumes were pooled in preparation for Illumina sequence analysis. Alternatively, the Ribo-Zero RNA capture library was used for newer samples and was prepared as described following Illumina protocols. Briefly, total RNA samples (100-500 ng) were hybridized with Ribo-Zero Gold to substantially deplete cytoplasmic and mitochondrial rRNA from the samples. Purified libraries were qualified on an Agilent Technologies 2200 TapeStation using a D1000 ScreenTape assay (cat# 5067-5582 and 5067-5583). The molarity of adapter-modified molecules was defined by quantitative PCR using the Kapa Biosystems Kapa Library Quant Kit (cat#KK4824). Individual libraries were normalized to 10 nM and equal volumes were pooled in preparation for Illumina sequence analysis. Poly-A capture libraries were sequenced as follows: Sequencing libraries (18 pM) were chemically denatured and applied to an Illumina TruSeq v3 single read flowcell using an Illumina cBot. Hybridized molecules were clonally amplified and annealed to sequencing primers with reagents from an Illumina TruSeq SR Cluster Kit v3-cBot-HS (GD-401-3001). Following transfer of the flowcell to an Illumina HiSeq 2500 instrument (HCS v2.0.12 and RTA v1.17.21.3), a 50-cycle single read sequence run was performed using TruSeq SBS v3 sequencing reagents (FC-401-3002). The Ribo-zero capture libraries were sequenced as follows: Sequencing libraries (25 pM) were chemically denatured and applied to an Illumina HiSeq v4 paired end flow cell using an Illumina cBot. Hybridized molecules were clonally amplified and annealed to sequencing primers with reagents from an Illumina HiSeq PE Cluster Kit v4-cBot (PE-401-4001). Following transfer of the flowcell to an Illumina HiSeq 2500 instrument (HCS v2.2.38 and RTA v1.18.61), a 125-cycle paired-end sequence run was performed using HiSeq SBS Kit v4 sequencing reagents (FC-401-4003). Due to the variable technical approaches for obtaining RNA transcript abundance data, downstream batch correction strategies were required to correct for technical differences between platforms (see below).

#### Transcript abundance estimations

Like WES, all RNA-seq samples were processed as part of the CGC-SevenBridges cloud interface in accordance with PDXNet approved pipelines. These steps are briefly described here. First, fastq.gz files were decompressed for qc and mouse read removal purposes. Next, files were QC-checked using FASTQC^77^. Either single end or read-pairs were split (based on metadata and sequencing technique) for classification by Xenome for mouse read identification and removal^76^. Next, the workflow implements the RSEM Calculate Expression tool v1.2.31^96^ using the STAR aligner^97^. Finally, additional QC reports were generated by the Picard CollectRnaSeqMetrics tool (http://broadinstitute.github.io/picard/index.html). The resulting files from this work flow include transcript-specific results, aggregated gene level data, as well as RNA-seq bam files. Both the transcript-level and gene-level abundance results files capture RSEM expected counts, transcripts per million (TPM), and fragment per kilobase million (FPKM) estimates. For this manuscript, RSEM estimates were used for in-depth sample correlations, and comparative RNA-seq analyses.

#### RNA-seq count normalization correction strategies and analysis

RNAseq transcript abundance was used for PAM50 gene-based classification and in-depth sample correlation. We implemented the following steps to normalize transcript abundance data for these analyses. First, we normalized RSEM expected counts using DeSeq2^98^. Second, normalized counts were offset by 1 and log10 transformed. For gene-level comparison analysis, we performed root-mean-squared scaling for each gene. For correlation analysis, we wanted to remove likely housekeeping genes that may confound the relationship between samples. To accomplish this objective, we estimated the standard deviation of transcript abundance for each gene. We removed genes with a standard deviation less than 0.1, which left 80% of the dataset for pairwise Spearman correlations. The resulting correlation matrix was then leveraged to estimate intra- and inter-sample correlation differences. More specifically, we used a non-parametric Mann-Whitney U test to compare intra-sample correlation estimates (ignoring the diagonal) to the pairwise correlation estimates of other models. Finally, we used the ComplexHeatmap R^99^ to display all heatmaps.

#### scRNA sequencing

We extended our transcriptomic assessment of HCI-043 by performing single-cell RNA sequencing (scRNA-seq) on frozen PDX and PDxO tumor material. Samples were dissociated to single cells using the Miltenyi Human Tumor Dissociation Kit and GentleMACS dissociator. Samples were incubated on the GentleMACS at 37°C for 1 hour with 1,865 rounds per run. A 70 mm MACS smart strainer was used to deplete cell doublets before loading onto the 10x Genomics Chromium Controller. We used the 10X Genomics Single Cell 3’ Gene Expression Library Prep v3. Following library QC, 2 x 150bp sequencing was performed using Illumina’s NovaSeq6000 for 600-700M read-pairs per sample on S4 flow cells. We used Cell Ranger version 3.0.2 (10x Genomics) to perform scRNA-seq data alignment and gene abundance counting. We first aligned the fastq files from the PDX and PDxO samples to human-mouse hybrid reference hg19-mm10 provided by 10X Genomics. Cells were categorized into mouse cells, human cells, and multiplets by the Cell Ranger pipeline. This information was then later used to filter for human cells. We then aligned the same fastq files to human reference GRCh38-3.0.0 and generated single-cell expression tables. PDX and PDxO samples were aggregated using the Cell Ranger command “aggr” with depth normalization. Then, raw expression data were loaded into R version 3.5.2 and analyzed using the Seurat package version 3.1.0^100^. The criteria for filtering high quality cells for downstream analysis are as follows: 1) cells have more than 2,000 and less than 10,000 genes detected; 2) cells have less than 20% mitochondrial transcript counts; 3) cells are categorized as human cells. Next, we used “sctransform”^101^ for data normalization, “UMAP” for dimensional reduction, and graph-based clustering approach to cluster the cells.

#### Methylation sequencing

Reduced representation methylation sequencing was performed on 500ng of genomic DNA isolated from each sample as described previously^102^. Briefly, DNA is digested with MspI to enrich for CG rich regions of the genome. DNA fragment overhangs are filled with Klenow Fragment (3’-5’ exo-) (NEB) to leave an A-overhang. Adapters containing methyl-C are ligated to create a library. The library is treated with sodium bisulfite (EZ DNA Methylation Gold Kit; Zymo Research), or enzymatic methylation conversion (Enzymatic Methyl-seq Conversion Module, NEB Product #E7125L) to convert unmethylated cytosine to uracil. The converted library is PCR amplified using Taq polymerase and barcoded primers. During PCR uracil is copied as a thymine and methyl-cytosine is copied as cytosine. Equimolar quantities of barcoded libraries are pooled and sequenced on the illumina HiSeq 2500 or NovaSeq to obtain at least 50 million reads per sample. PhIX genome control library (Illumina) is included at 20-25% in each sequencing lane to improve cluster identification and base-balancing during sequencing. Sequencing data quality was assessed using FastQC v0.11.4 (http://www.bioinformatics.babraham.ac.uk/projects/fastqc/) to ensure the read starts with the MspI recognition sequence, and includes adequate base balancing (20-40% A,C,G and T at each cycle after the MspI recognition sequence). Adapters were trimmed from the sequencing reads using Trim Galore! v0.4.4 (http://www.bioinformatics.babraham.ac.uk/projects/trim_galore/), using the options --rrbs and --fastqc. Alignment to the hg19 reference genome was performed using Bismark v0.19.0^103^, using the options -bowtie2, --non_bs_mm, -N 1, and --multicore 6. Library quality was considered sufficient if greater than 60% of reads uniquely aligned to the genome. Conversion efficiency was considered sufficient when the percent of methylation observed in the CHH genome context was less than 3%. Library genome coverage was considered sufficient if more than 700,000 CG positions in the genome had a depth of at least 10 reads. For each library that met these quality metrics, methylation percentages at individual CpG positions in the reference genome were quantified using the Bismark Methylation Extractor v0.19.0 program, using the options --zero_based and --bedGraph. The pearson correlation in genome-wide methylation between samples was quantified and visualized using the cor function and the pheatmap package version 1.0.10 in the statistical software R version 4.0.2.

### ESR1 mutation testing

Assessment of the four hotspot ESR1 mutation in PDX tumors was conducted with droplet digital PCR (ddPCR). Briefly, genomic DNA (gDNA) and total RNA were extracted from each tumor using the Qiagen AllPrep DNA/RNA Mini kit. cDNA was synthesized from 500 ng RNA using PrimeScript RT Reagent Kit (Takara, RR037). A reaction mixture was prepared by mixing 50 ng gDNA or cDNA templates, ddPCR supermix for probes (no dUTPs; #1863024, Bio-Rad) and corresponding primer/probe set for specific ESR1 mutations (Y537S/Y537C/Y537N/D538G) as previously described^104^. Droplets were then generated using the QX100 Bio-Rad droplet generator with 20 μl reaction mixture and 70 μl droplet generation oil (#1863005, Bio-Rad). The ESR1 ligand binding domain fragment was amplified in each sample and signals from WT and mutant probes of each droplet were read using the Bio-Rad QX100 droplet reader. Mutation allele frequencies were further calculated using Quantasoft Software (#1864011, Bio-Rad). Positive and negative control droplets were included in each run to exclude potential contamination artifacts, and to control for proper gating of alleles. gDNA extracted from genome-edited MCF7 ESR1 mutant cells^105^ were used for positive controls of Y537S and D538G detection, and mutant DNA oligos were used for positive controls of Y537C and Y537N detection. gDNA from MCF7 ESR1 WT cells was used for negative controls.

### Generation of PDxoX

To generate PDxoX, the matrigel dome of mature PDxO cultures were gently disrupted by pipetting and transferred into a conical tube. After one wash step with cold PDxO base medium, the organoid pellet was resuspended in 20 ul matrigel on ice. Each mouse was injected with 20 ul of matrigel/organoid mixture into the mammary fat pad as previously described^47^.

### PDX drug treatments

For *in vivo* drug testing, SCID/Bg, NSG, or NRG mice were implanted orthotopically with PDX tumor fragments. NRG mice were used for most experiments with cytotoxic chemotherapy due to their increased tolerance for DNA damage relative to SCID mice. When tumors reached approximately 100 mm^3^ in size, drug treatment (or vehicle control treatment) was initiated. Tumor size was monitored 1-2 times per week, depending on the study. In some cases, treatment was stopped to see if tumors recurred, then restarted to determine resistance or continued sensitivity. Some of the *in vivo* drug testing experiments (docetaxel and navitoclax) were performed by the Patient-derived Xenograft and Advanced *In Vivo* Models (PDX-AIM) Core Facility of Baylor College of Medicine (M.T. Lewis, Director) in SCID/Bg mice, as a way to cross-validate results between PDxO drug responses and PDX responses in the same lines maintained at another institution.

## Supporting information

Supplemental Figures part 1

Supplemental Figures part 2

## Acknowledgments

We thank Carole Davis, Kelsey Embrey, Curi Bogert, Denise Spicer, and Kimberley Davenport for clinical data acquisition and Fadi Haroun, Brock Johnson, Shanna Kuhn, Dakota Simon, Taylor Roy, and Diane Hernandez for technical assistance. We also thank the University of Utah Health Science Centers Flow Cytometry and Cell Imaging Core facilities. We appreciate assistance from the NCI PDXNet Consortium for development of genomics data sharing and analysis pipelines through CGC/Seven Bridges Genomics.

## Supplementary Figure legends

**Supplementary Figure 1:**

Detailed description of the PDX establishment pipeline.

**Supplementary Figure 2:**

**(a)** Image showing a breast cancer metastasis to mouse liver that was used to establish PDX subline HCI-028LV. **(b)** Side-by-side genomics comparison of WES and CNV are presented for HCI-028 and HCI-028LV. **(c)** Immunohistochemistry of parental metastases and PDX llne. Stainings: H&E, ER, PR, HER2, CK-pan (Cytokeratin-pan), E-Cad (E-cadherin), hVim (human vimentin), msCD45, CK-CAM5.2 (human-specific cytokeratin CAM5.2), hCD45, EpCAM, and hMito (human mitochondria).

**Supplementary Figure 3:**

**(a)** Image showing a breast cancer metastasis to mouse ovary that was used to establish PDX subline HCI-031OV. **(b)** Side-by-side genomics comparison of WES and CNV are presented for HCI-031 and HCI-031LV. **(c)** Immunohistochemistry of parental metastases and PDX llne. Stainings: H&E, ER, PR, HER2, CK-pan (Cytokeratin-pan), E-Cad (E-cadherin), hVim (human vimentin), msCD45, CK-CAM5.2 (human-specific cytokeratin CAM5.2), hCD45, EpCAM, and hMito (human mitochondria).

**Supplementary Figures 4-40:** Immunohistochemistry of HCI PDX lines comparing the original patient tumors to the resulting PDX tumors. Stainings: H&E, ER, PR, HER2, CK-pan (Cytokeratin-pan), E-Cad (E-cadherin), hVim (human vimentin), msCD45, CK-CAM5.2 (human-specific cytokeratin CAM5.2), hCD45, EpCAM, and hMito (human mitochondria).

**Supplementary Figure 41:**

HCI-018 PDX tumor growth in mice implanted with E2 pellets only versus mice that received E2 pellets followed by E2 supplementation in drinking water.

**Supplementary Figures 42-45:**

Establishment of estrogen-independent (EI) ER+ PDX sublines. **(a)** Tumor growth of parental ER+ line under standard estrogen supplementation conditions (E2) versus in ovariectomized (OVX) mice with no E2 supplement. **(b)** Growth of the established EI subline arising from the OVX condition in panel (a), subsequently transplanted into OVX or intact mice with no E2 supplement. EI PDX lines are maintained in intact mice with no E2 supplement. For HCI-013 only, the tumor from the OVX condition in panel (a) was first expanded in culture for two weeks in phenol-red free medium with charcoal-stripped serum prior to implantation into OVX mice. Growth in intact mice is the subsequent passage. **(c)** Immunohistochemistry of estrogen-independent PDX HCI lines. Stainings: H&E, ER, PR, HER2, CK-CAM5.2 (Cytokeratin CAM5.2). Scale bar corresponds to 100 um.

**Supplementary Figure 46:** Droplet digital PCR detection of Y537S homozygous ESR1 mutation in HCI-044 PDX tumors **(a)** and Y537S low allele frequency tumors in HCI-018 PDX **(b)** and HCI-032 PDX **(c)**. 2D scatter plots of ddPCR results showing fluorescent detection of individual droplets with either gDNA or cDNA. Blue and green dots represent droplets with WT or mutant ESR1 genotypes indicated on the right panel of each plot, respectively. Orange dots represent droplets containing both WT and mutant ESR1 DNA. Black dots represent droplets without DNA. Mutation allele frequencies are labelled accordingly.

**Supplementary Figure 47:**

**(a)** Brightfield images tracking organoid growth in PDxO HCI-002. In the day 3 panel the * identifies a bubble trapped in the Matrigel during the embedding process, which gradually disappears during culture. In the day 12 panel the ** identifies a piece of debris, a common occurrence in the glass bottom plates required to acquire images. Scale bar represents 500 um. Far right, Calcein AM stain to show live cells (green). **(b)** Effect of various culture additives on cell viability, 13 days after first dissociation for PDxOs HCI-001 and HCI-002. **(c)** Effects of culture additives on cell viability for other TNBC PDxO lines, HCI-001 and HCI-015. **(d)** Comparison of doubling times during the first 60 days of culture to previously published organoid growth conditions. Statistical tests: (b) ordinary two-way ANOVA, uncorrected Fisher’s LSD with single pooled variance. **p = 0.004 *p = 0.01. (d) ordinary two-way ANOVA, uncorrected Fisher’s LSD with single pooled variance. **p = 0.002. Data in b-d are shown as mean ± SEM, with n=3 per condition.

**Supplementary Figure 48:**

**(a)** Growth response of HCI-011 PDxO to culture medium additives, quantified as live cell area. Error bars indicate +/− SEM. **(b-c)** Effect of other common breast cancer medium supplements on growth of HCI-011 PDxOs, quantified by Cell Titer Glo 3D (CTG3D) assay to measure ATP content. **(d)** HCI-011 PDxO organoid size (radius) after addition of Y-27632, NAC and FGF2 to PDxO cultures. Each gray dot represents one organoid, *p < 0.05, ***p < 0.001. **(e)** Comparison of culture doubling time of HCI-011 and HCI-017 when established in our optimized ER+ PDxO media or organoid media published by Sachs et al., 2018. *p < 0.05, error bars indicate +/− SEM.

Statistical tests: (b) ordinary two-way ANOVA, uncorrected Fisher’s LSD with single pooled variance. (d) ordinary one-way ANOVA, uncorrected Fisher’s LSD with single pooled variance; (e) ordinary two-way ANOVA, uncorrected Fisher’s LSD with individual variances computed from each comparison. *p < 0.05, **p < 0.01,***p < 0.001, error bars indicate +/− SEM

**Supplementary Figure 49:**

**(a-b)** mRNA expression level of *ESR1* and its downstream target *TFF1* in ER^+^ PDxO lines HCI-003 (a) and HCI-017 (b) over time in culture compared to HCI-011 PDX, PDxoX or MCF7 or T47D cell lines cultured in 2D or 3D for 6 days. Data are normalized to human GAPDH Ct. **(c)** *TFF1* expression in ER^+^ HCI-003 PDxOs and HCI-017 PDxOs stimulated with E2 for 8h after 4 days of culturing in phenol red-free medium with charcoal-stripped FBS. Data are normalized to human GAPDH Ct and the control condition (no E2). **(d)** Relative live cell area in ER^+^ HCI-003 and HCI-017 PDxOs stimulated with E2 for 8h after 4 day starvation as in panel c. Data are normalized to the control condition (no E2). **(e)** Immunofluorescent staining of ER (green) and EpCAM (red) on cytospin slides of HCI-003 and HCI-017 PDxOs without E2 stimulation. Scale bars represent 75 um. **(f)** PDxO xenograft (PDxoX) tumor growth response of HCI-003 (left) and HCI-017 (right) to 40 mg/kg or 200 mg/kg fulvestrant treatments versus vehicle controls. Data shown are relative to 100% tumor volume at treatment start, (HCI-003 n=3 for vehicle, n=3-4 for treatment; HCI-017 n=3 for vehicle, n=2-4 for treatment). **(g)** Tumor growth rate for data in panel g. **(h-i)** Re-treatment of PDxO xenograft (PDxoX) HCI-003 (h) and HCI-011 (i) with 200 mg/kg fulvestrant after off-treatment recurrence, in order to select resistance. Graph shows individual tumors. Statistical tests: (c, d) Ordinary two-way ANOVA, uncorrected Fisher’s LSD with single pooled variance;. (f) *p < 0.05, **p < 0.01,***p < 0.001, error bars indicate +/− SEM.

Statistical test: ordinary two-way ANOVA, uncorrected Fisher’s LSD with individual variances computed from each comparison. (g) ordinary one-way ANOVA, Tukey’s multiple comparison with single pooled variance *p < 0.05, error bars indicated +/− SEM.

**Supplementary Figure 50:**

**(a)** Immunohistochemistry of HER2 showing variable HER2 staining in PDxOs and their parental PDX tumor with HER2+ histories. HCI-005 (top) and HCI-008 (bottom) are shown. The scale bar corresponds to 50 um.

**(b)** Effect of additional medium supplements on growth of HCI-005 PDxOs, quantified by CTG3D assay (left) and live cell area (right). Statistical test: ordinary two-way ANOVA, uncorrected Fisher’s LSD with single pooled variance. *p < 0.05, **p < 0.01,***p < 0.001, error bars indicate +/− SEM

**Supplementary Figure 51:**

Expression of genes used to characterize PDxO cultures over time. HCI-001, HCI-010 and HCI-027 PDxOs are shown at different culture time points; technical replicates of n=4. Data are normalized to human GAPDH Ct.

**Supplementary Figure 52:**

PDxO xenograft (PDxoX) tumor volumes (upper panel) and tumor growth rates (middle panel) compared to parental PDX lines for HCI-001 **(a)**, HCI-010 **(b)**, HCI-025 **(c),** and HCI-027 **(d)**. PDxOs were in culture for 51-64 days (PDxoX early) or 113-123 days (PDxoX late) prior to xenografting. Quantification of Ki67+ nuclei of parental PDX (lower), early (51-68 days) and late (113-127 days) PDxO cultures and early and late PDxoX. Data are averaged from 3 individual sets of IHC staining, normalized to total hematoxylin^+^ nuclei count for each image. All data are shown as mean ± SEM with technical replicates of n=4. Statistically significant differences for Ki67 quantification are indicated by *p < 0.05, **p < 0.01,***p < 0.001.

Statistical tests growth rates: ordinary one-way ANOVA, Tukey’s multiple comparisons test with a single pooled variance. HCI-010 PDX only generated n=1, requiring use of two-tailed unpaired t test; statistical tests Ki67 quantification: ordinary one-way ANOVA, Tukey’s multiple comparisons test with a single pooled variance.

**Supplementary Figure 53:**

Histology showing H&E stains of PDxO lines HCI-001, HCI-002, HCI-010, HCI-025 and HCI-027. Stains of the parental PDX tumor or PDxOs cultured for 0, 60 or >120 days are shown. PDX tumors retain morphology when compared to re-implanted PDxoX after being cultured as PDxoX for 60 or 120 days. Scale bar corresponds to 50 um.

**Supplementary Figure 54:**

Immunohistochemistry showing CK-CAM5.2 stains for PDxO lines HCI-001, HCI-002, HCI-010, HCI-025 and HCI-027. Stains of the parental PDX tumor or PDxOs cultured for 0, 60 or >120 days are shown. Scale bar corresponds to 50 um.

**Supplementary Figure 55:**

Immunohistochemistry showing Ki67 stains for PDxO lines HCI-001, HCI-002, HCI-010, HCI-025 and HCI-027. Stains of the parental PDX tumor or PDxOs cultured for 0, 60 or >120 days are shown. Scale bar corresponds to 50 um.

**Supplementary Figure 56:**

Immunohistochemistry showing human vimentin stains for PDxO lines HCI-001, HCI-002, HCI-010, HCI-025 and HCI-027. Stains of the parental PDX tumor or PDxOs cultured for 0, 60 or >120 days are shown. Scale bar corresponds to 50 um.

**Supplementary Figure 57:**

**(a)** CNVkit segmentented copy numbers plot^106^ illustrates log2 CN ratios for the indicated models. The annotations (left) display HCI-line, model, passage, technology (i.e., SNPchip platform). Segmented copy number data is presented as log2 CN ratios indication amplifications (red) and deletions (blue). **(b)** Density plot of log2 CN ratios are colored according to the sequencing platform. Vertical bars indicate the thresholds set to define discrete amplifications and deletions.

**Supplementary Figure 58:**

Sixteen heat maps illustrate individual drug responses to 45 compounds. Coloration of these heatmaps indicates CellTiter-Glo 3D cell viability assays that were normalized to day 0, ranging from 0 (black, cytotoxic) to 2+ (yellow, growth). Values at 1 are considered cytostatic. Color scaling is performed relative to each model. Drugs are sorted left to right from largest GRaoc to smallest, indicating decreasing drug efficacy in each model, respectively.

**Supplementary Figure 59: Biological replicates exhibit reproducible drug responses**

**(a)** Two compounds, 5-fluorouracil and FK866 were screened at eight-point dose response curves (y-axis) at different days in culture (x-axis). Individual replicates and dose responses from a CellTiter-Glo 3D cell viability assay were normalized to day 0 for the model HCI-001. Values range from 0 (black) to 4 (light-yellow), where zero indicates cytotoxic effects and yellow shows growth phenotypes. Values at 1 are considered cytostatic. **(b)** As in the previous panel two compounds, ibrutinib and romidepsin, were screened in model HCI-002 at different times in culture. Here, response values range from 0 (red) to 7 (light-yellow). Values at 1 are considered cytostatic. **(c)** Individual biological replicates for 16 models are shown for two different targeted therapies in the Notch pathway (LY3039478 and RO4929097), as well as two targeted therapies in the mTOR pathway (vistuserib and sapanisertib). Colored as previously described in panel (a). Response values range from 0 (black) to 5 (light-yellow). Values at 1 are considered cytostatic.

**Supplementary Figure 60:**

**(a)** An illustration of dose response curve statistics that can be calculated using the R-package GRMetrics. The y-axis displays growth rate adjusted estimates from the CellTiter-Glo 3D cell viability assays. The x-axis shows log-fold change of 8-point dose concentrations. Each dot represents one of 12 replicates (3 biological replicates, with 4 technical replicates each). Annotations include EC50 (half maximal effective concentration); GR50 (concentration at which GR value = 0.5); cytostatic (concentration at which model is neither growing nor shrinking); and GRaoc (area over the dose response curve that estimates both sensitivity and cytotoxicity). **(b)** Ordered models based on GRaoc for navitoclax sensitivity. High values with darker colors suggest cytotoxic response to the compound. The color of model identifiers correspond to *in-vivo* data in panels d and e. **(c)** Heatmap displays drug response to navitoclax in PDxO screens. The coloration indicates CellTiter-Glo 3D cell viability assays in PDxO screens that were normalized to day 0 ranging from 0 (black, cytotoxic) to 3 (yellow, growth). Models are sorted by GRaoc estimate. **(d, e)** Responses to navitoclax *in vivo* for the most sensitive model predicted by PDxO screening (HCI-010; panel d), and four others: HCI-024, HCI-015, HCI-002 and HCI-027 (panel e) Data represent n=3-6 mice per group.

**Supplementary Figure 61:**

**(a)** Stacked heat map displays ordered GRaoc calculations for each model’s response to docetaxel from dark (cytotoxic) to light (growth). The color of model identifiers correspond to *in-vivo* modeling in panel c. **(b)** Heatmap displays drug response to docetaxel in PDxO screens. The coloration indicates CellTiter-Glo 3D cell viability assays in PDxO screens that were normalized to day 0 ranging from 0 (black, cytotoxic) to 2.5 (yellow, growth). Models are sorted by GRaoc estimate. **(c)** Results of *in vivo* treatment for HCI-023, HCI-015, HCI-019, HCI-016, HCI-002, HCI-027, HCI-010, HCI-024, and HCI-001. Data represent n=3-6 mice per group.

**Supplementary Figure 62:**

**(a)** Stacked heat map displays GRaoc calculations for each model’s response to the gamma-secretase inhibitor RO4929097 from dark (cytotoxic) to light (growth). The color of model identifiers correspond to *in-vivo* modeling in panel c. **(b)** Heatmap displays drug response to RO4929097 in PDxO screens. The coloration indicates CellTiter-Glo 3D cell viability assays in PDxO screens that were normalized to day 0 ranging from 0 (black, cytotoxic) to 2 (yellow, growth). Models are sorted by GRaoc. **(c)** *In vivo* drug treatment of PDXs HCI-015, HCI-027, HCI-010, and HCI-002 with RO4929097.

**Supplementary Figure 63:**

Synergy plot as a result of drug treatment of birinapant and SN-38 in HCI-002 (left) and HCI-023 PDxOs (right). Blue indicates synergy and red indicates antagonism. The number in the box represents the Lowe synergy score +/− standard deviation; *p<.05, **p<.001, ***p<.0001.

**Supplementary Figure 64:**

**(a)** Representative imaging data (upper panel) and a timeline representing the natural history of patient HCI-043’s breast cancer (lower panel) Events are as follows: A. Diagnosis of early recurrent disease metastatic to the liver (solid arrow). No skeletal metastases (empty arrow). B. No response to capecitabine; new onset skeletal metastases (empty arrow). C. Initiation of cabozantinib and atezolizumab; liver metastases still present (solid arrows). D. No response to cabozantinib and atezolizumab; progression of the hepatic metastases (solid arrow) and production of malignant ascites (empty arrows). E. After 3 cycles of the PDxO-informed eribulin treatment, the patient achieved a complete radiographic remission of the hepatic metastases (solid arrows). The malignant ascites also regressed somewhat (empty arrow). F. After 5 cycles on eribulin, there was complete remission of the malignant ascites (empty arrow) and continued complete remission of the hepatic metastases (solid arrow). However, new onset isolated metastasis in T12 vertebrae (arrowhead) required discontinuation of eribulin and treatment with radiation therapy. G. Recurrence of the hepatic metastases 2 months after withholding the eribulin (solid arrows). **(b)** H&E staining and PD-L1 staining of HCI-043 patient’s tumor. The tumor tested low but positive for PD-L1 on the basis of an FDA-approved commercially available test. **(c)** (upper panel) RNA-seq data showing expression of genes associated with the luminal androgen receptor (LAR) subtype in HCI-043 patient tumor. (lower panel) scRNA-seq data showing androgen receptor (AR) expression (red; left side) in all tumor cell clusters (middle) in HCI-043 PDX and PDxOs. Immunohistochemistry for the androgen receptor on the patient tumor was detected by a commercial vendor (PhenoPath; right side). **(d-e)** Dose response heatmaps showing results of drug screening on the pre-treatment breast tumor biopsy model HCI-043 (d) and the post-treatment metastatic ascites sample (e) from timepoint D on the timeline in panel a, HCI-051. Coloration of these heatmaps indicates CellTiter-Glo 3D cell viability assays that were normalized to day 0 ranging from black (cytotoxic) to yellow (growth), which have been scaled respectively. The drug order on both plots is sorted by GRaoc.

## KEY RESOURCES TABLE

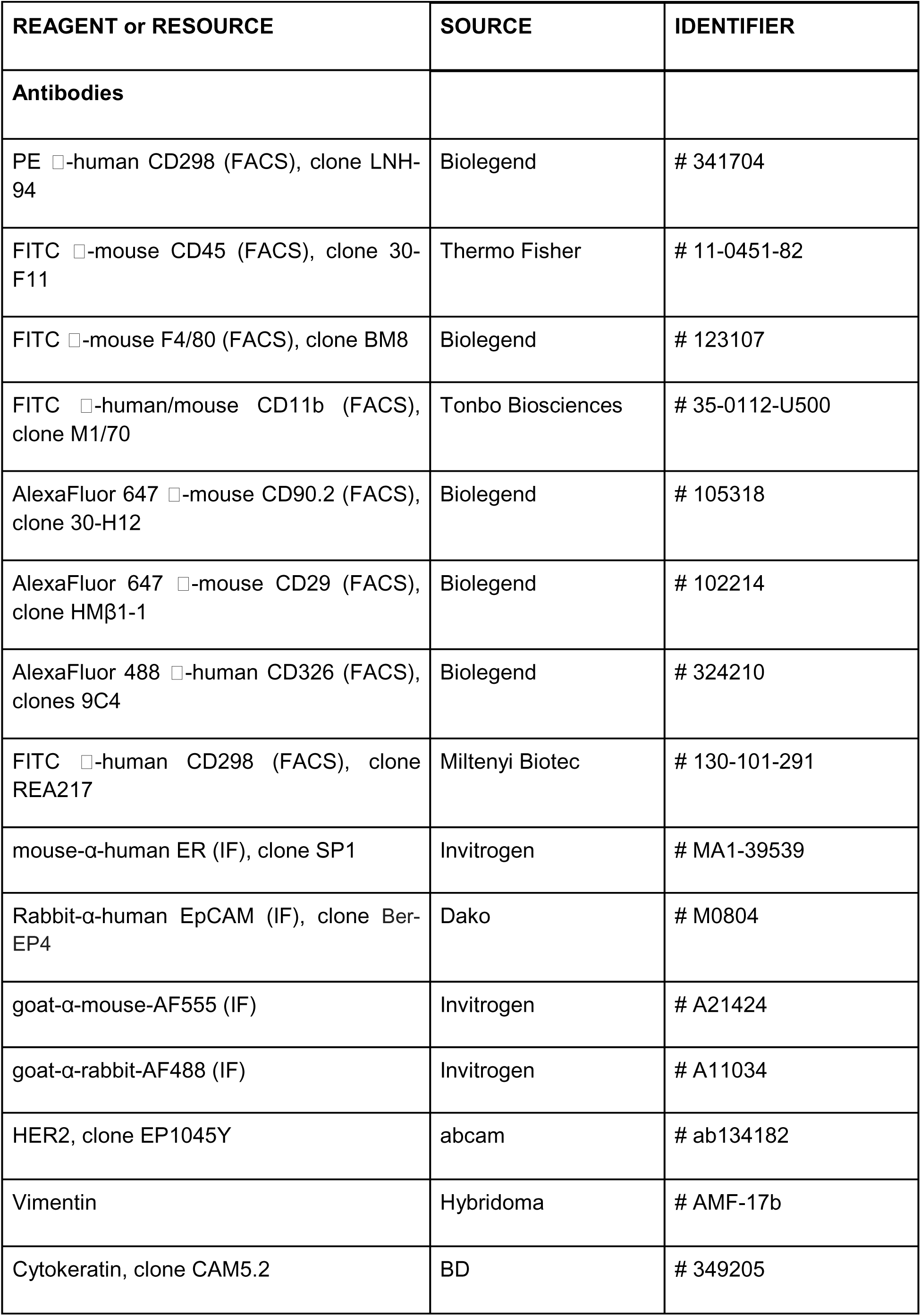
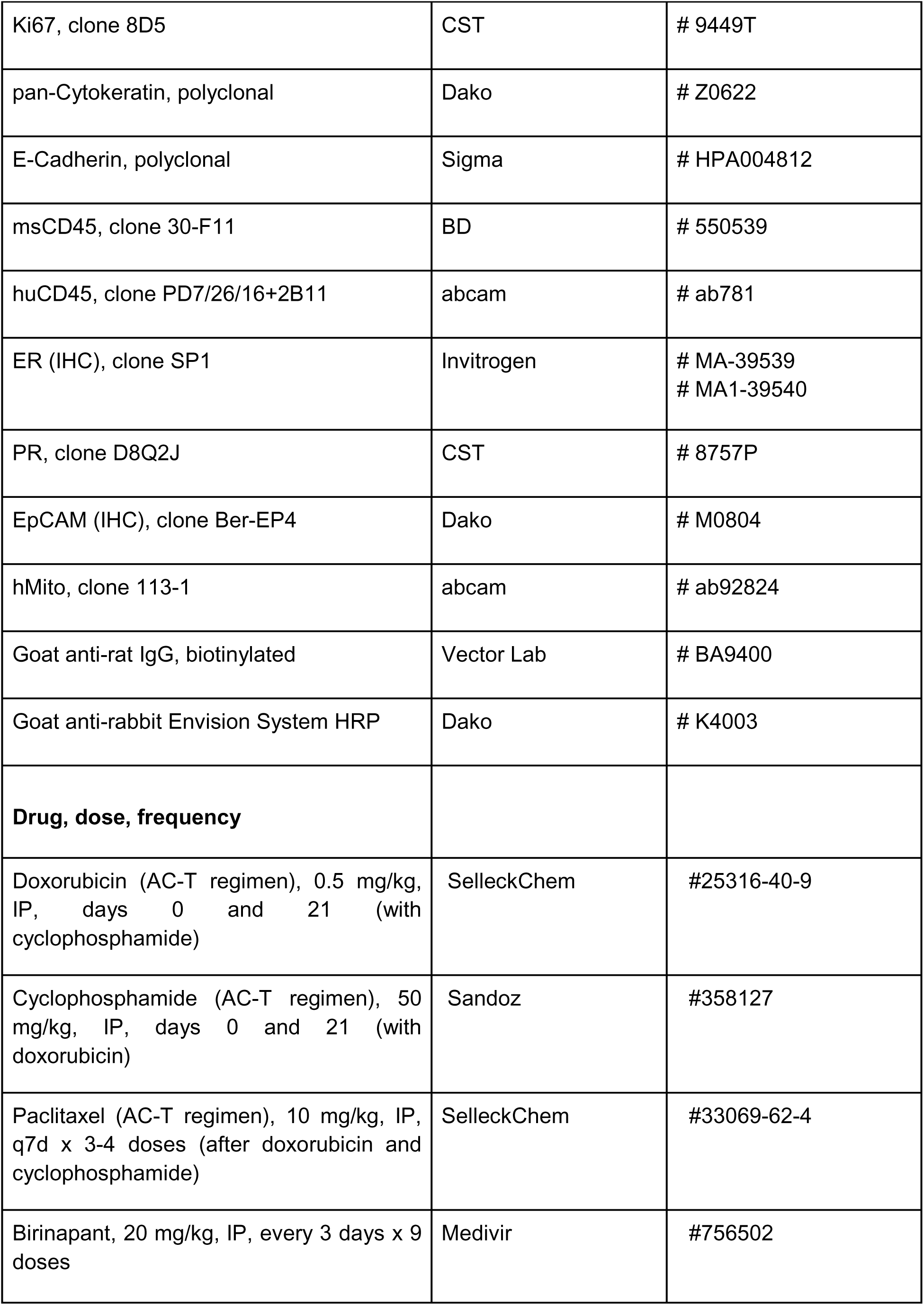
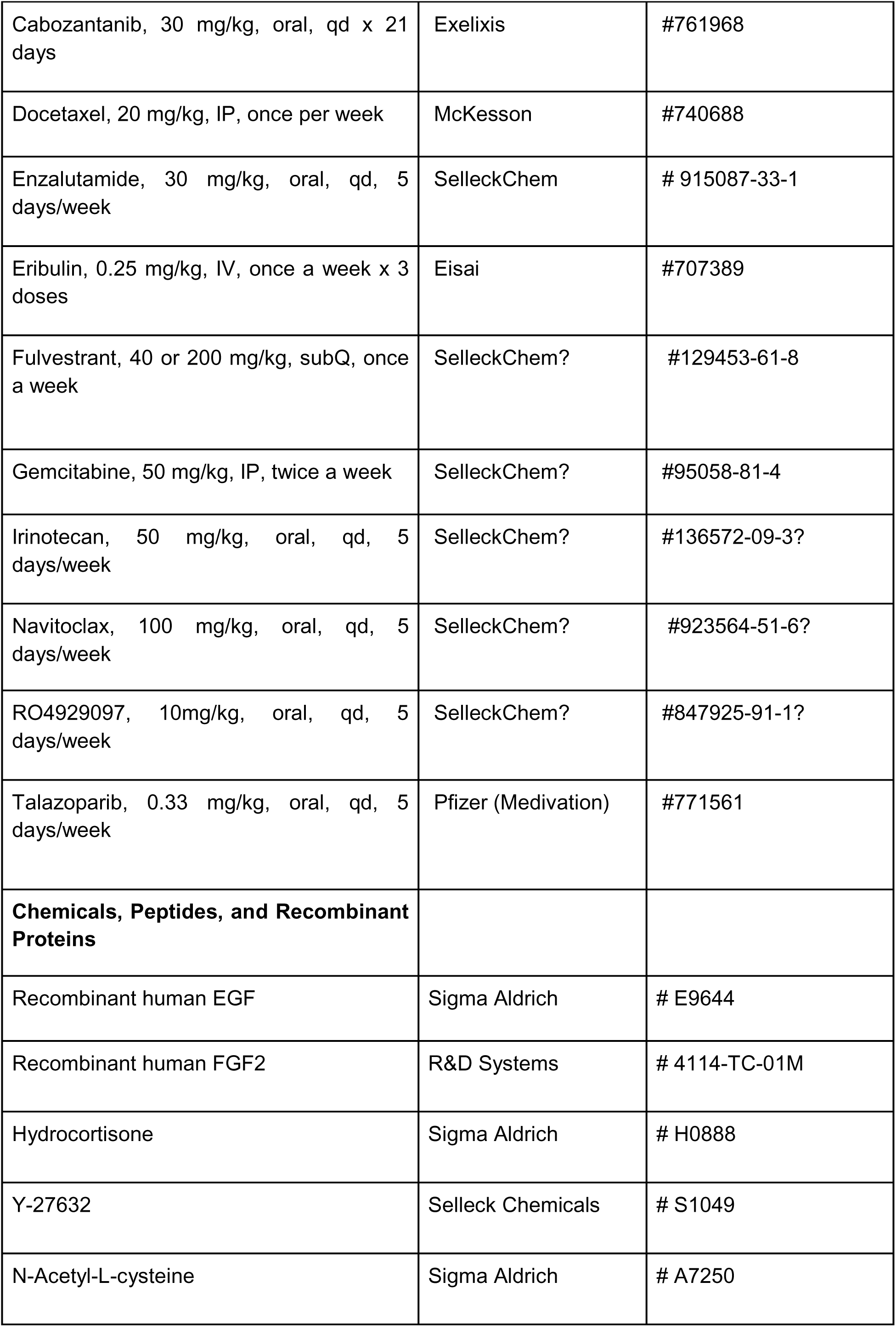
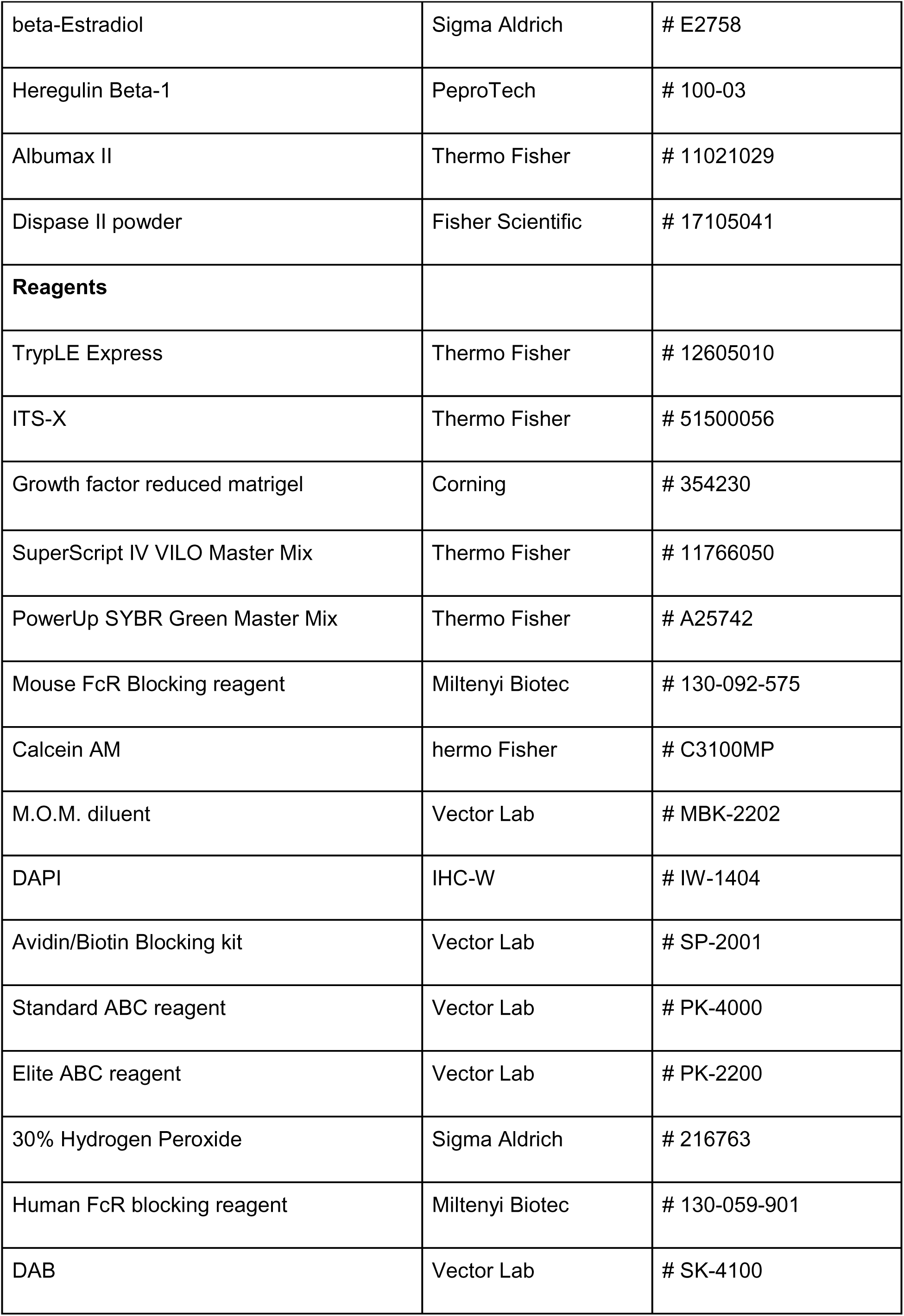
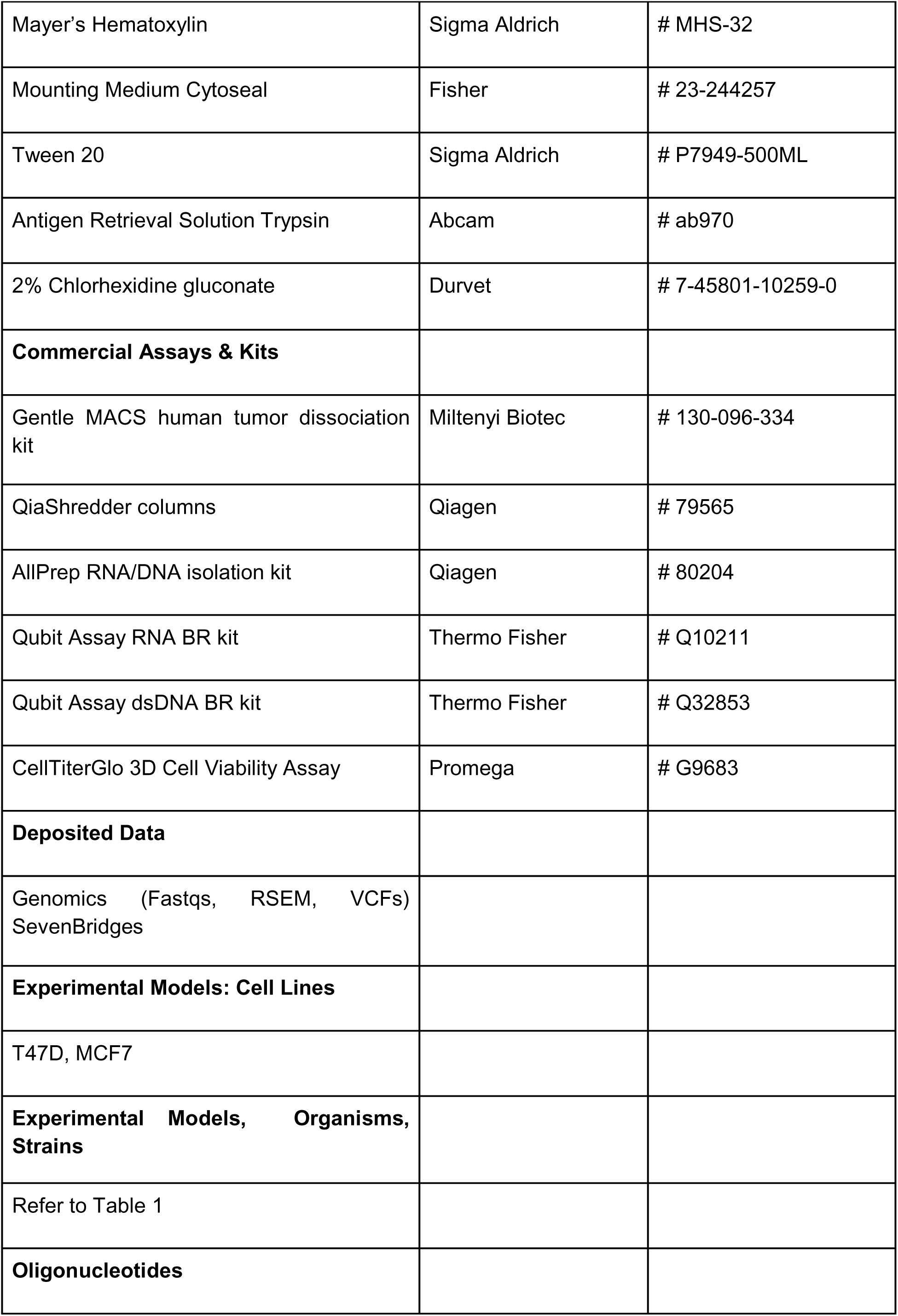
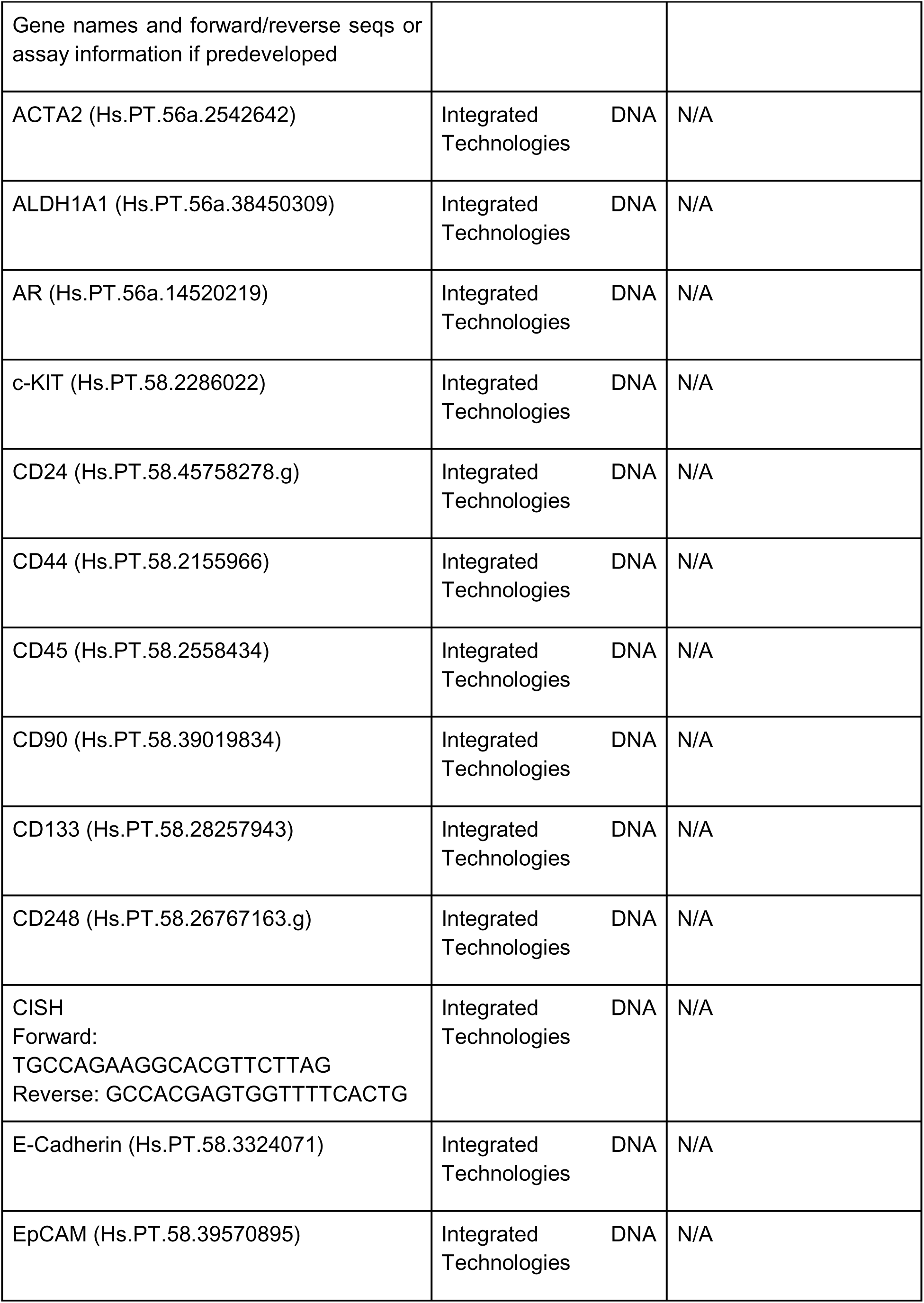
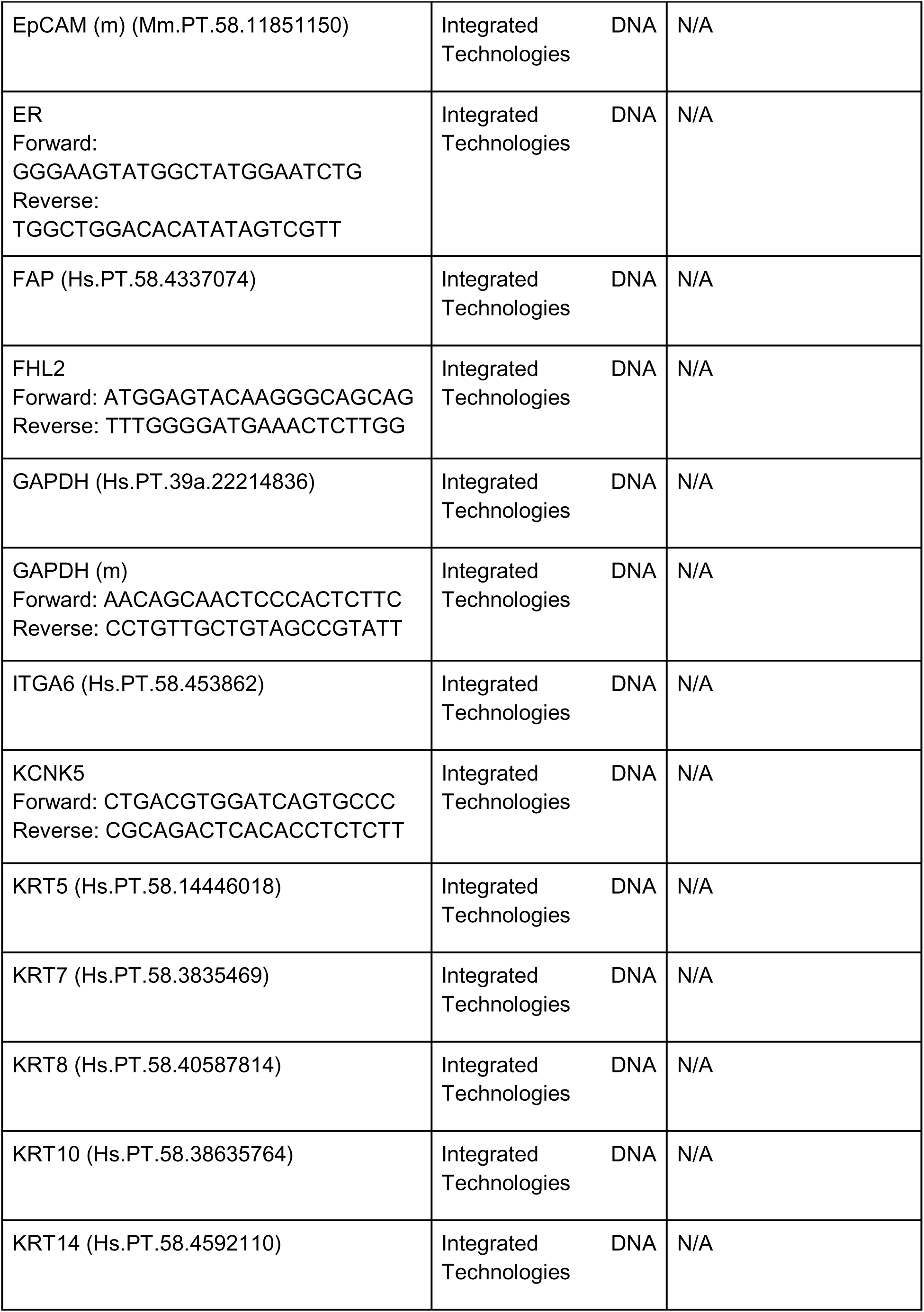
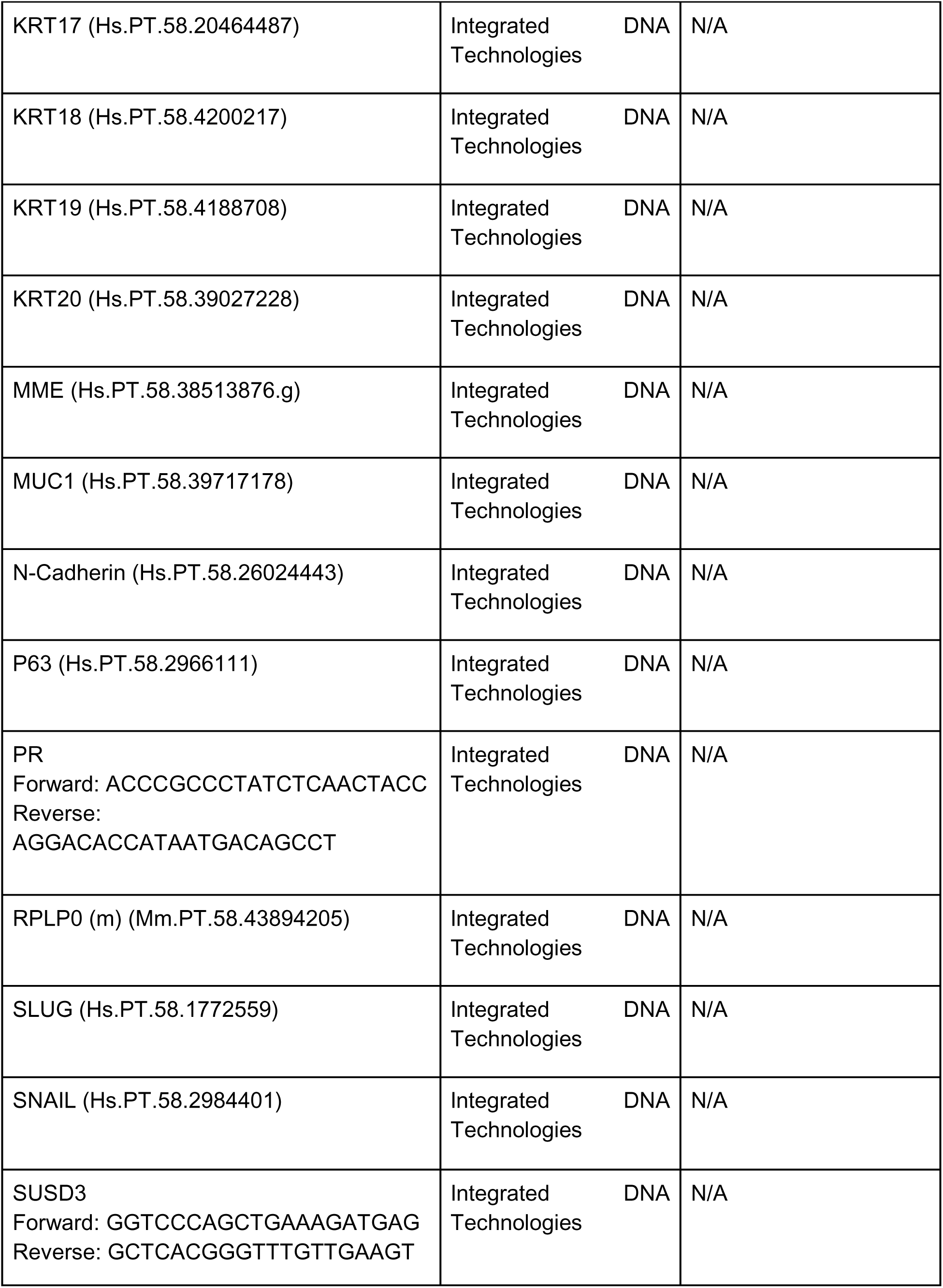
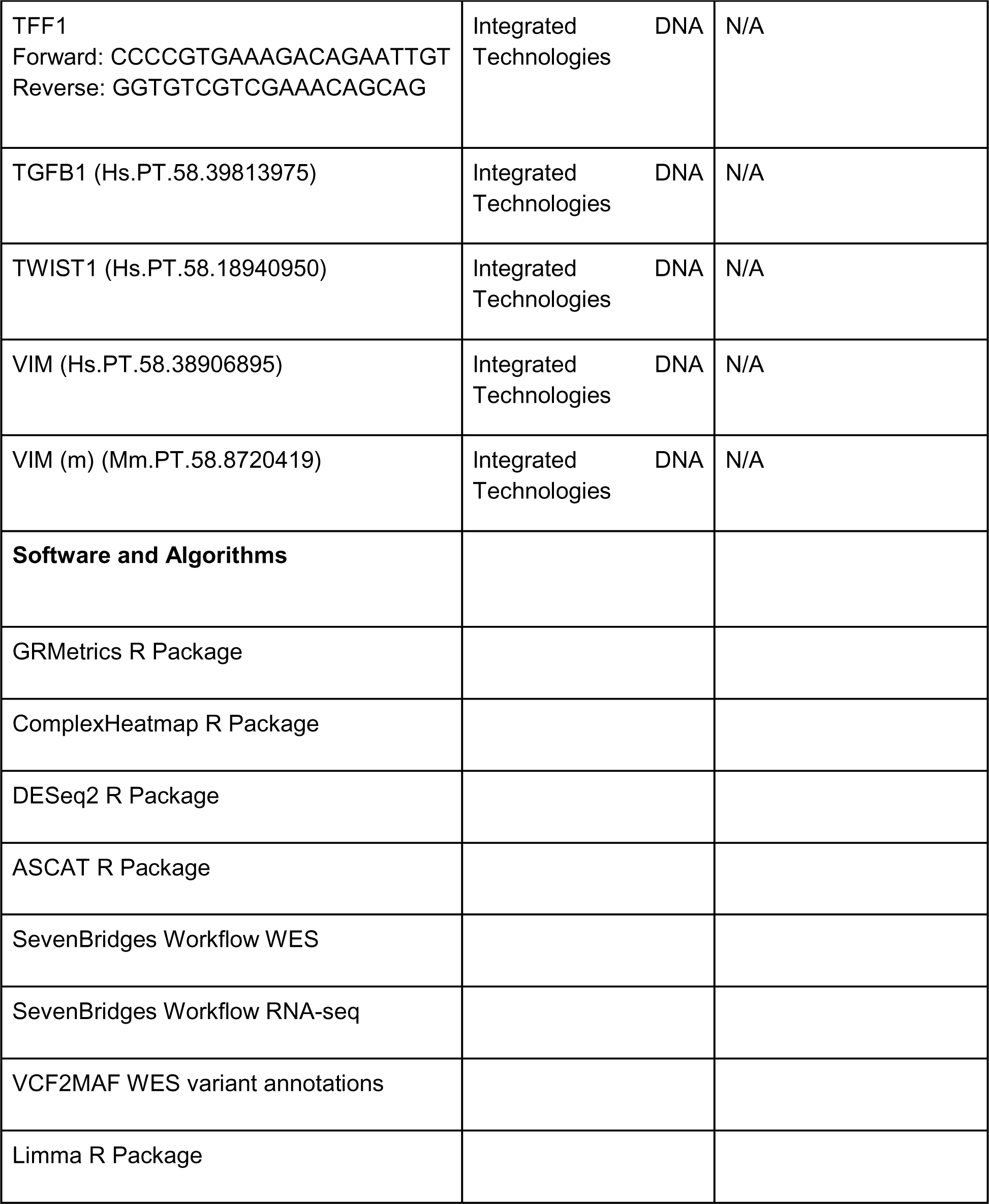

