## Supplemental Figures part 1 for "A breast cancer patient-derived xenograft and organoid platform for drug discovery and precision oncology"

Supplementary Fig. 1

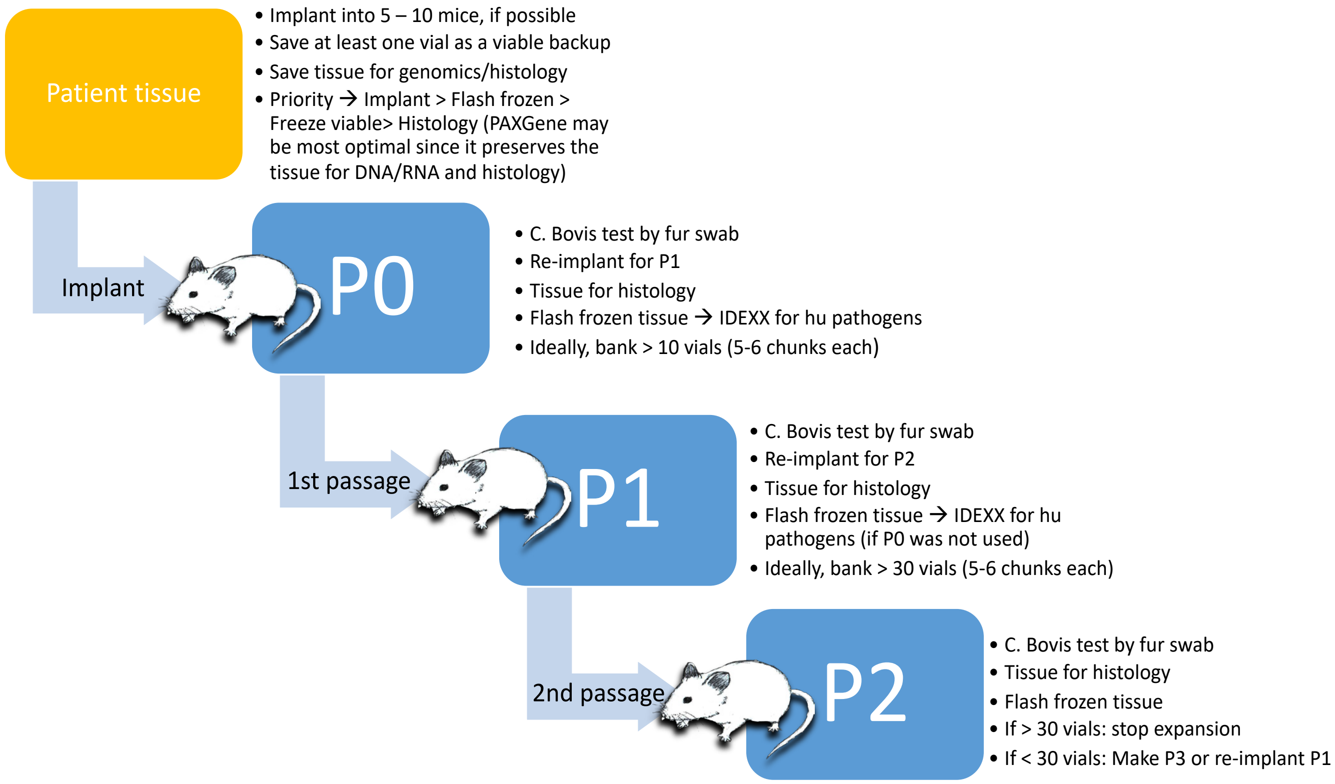

Supplementary Fig. 2

#### HCI-028LV

a

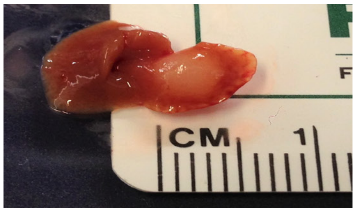


b

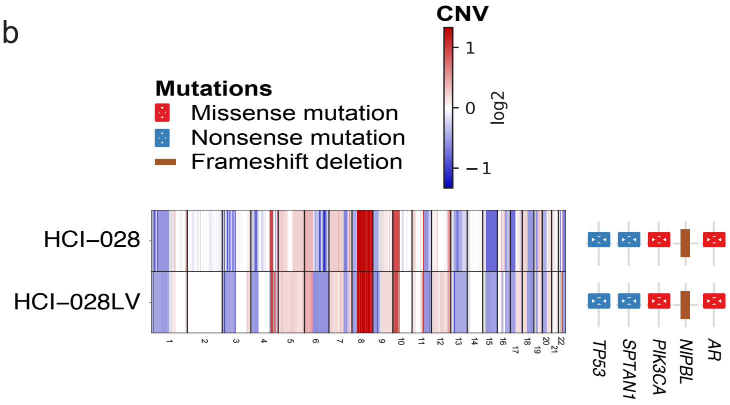


c

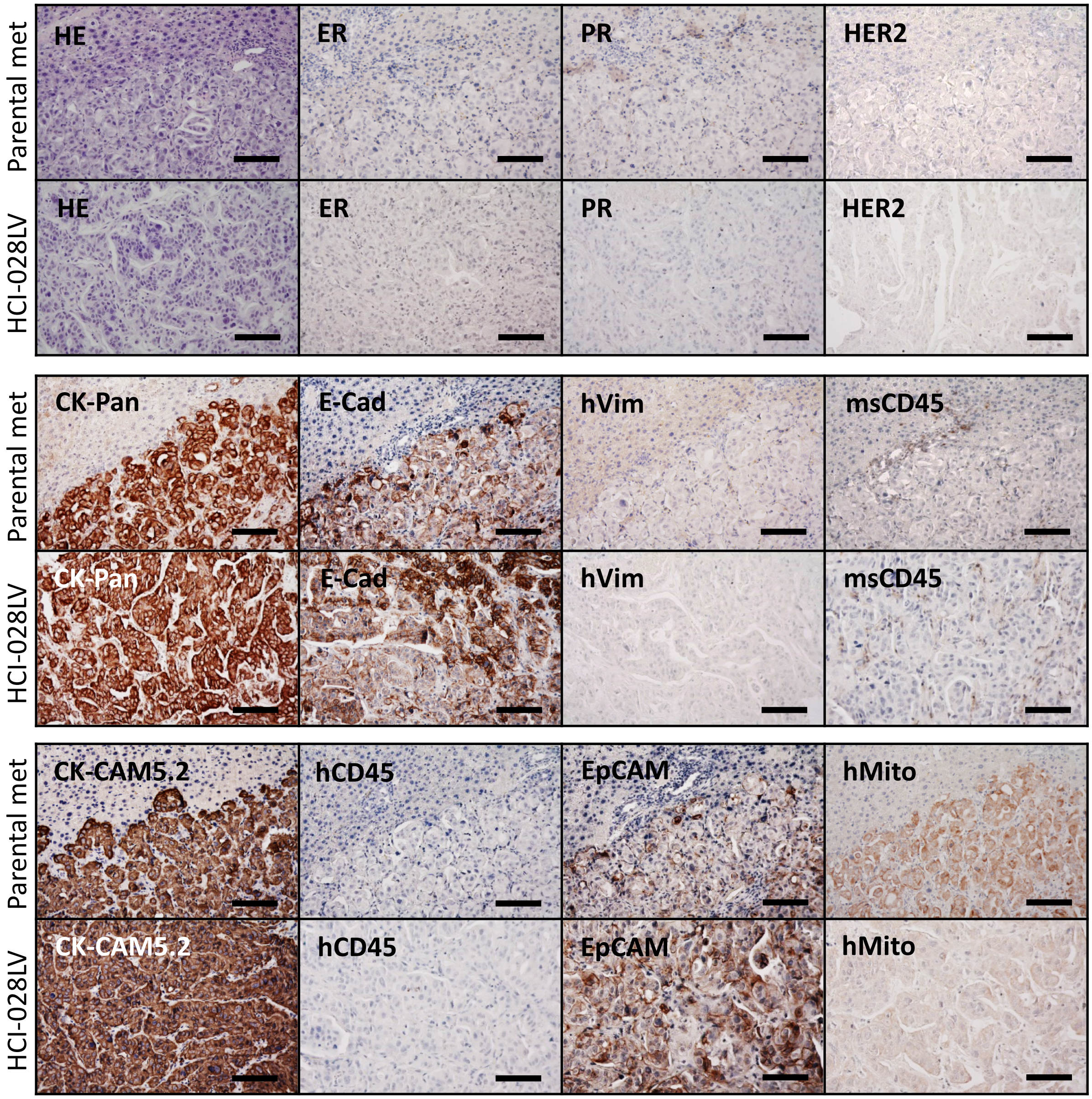


Scale bars = 100 μm

Supplementary Fig. 3

**HCI-031OV**

a

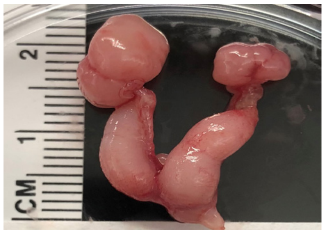


b

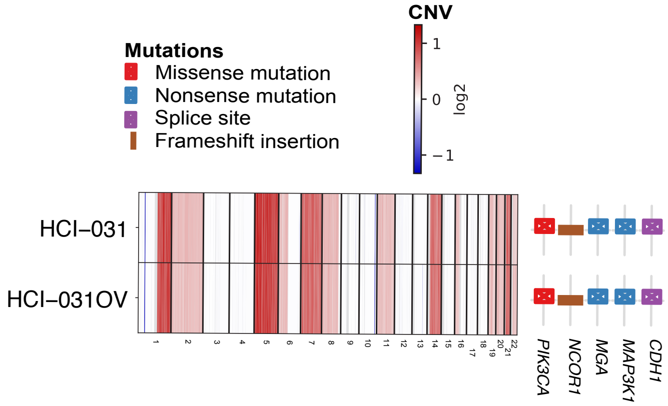


c

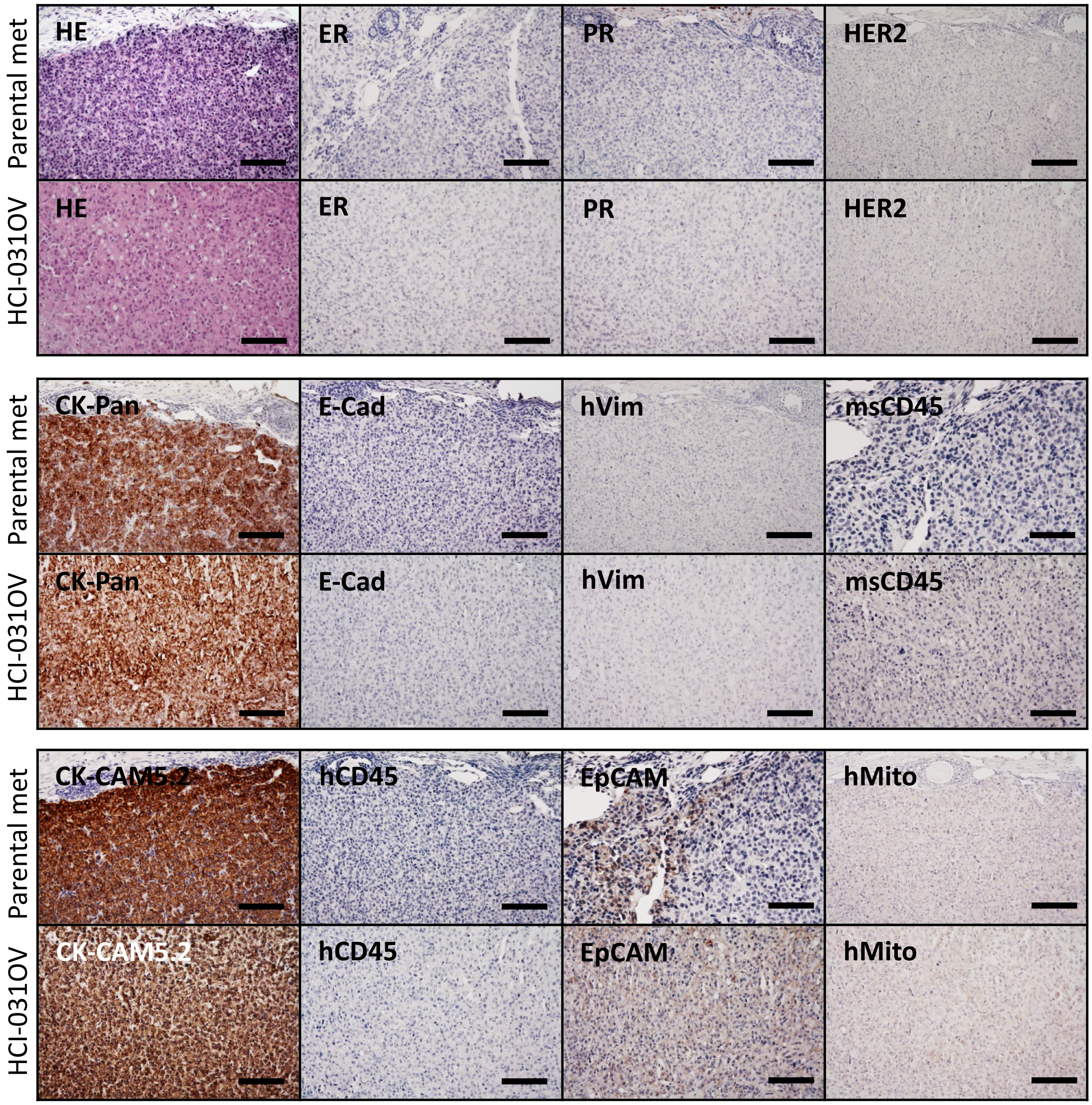


Scale bars = 100 µm

Supplementary Fig. 4

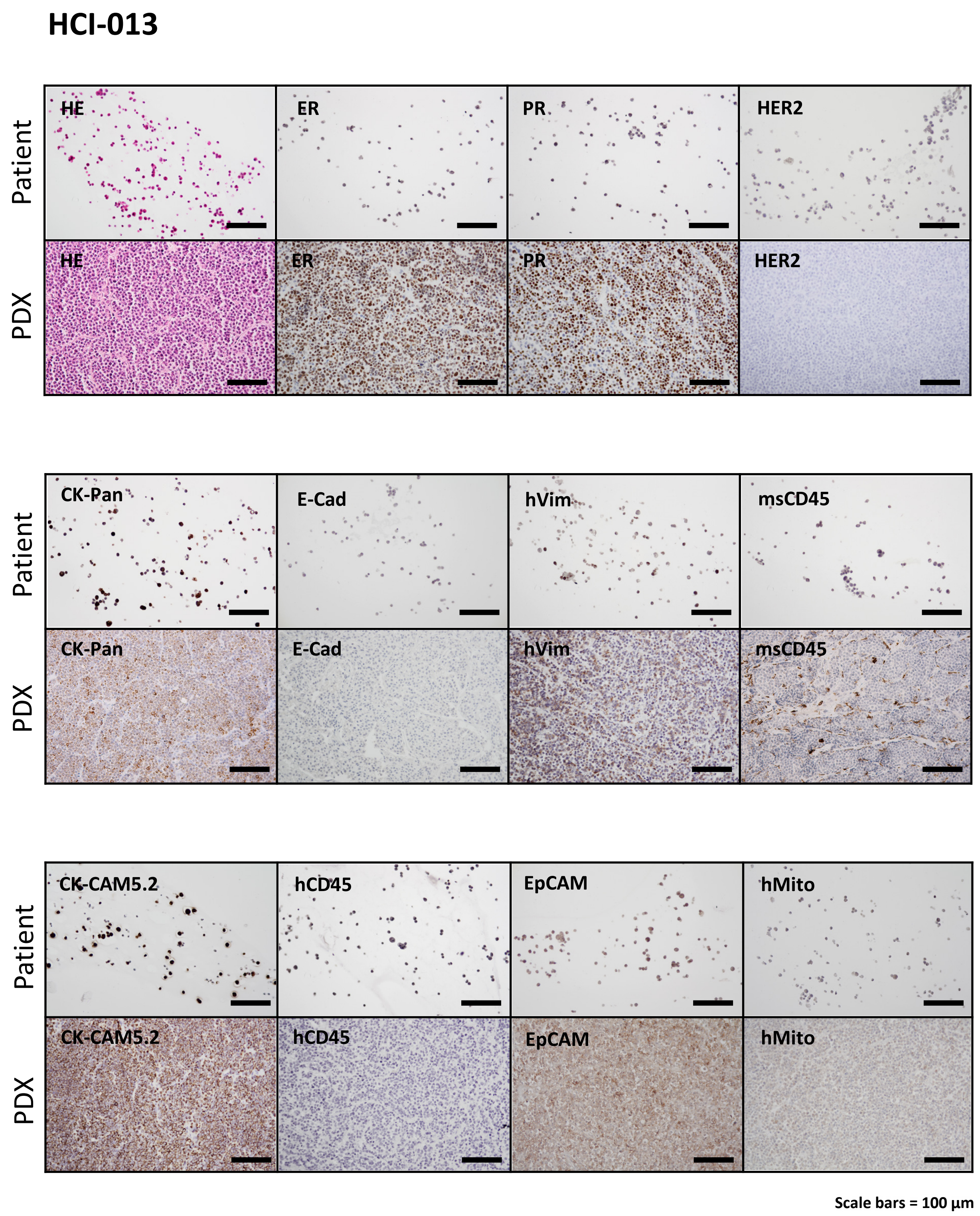

Supplementary Fig. 5

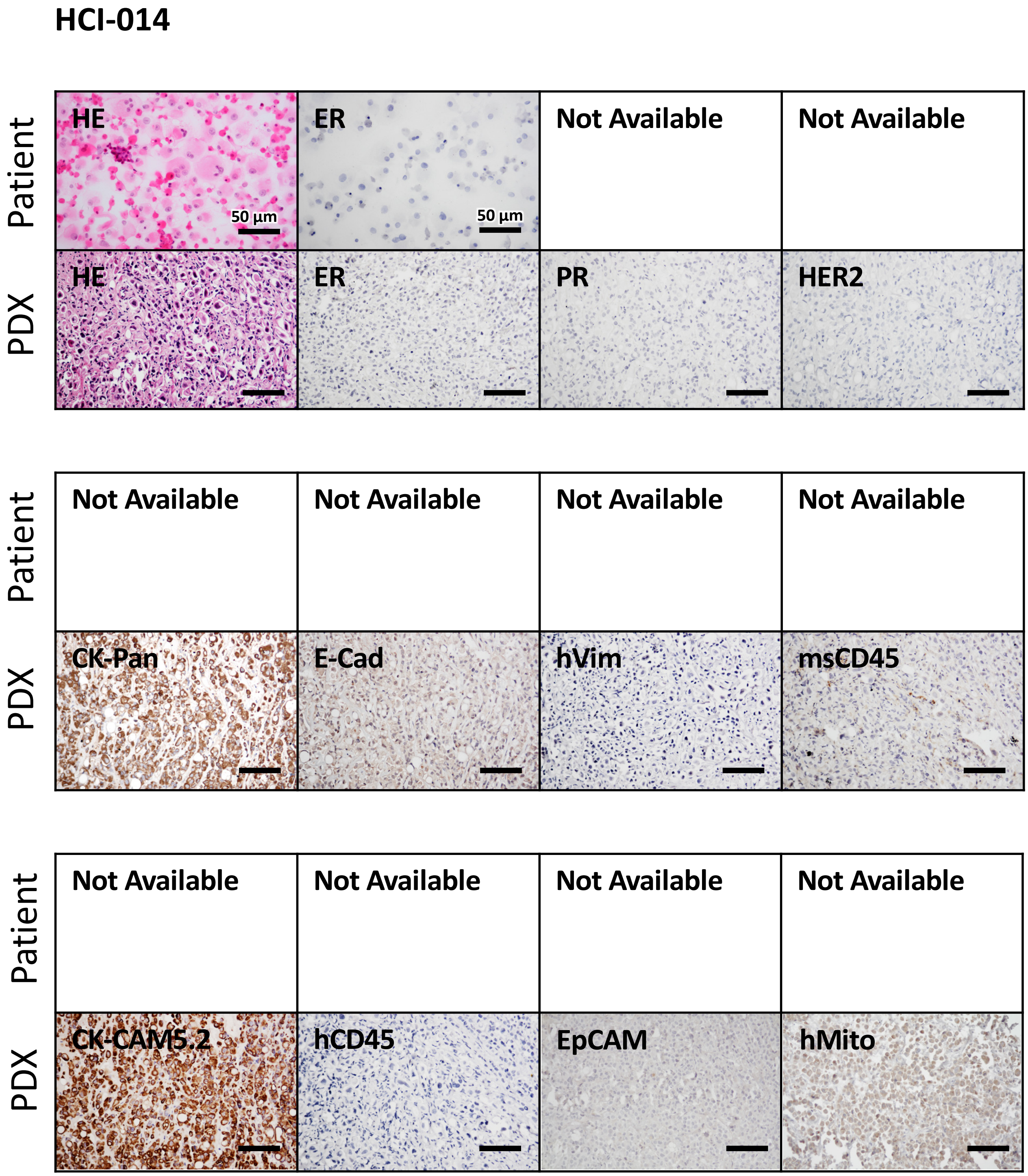


Scale bars = 100 µm, unless indicated otherwise

Supplementary Fig. 6

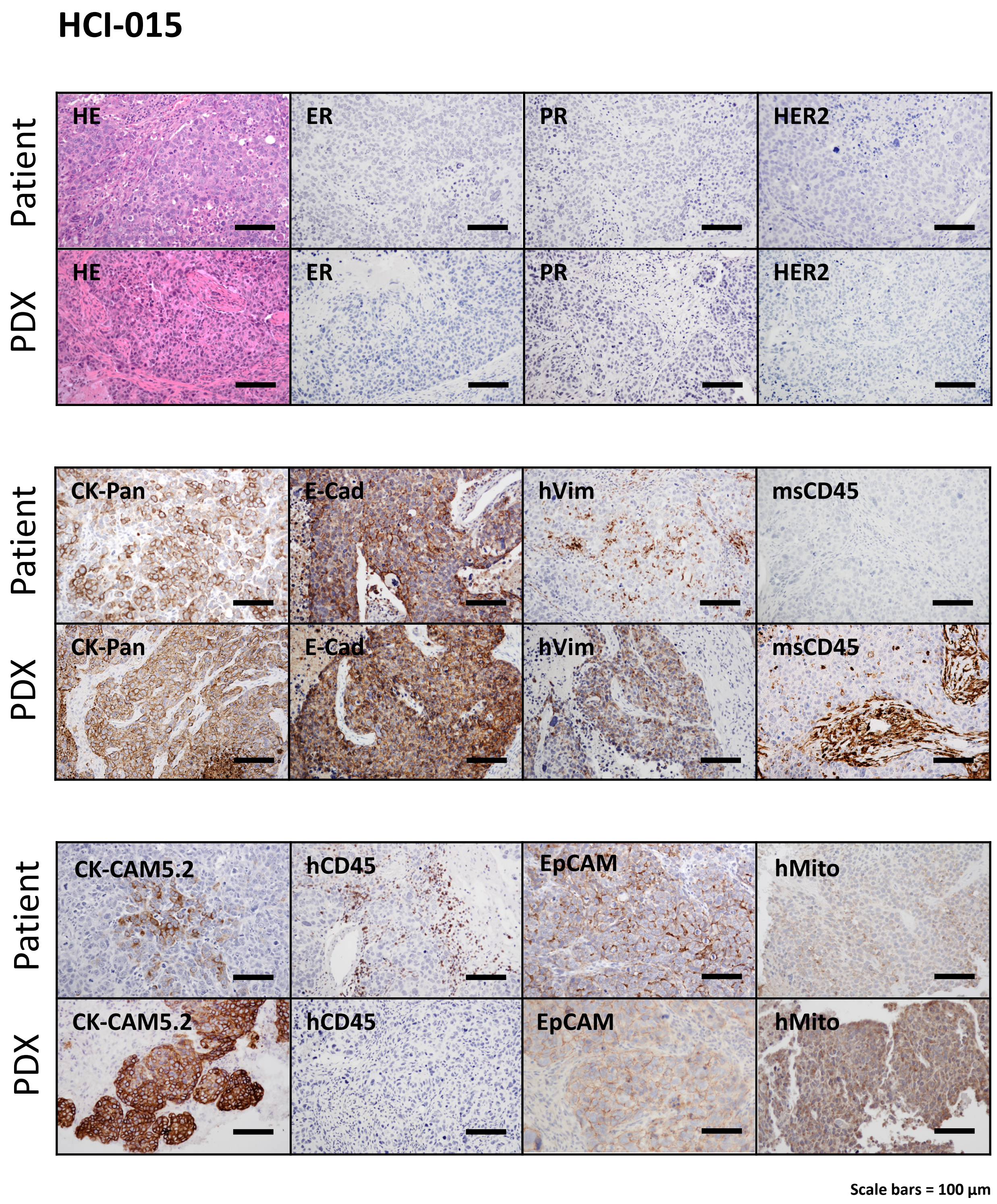

Supplementary Fig. 7

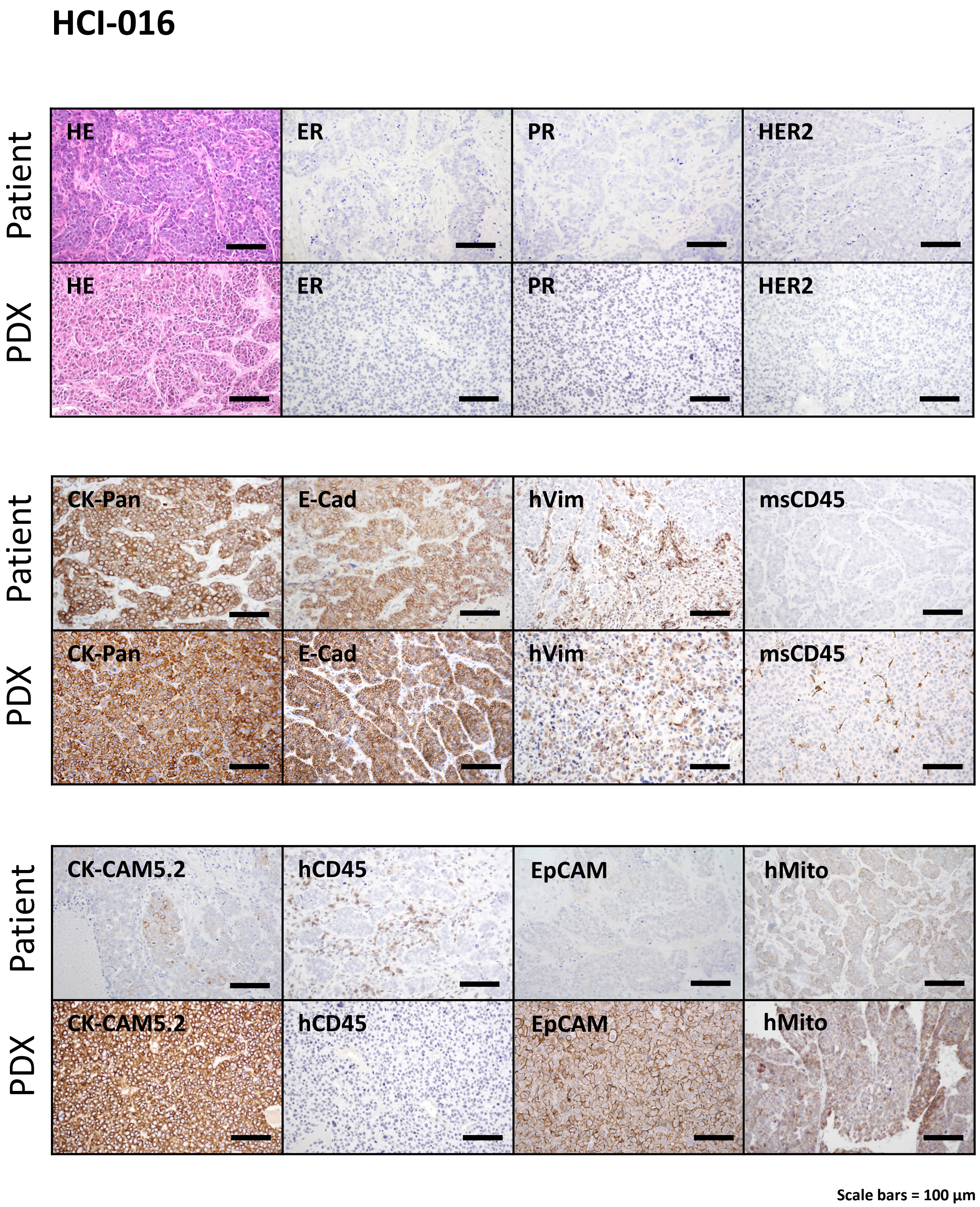

Supplementary Fig. 8

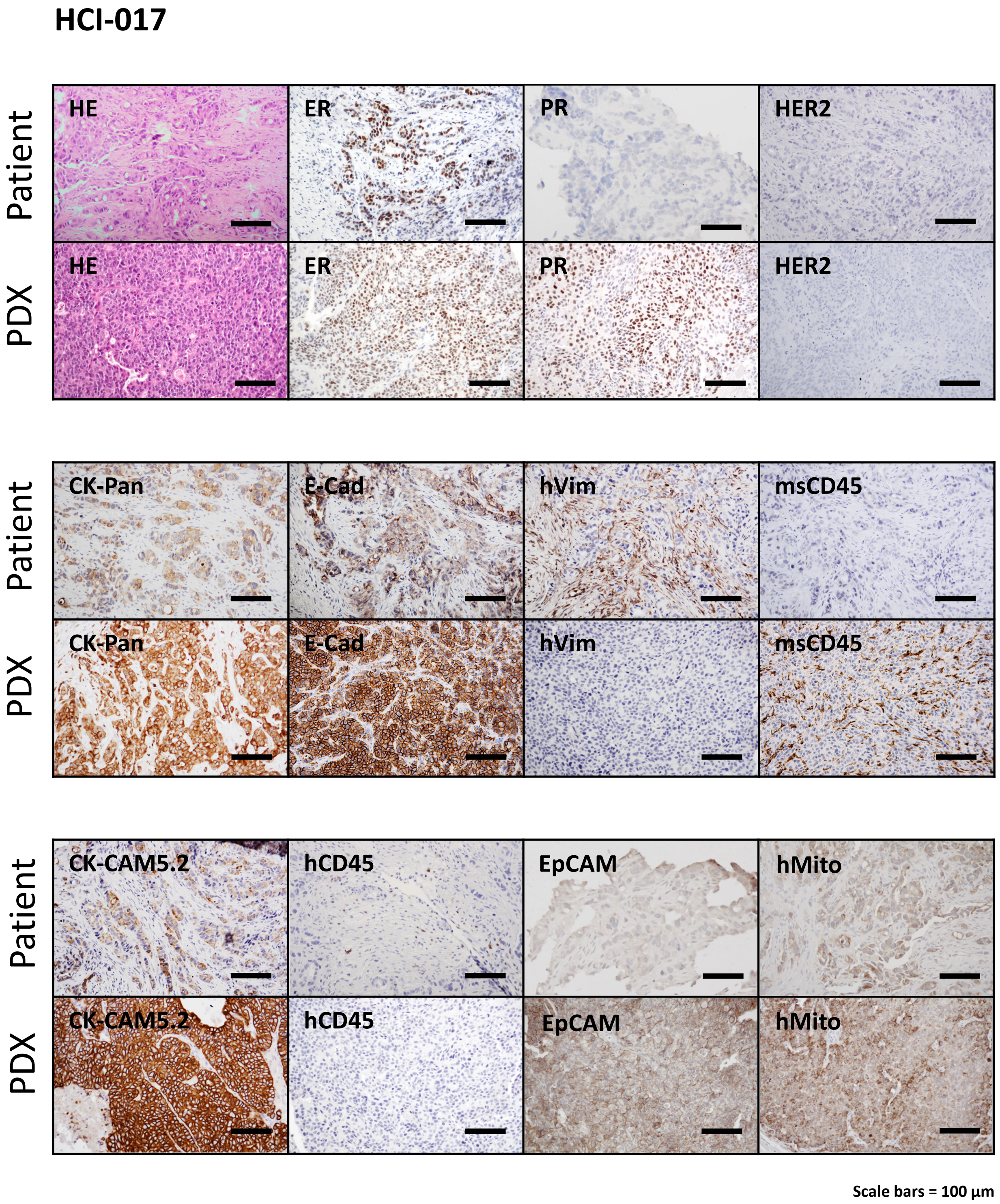

Supplementary Fig. 9

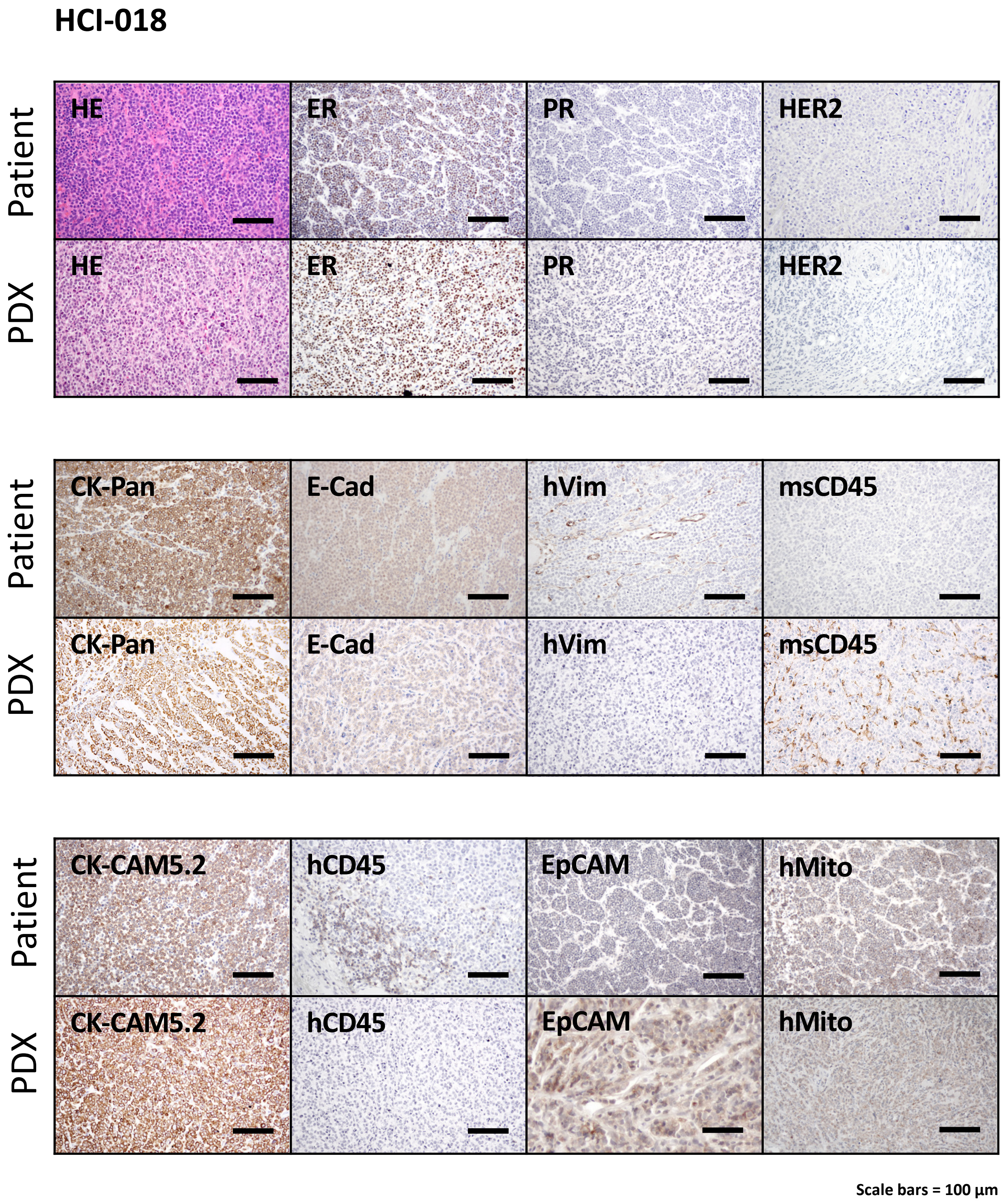

Supplementary Fig. 10

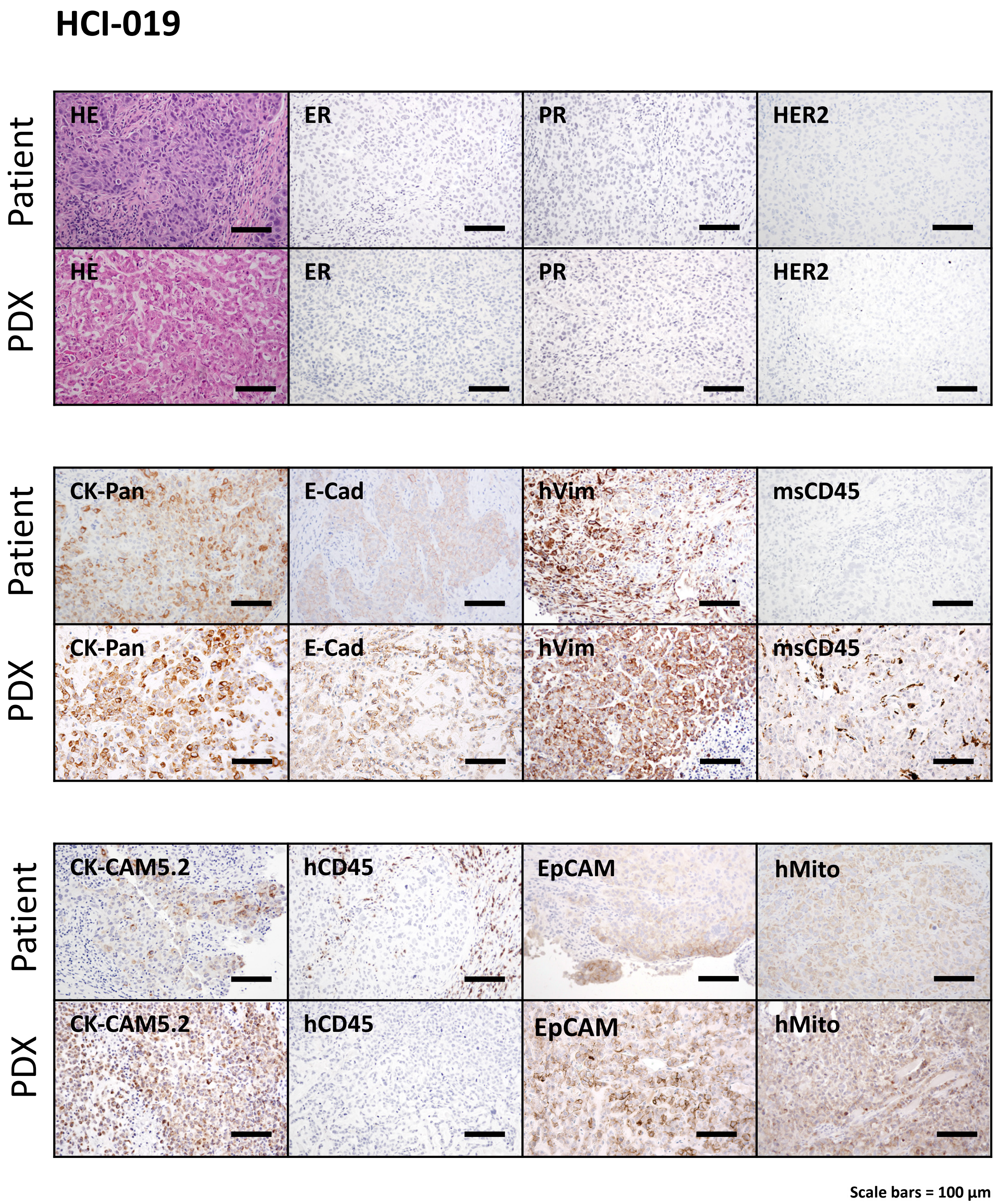

Supplementary Fig. 11

### HCI-023

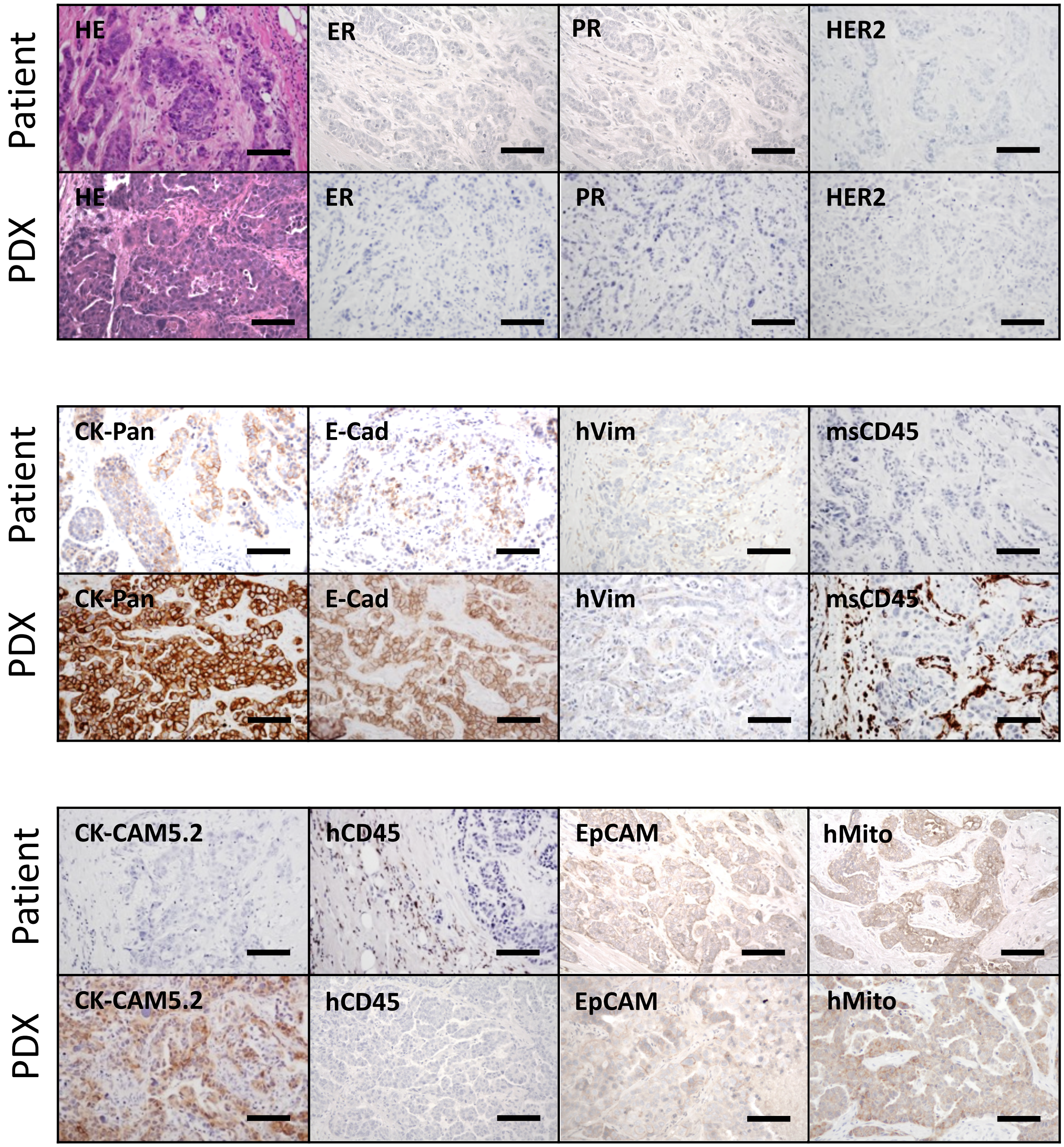


Scale bars = 100 µm

Supplementary Fig. 12

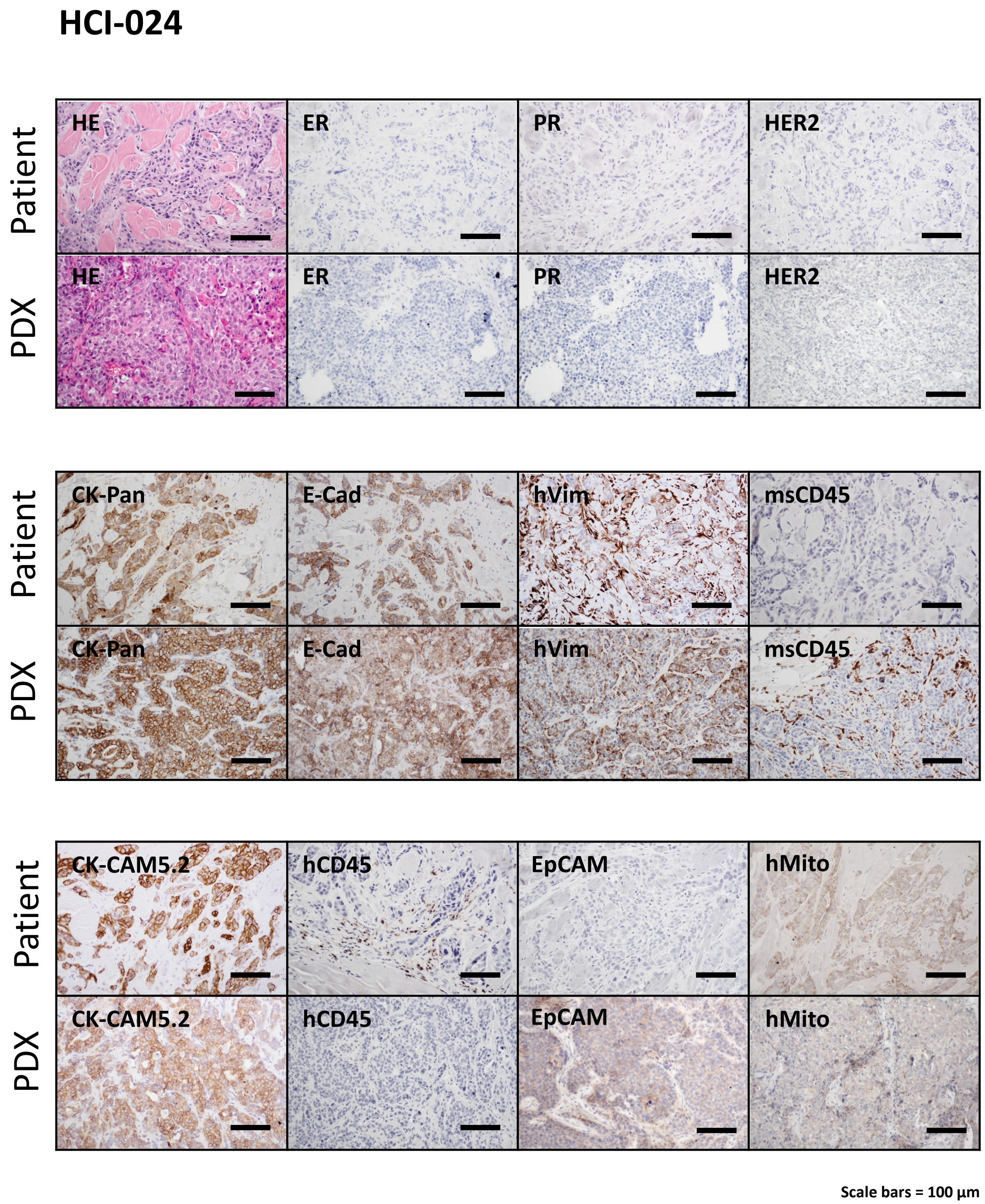

Supplementary Fig. 13

### HCI-025

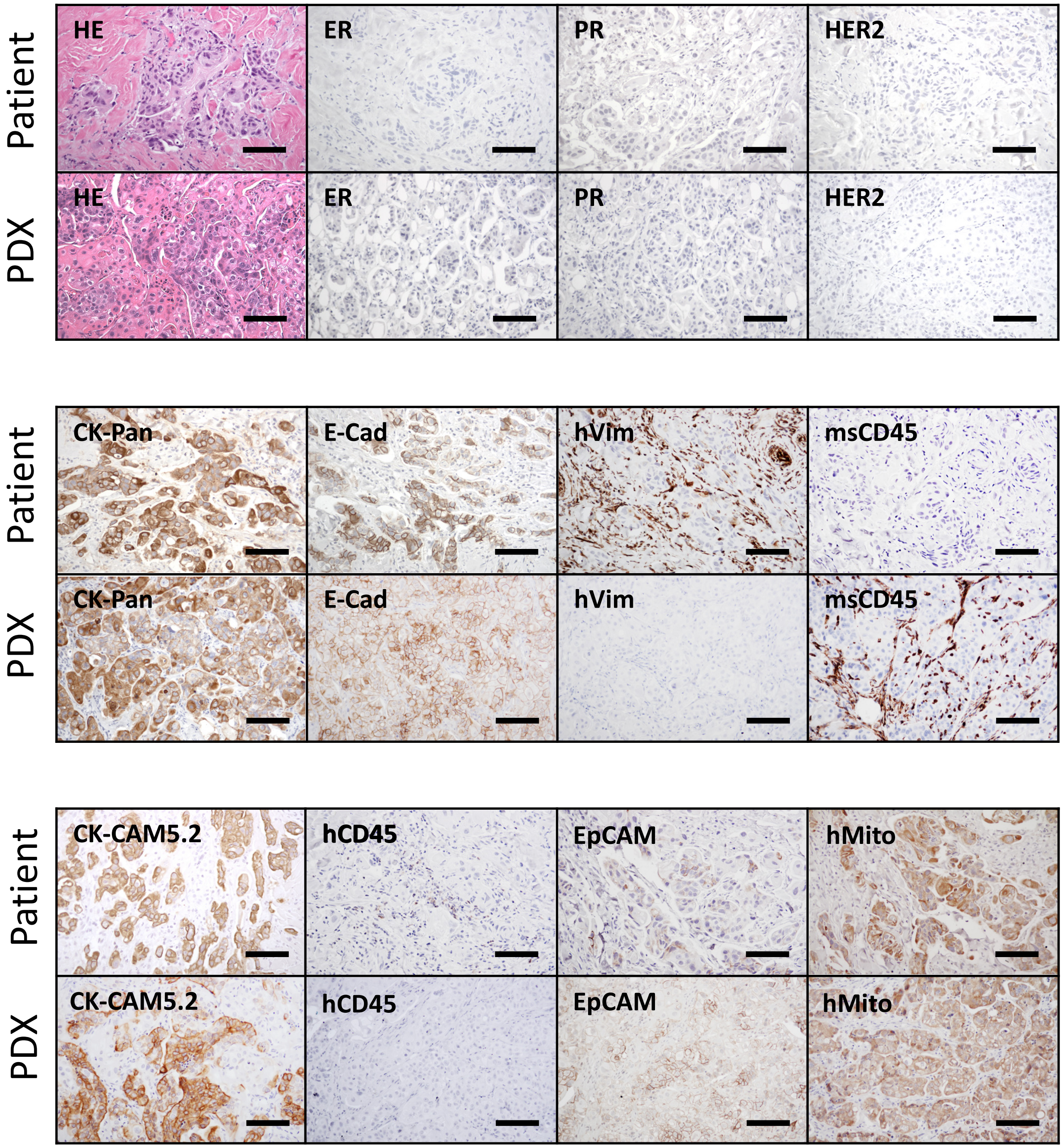


Scale bars = 100 µm

Supplementary Fig. 14

### HCI-026

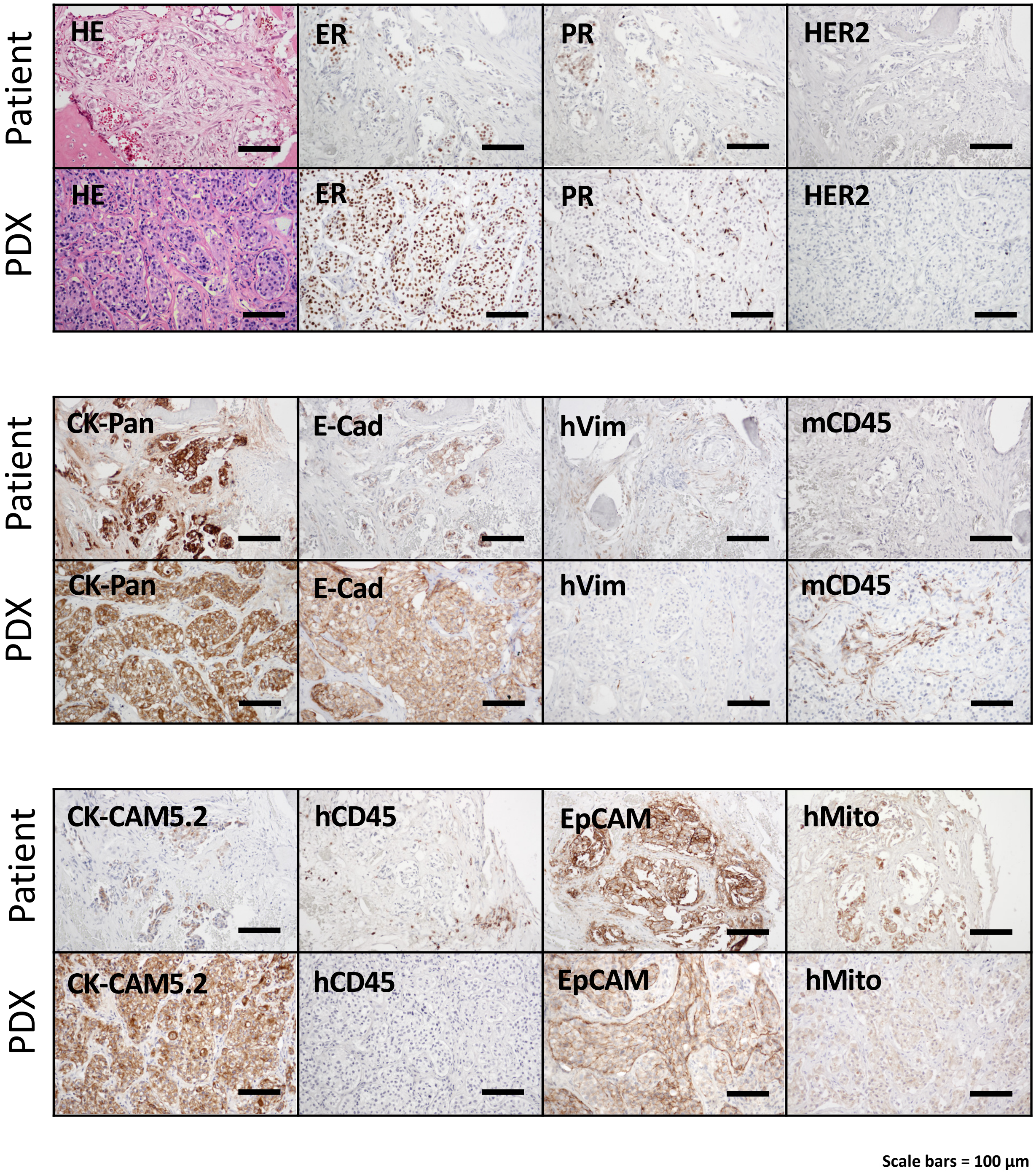

Supplementary Fig. 15

### HCI-027

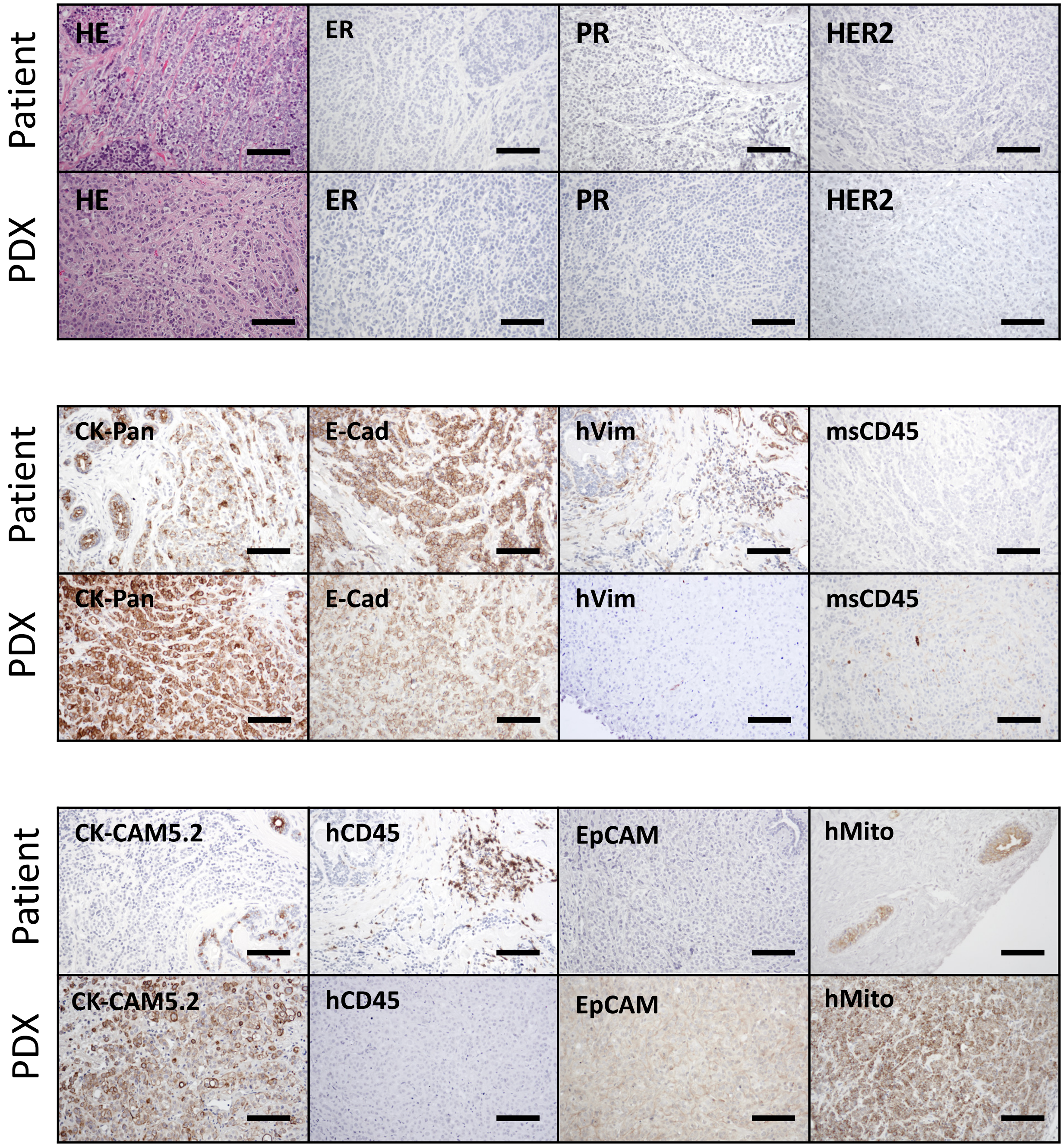


Scale bars = 100 µm

Supplementary Fig. 16

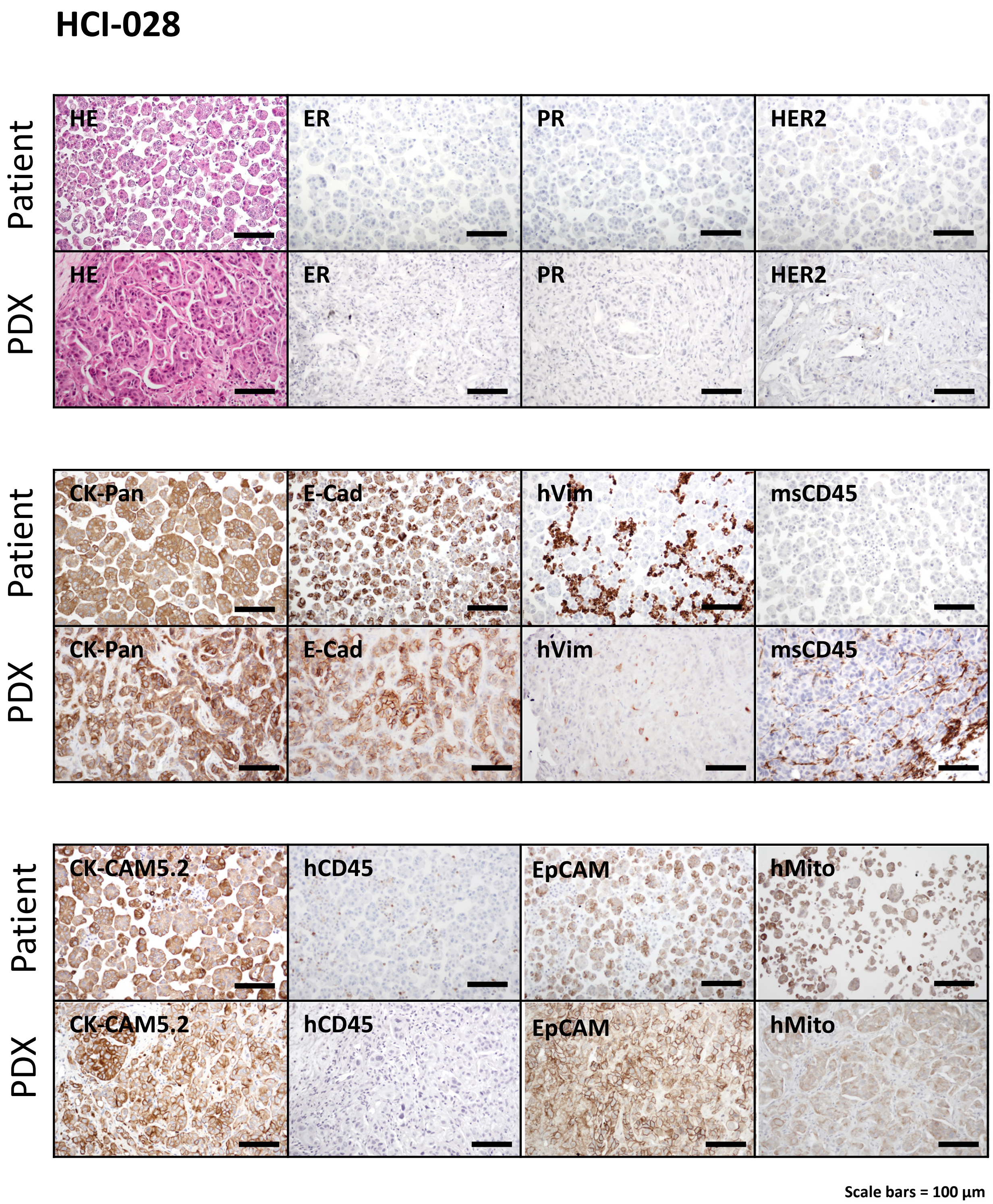

Supplementary Fig. 17

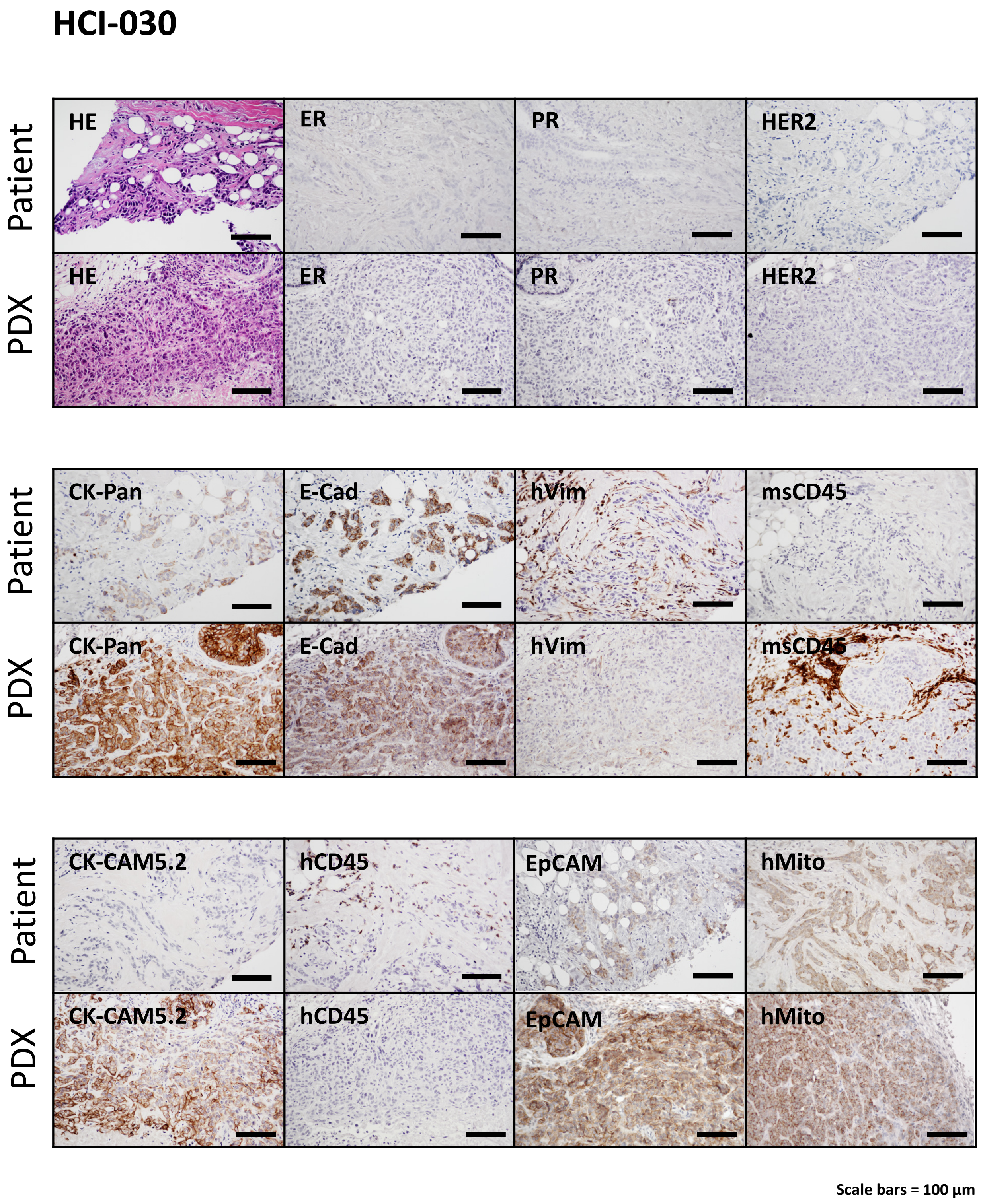

Supplementary Fig. 18

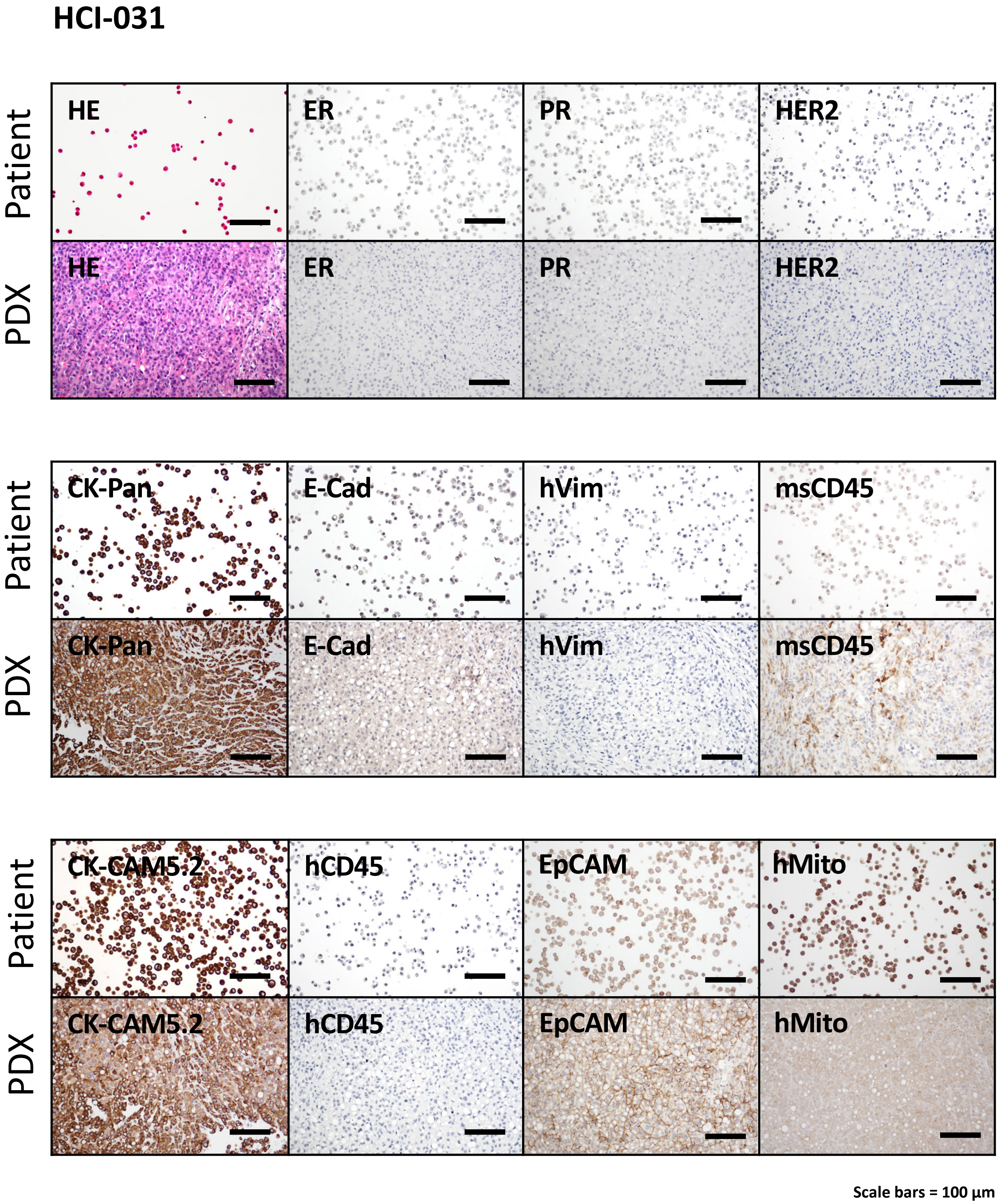

Supplementary Fig. 19

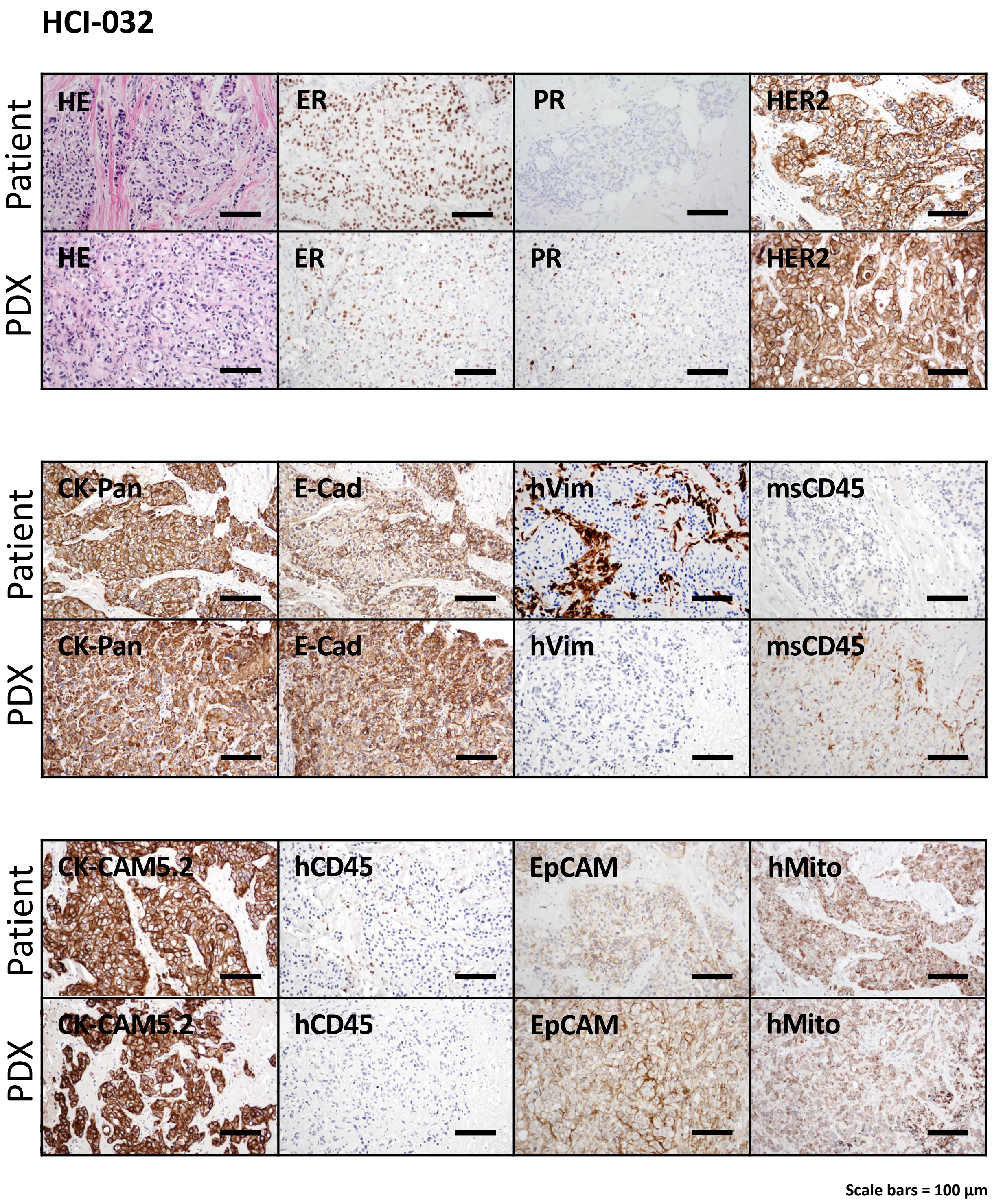

Supplementary Fig. 20

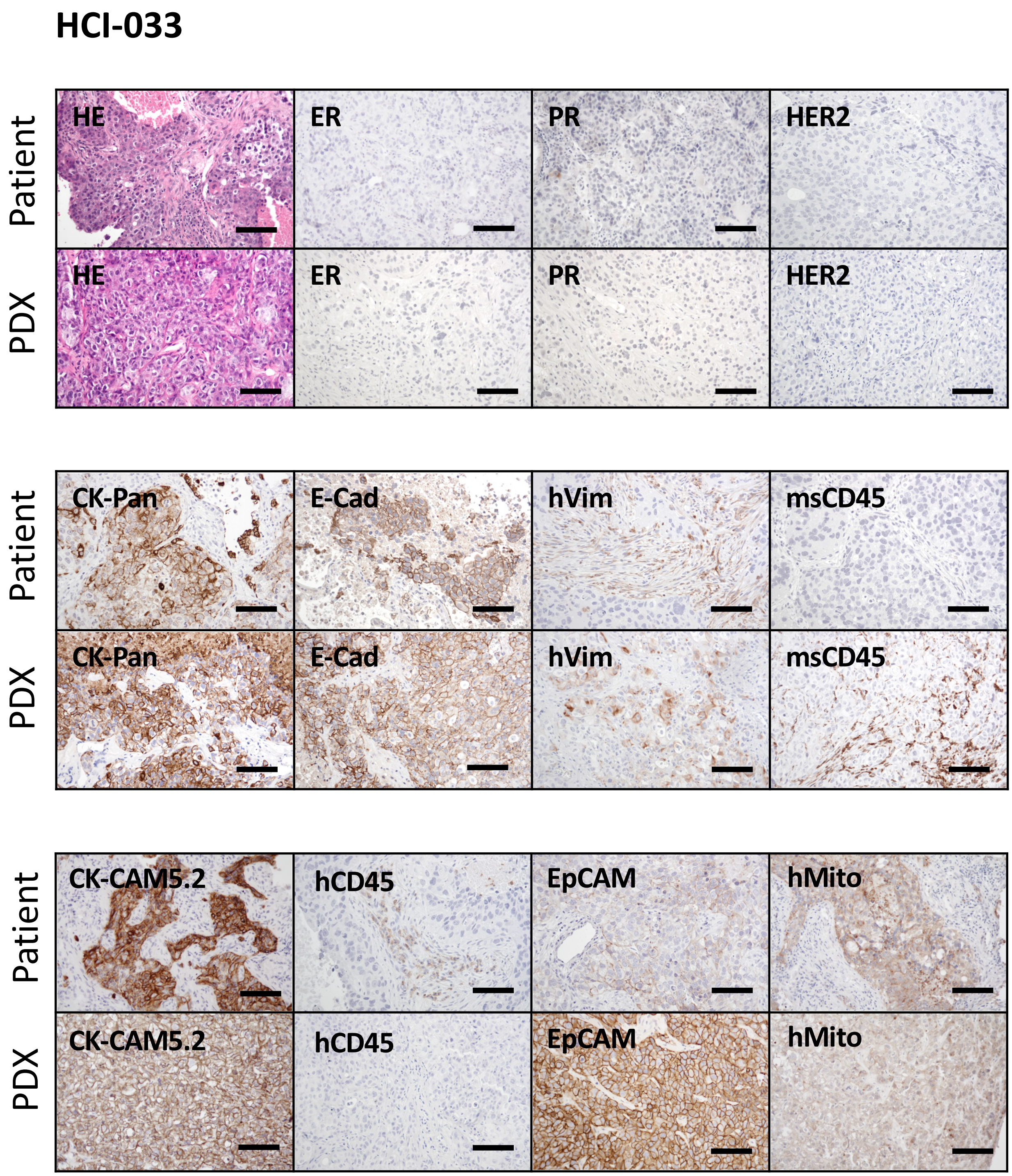


Scale bars = 100 µm

Supplementary Fig. 21

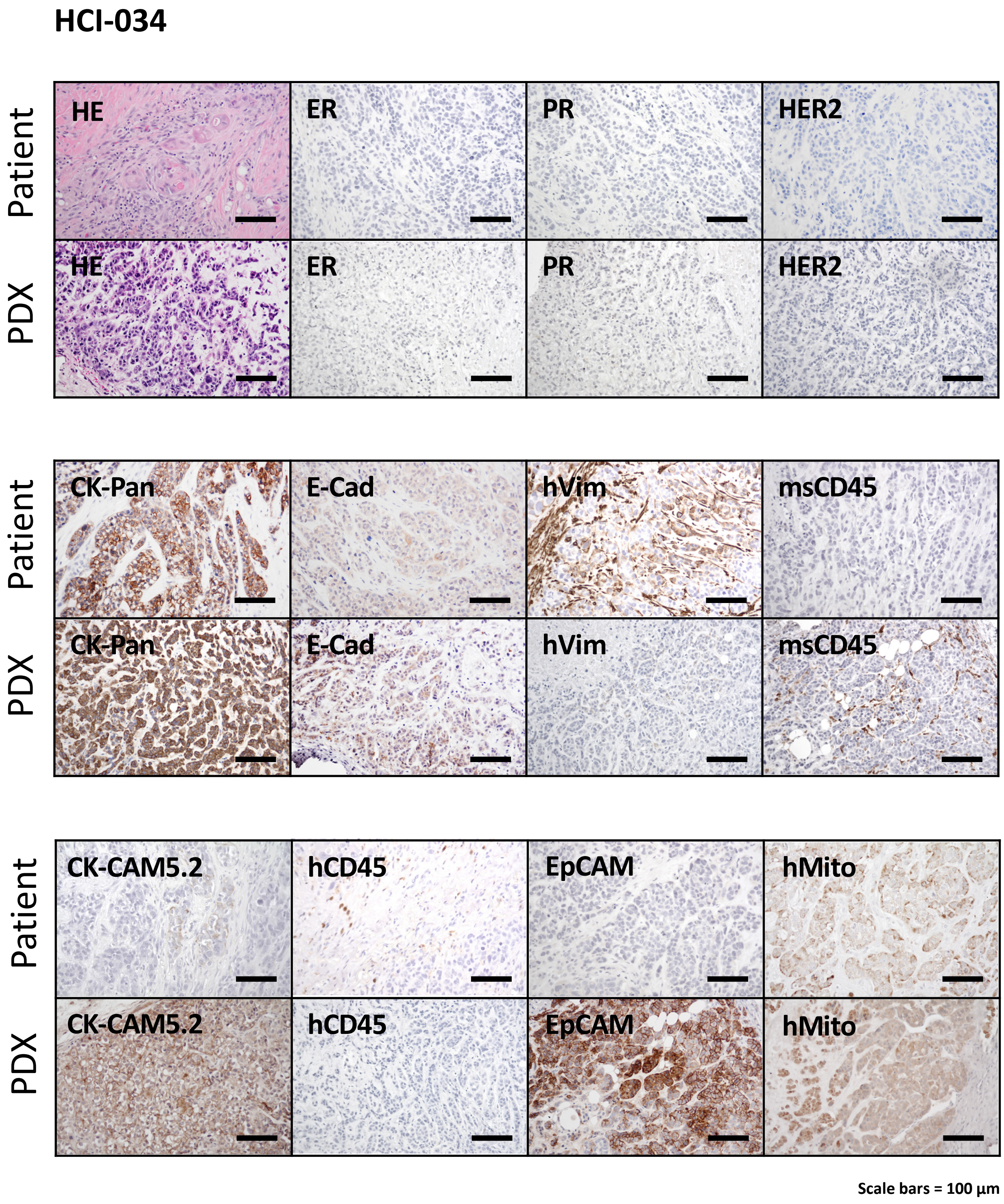

Supplementary Fig. 22

#### HCI-036

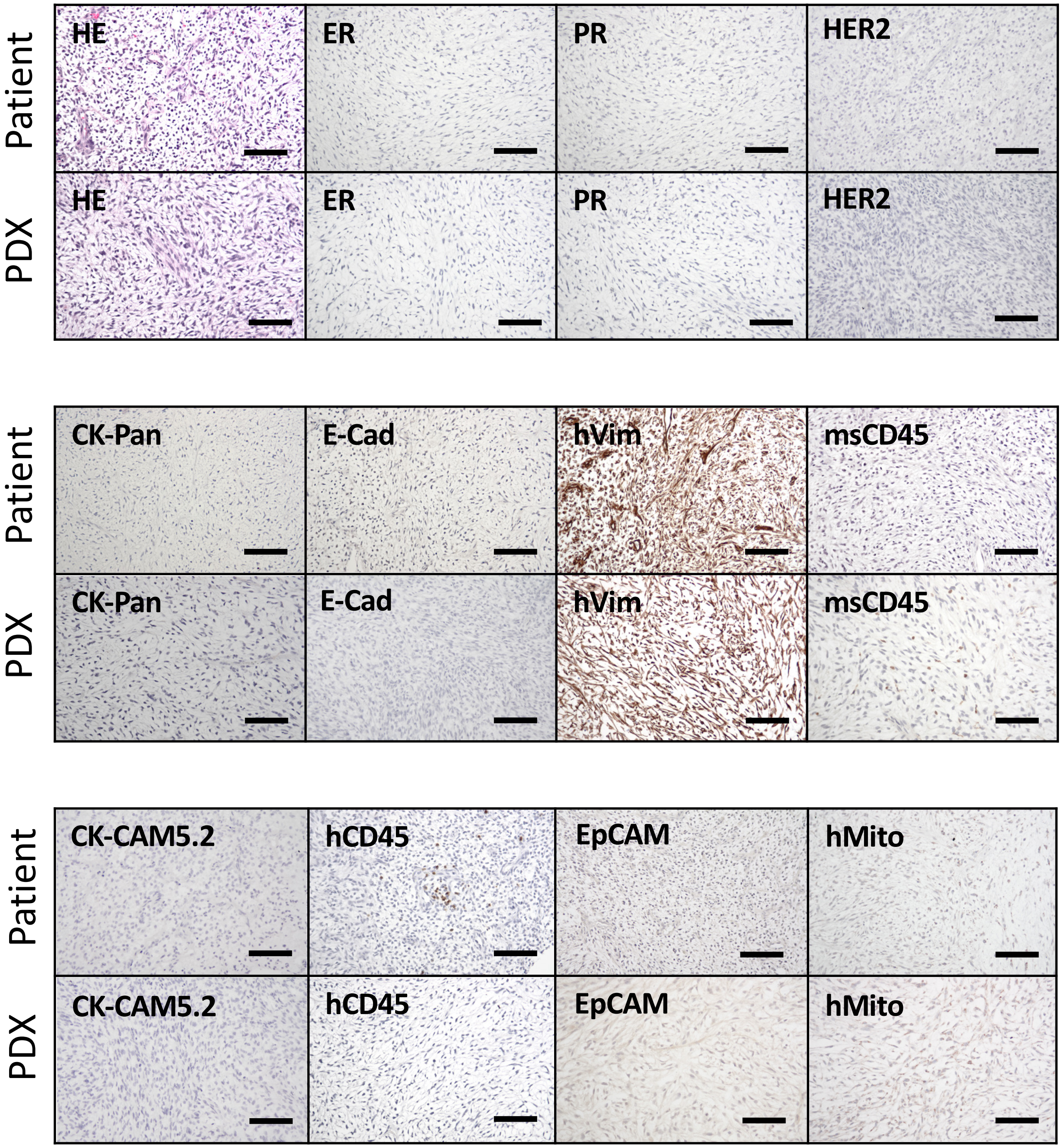


Scale bars = 100 µm

Supplementary Fig. 23

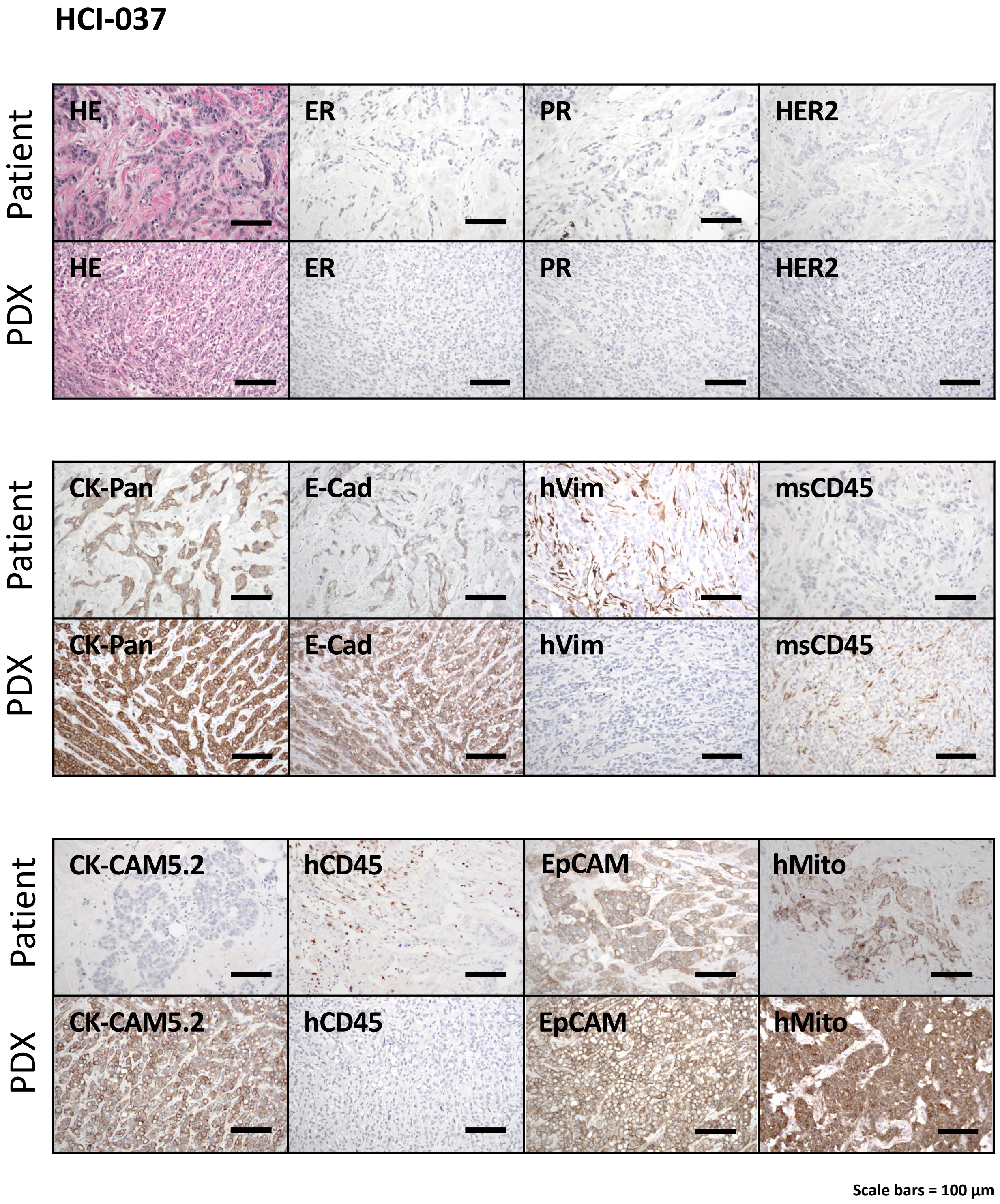

Supplementary Fig. 24

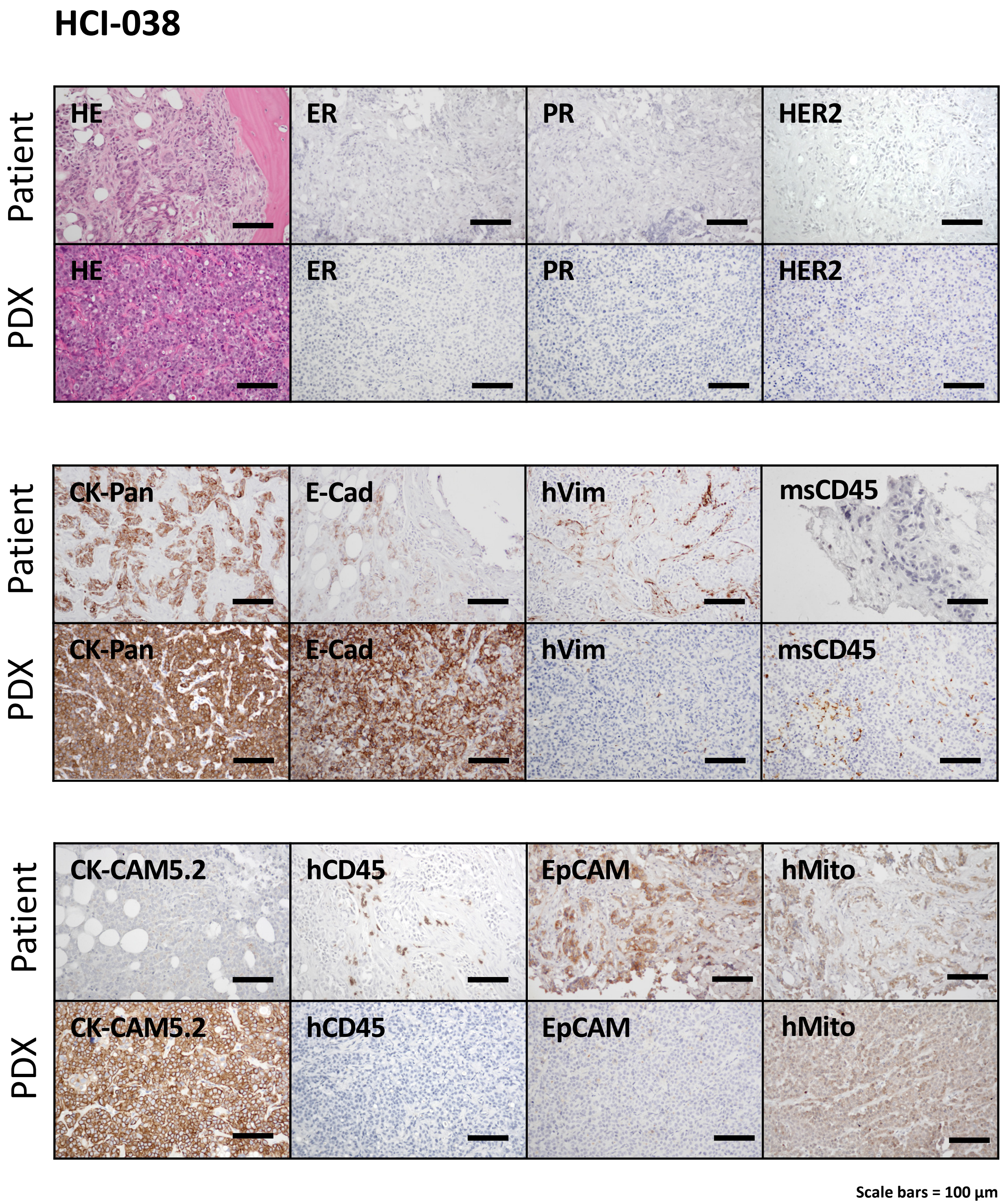

Supplementary Fig. 25

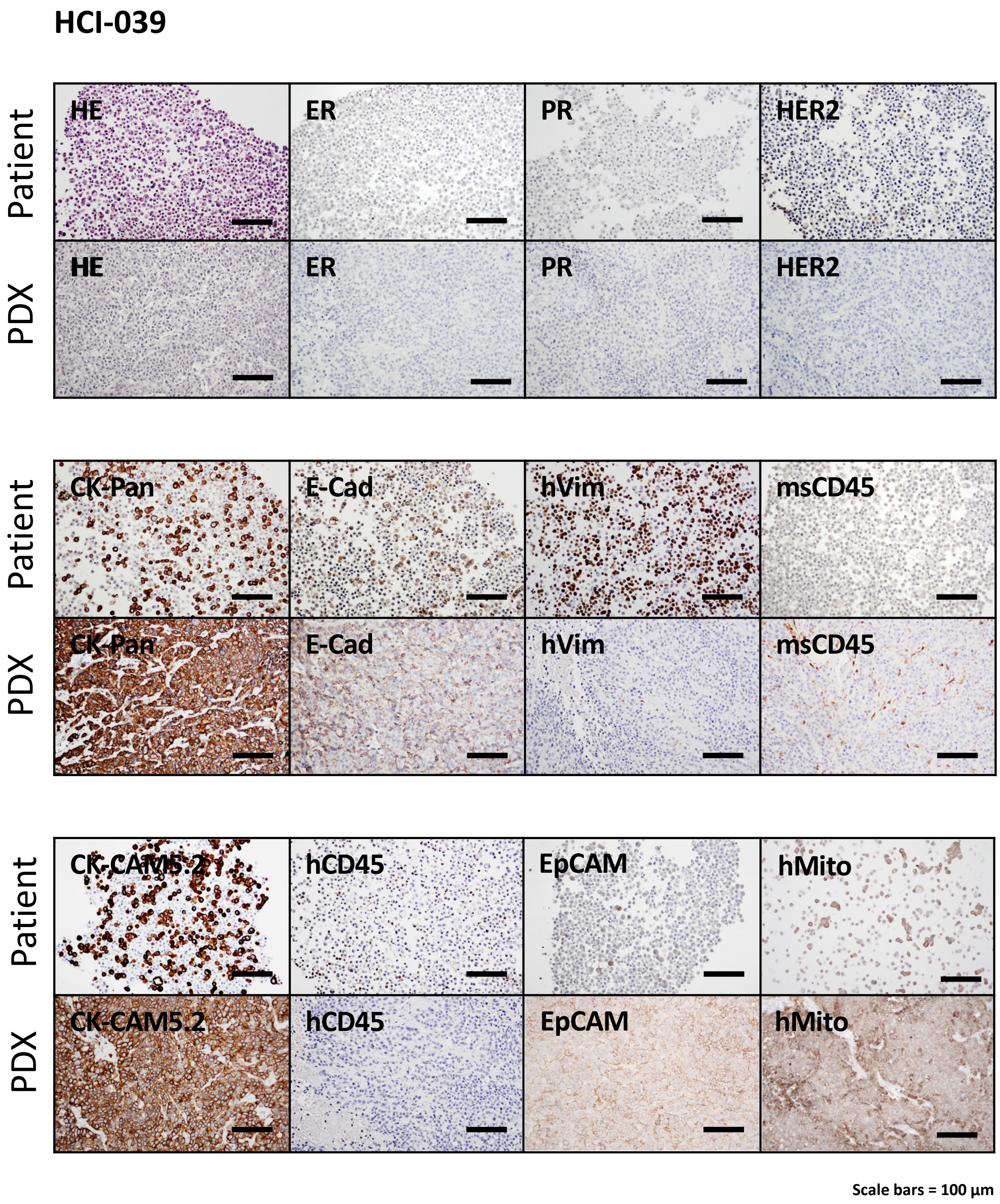

Supplementary Fig. 26

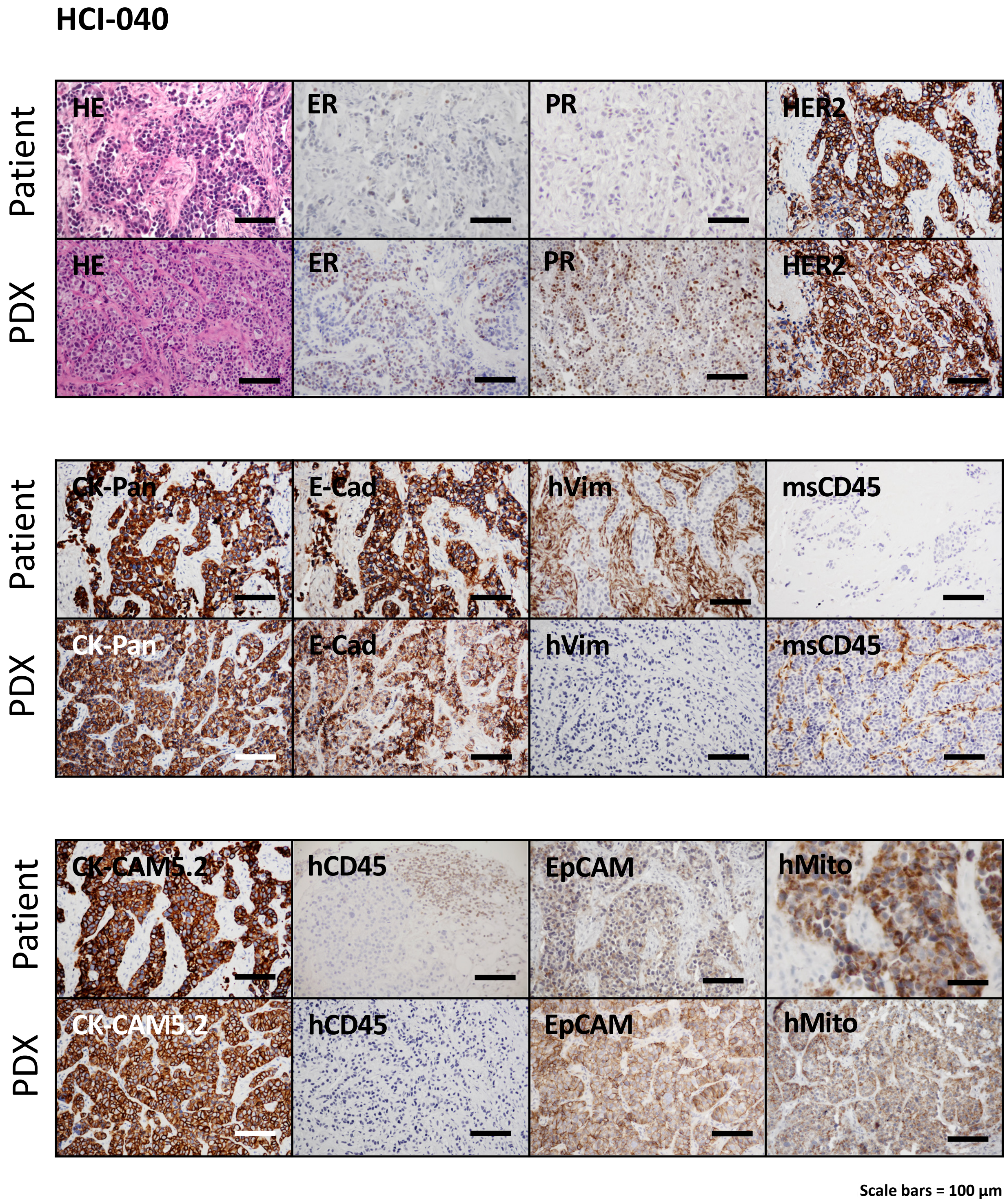

Supplementary Fig. 27

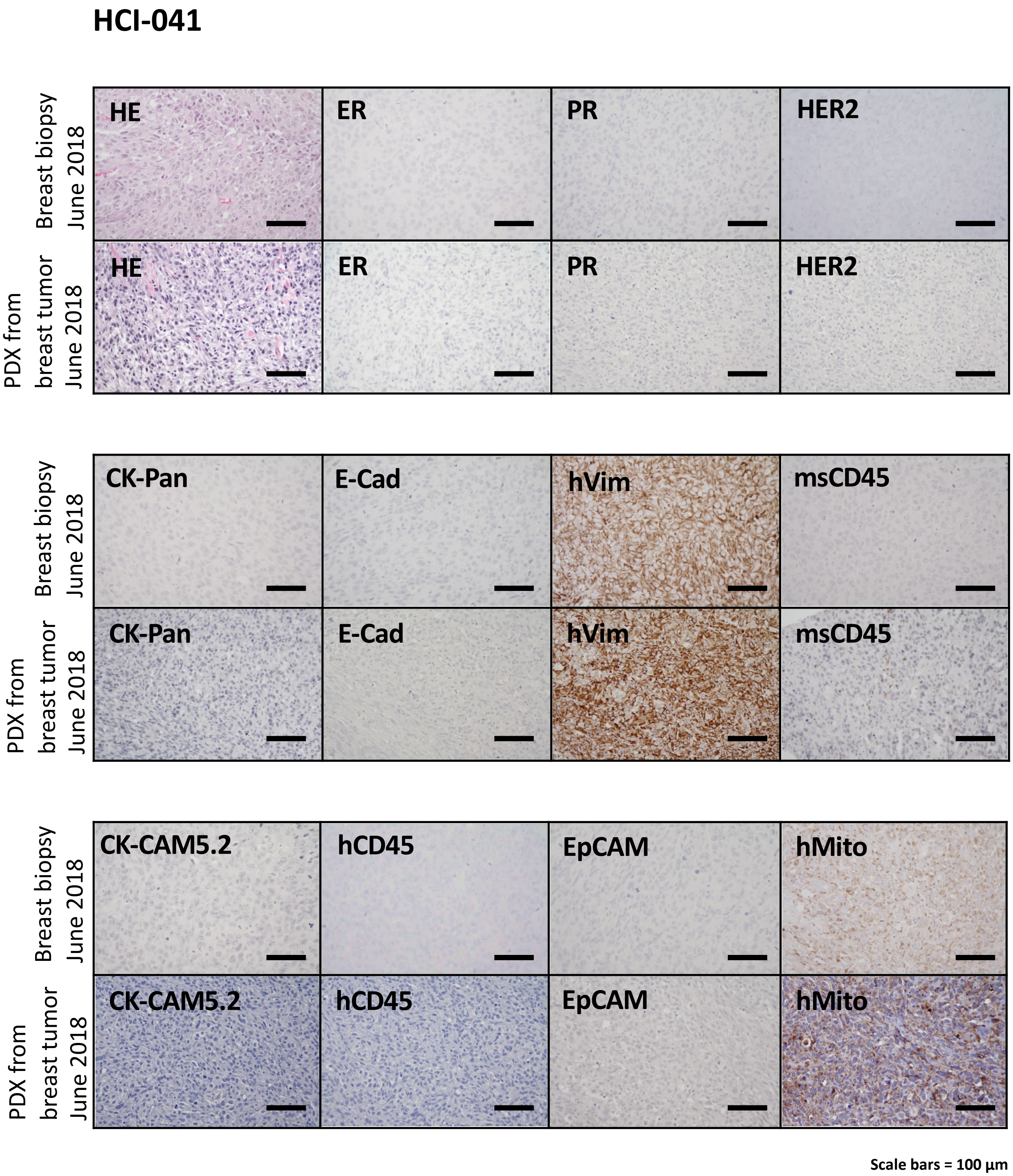
