## Supplemental Figures part 2 for "A breast cancer patient-derived xenograft and organoid platform for drug discovery and precision oncology"

Supplementary Fig. 28

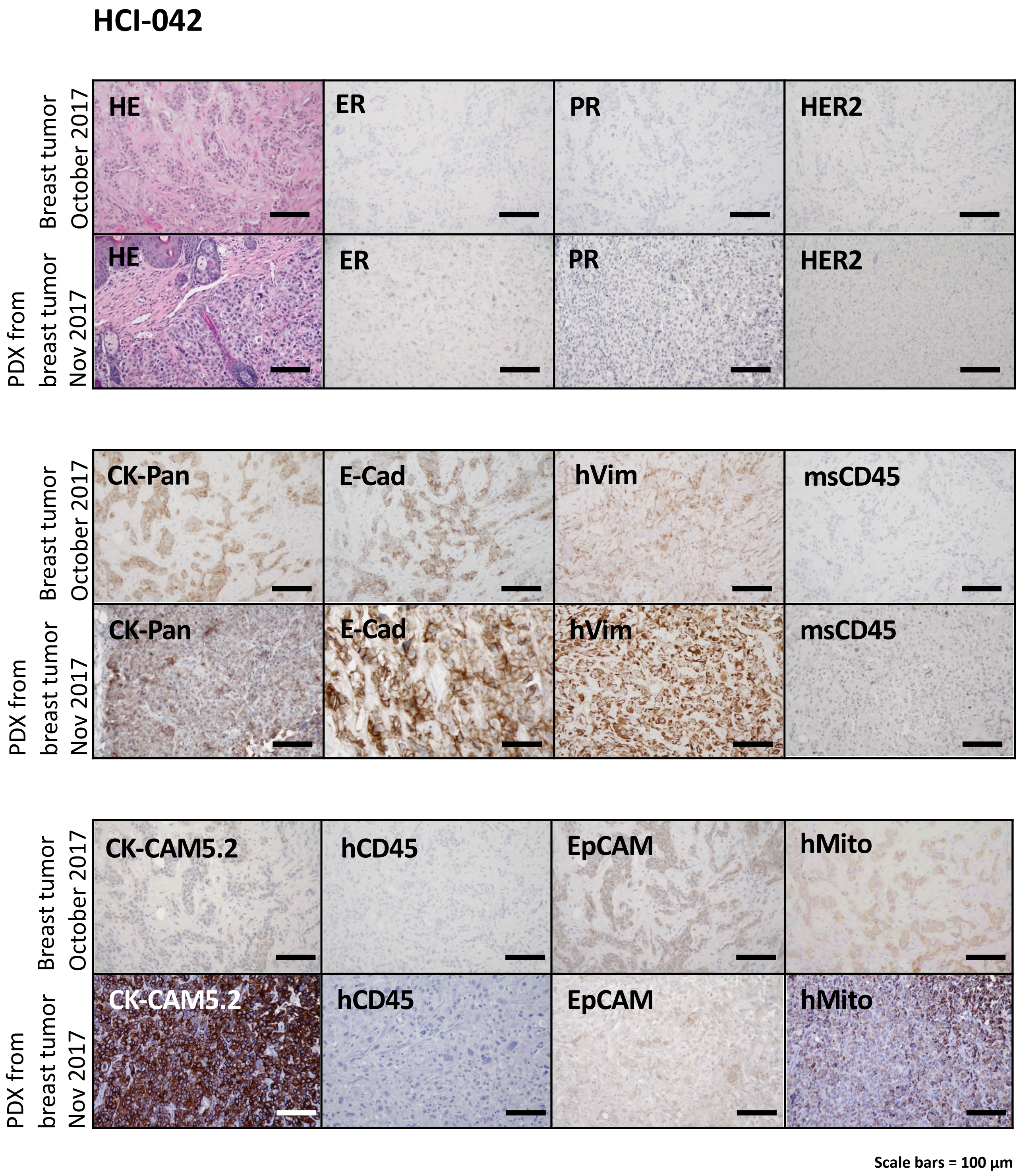

Supplementary Fig. 29

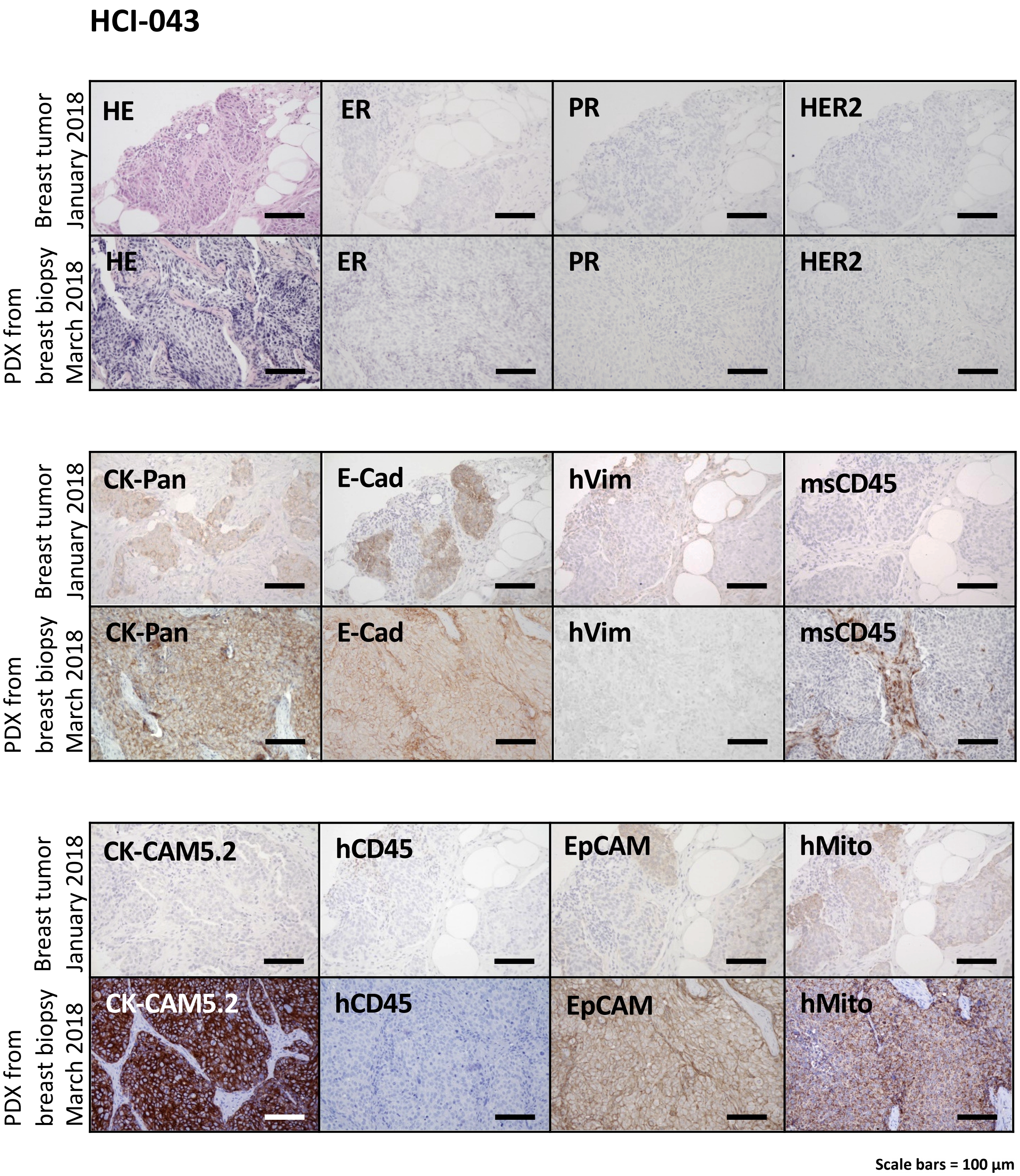

Supplementary Fig. 30

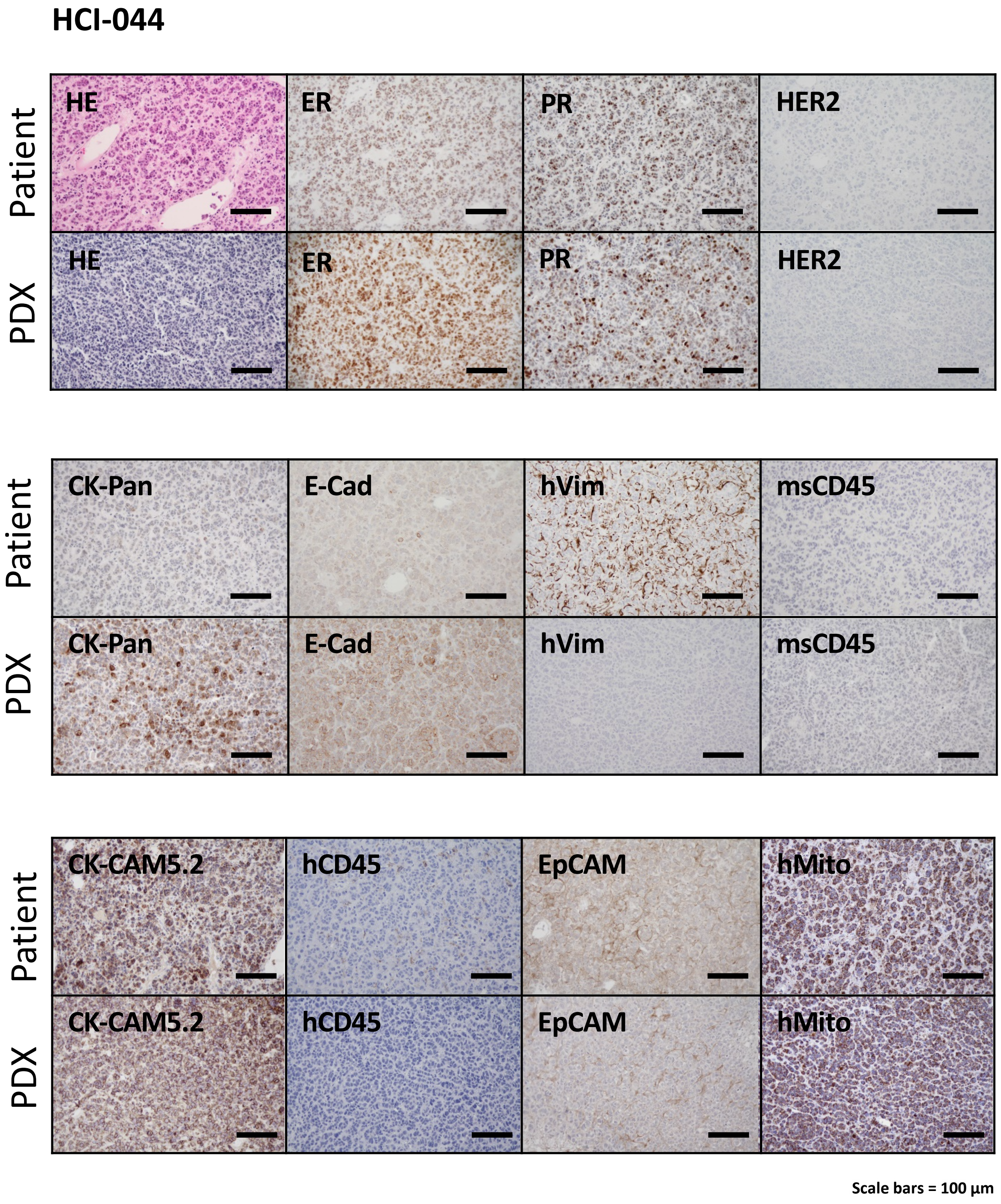

Supplementary Fig. 31

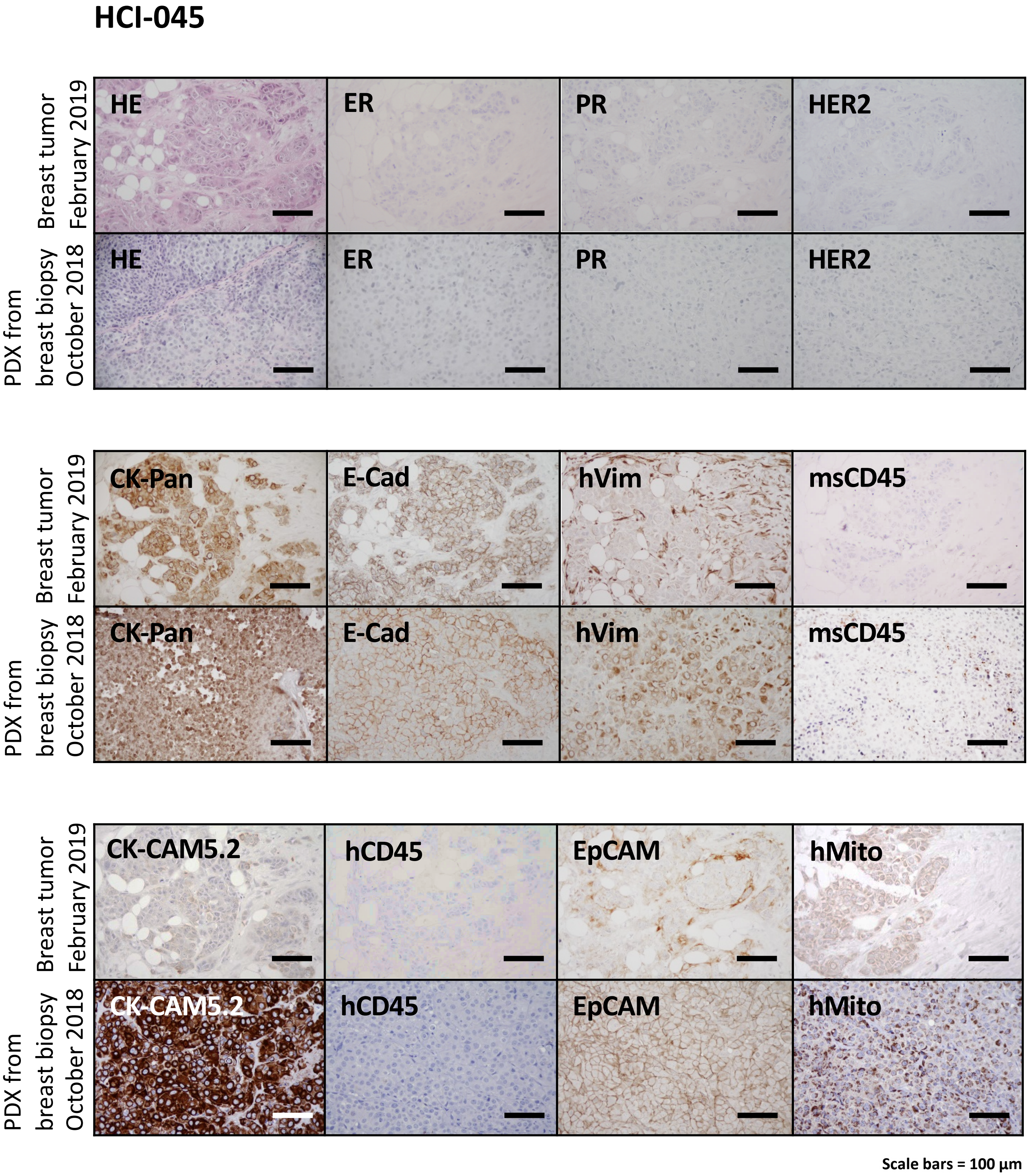

Supplementary Fig. 32

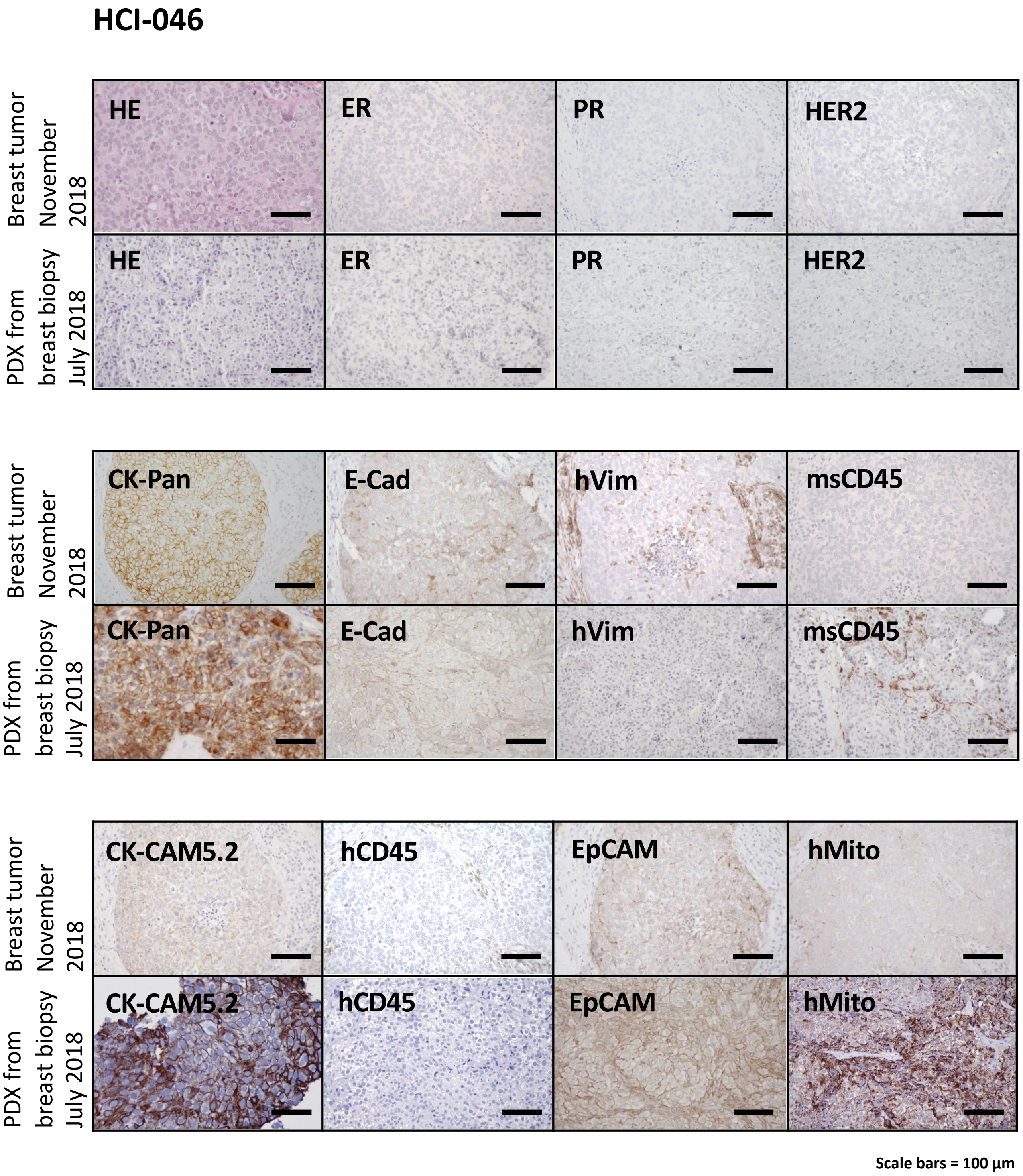

Supplementary Fig. 33

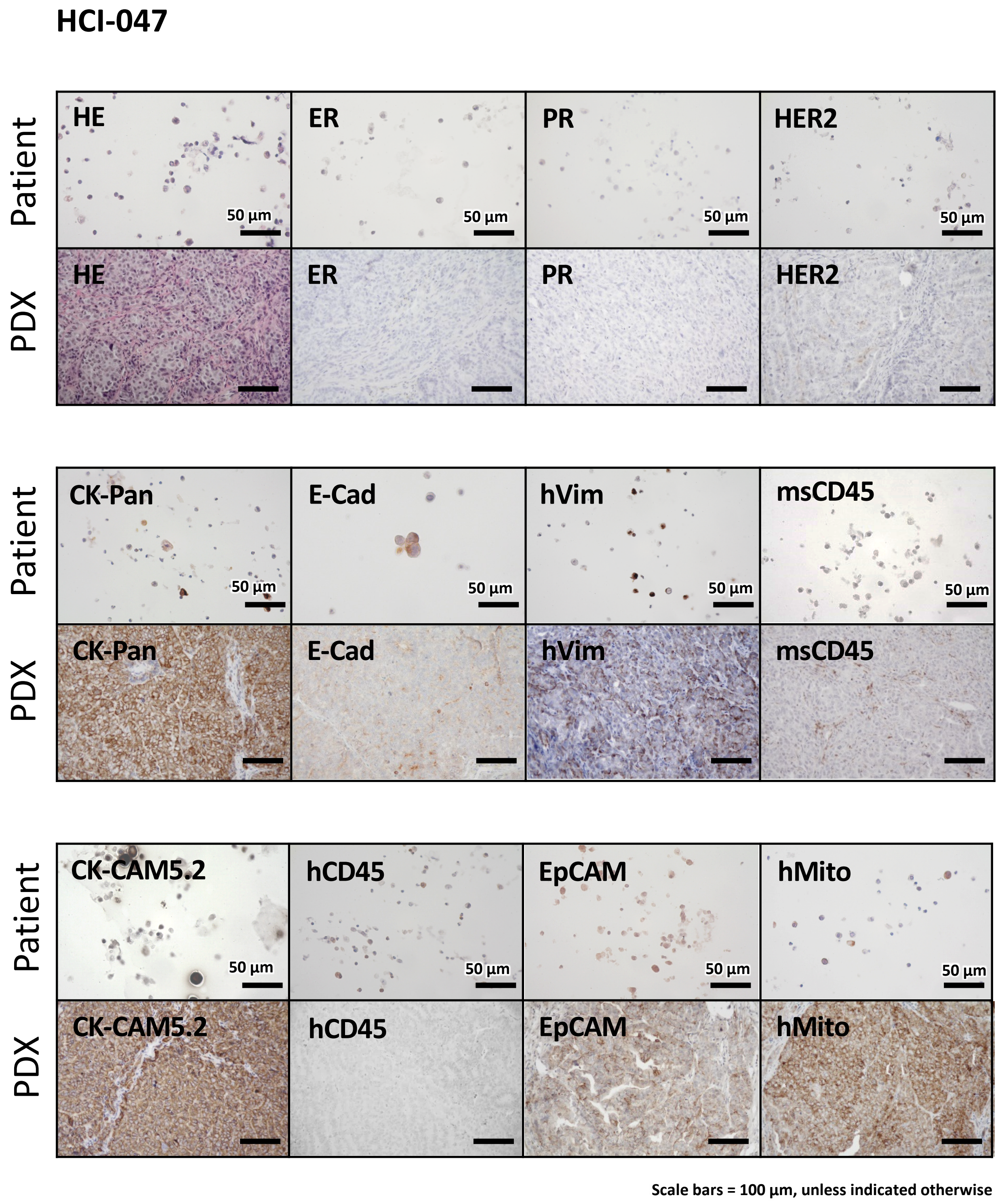

Supplementary Fig. 34

#### HCI-048

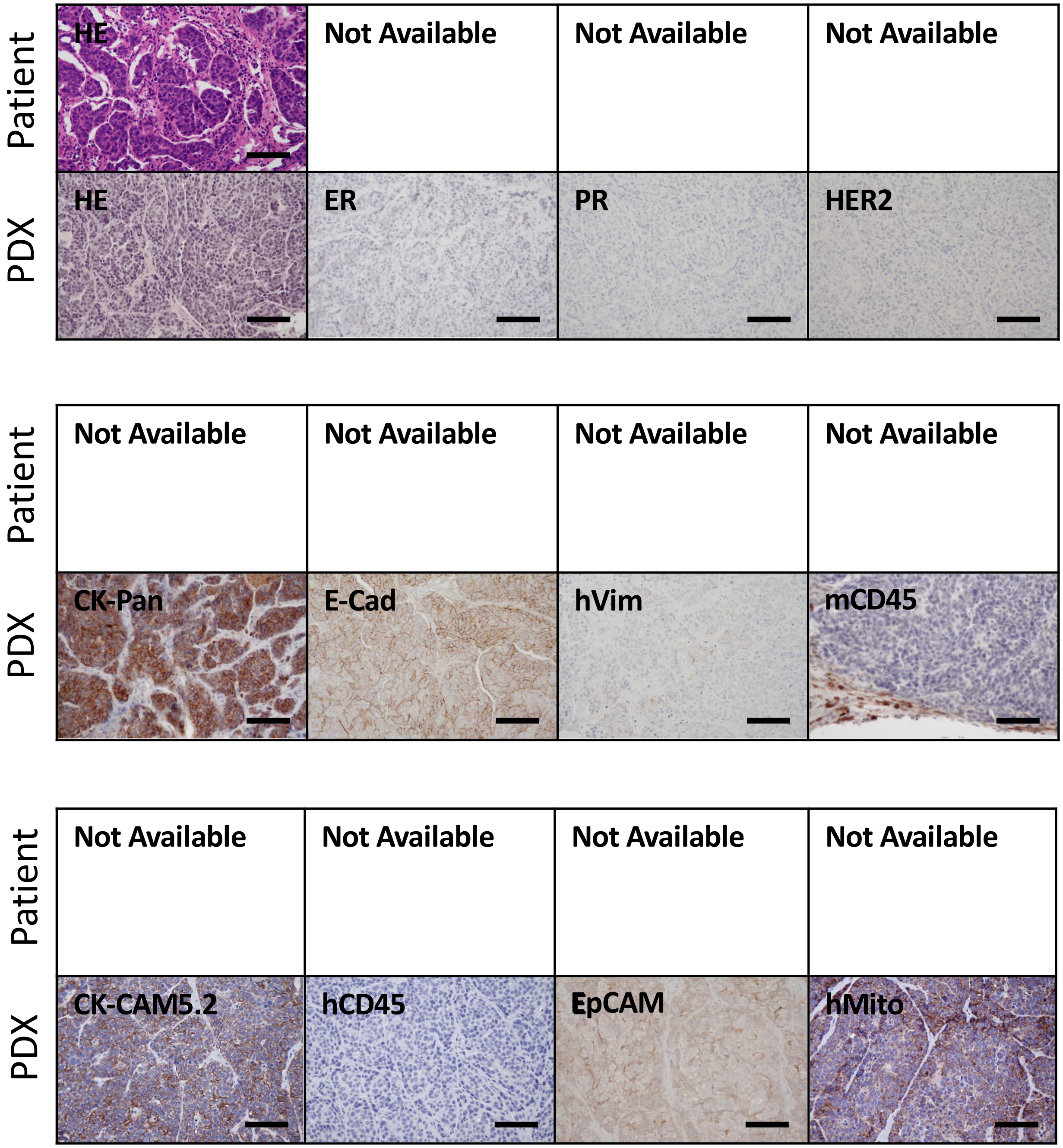


Scale bars = 100 µm

Supplementary Fig. 35

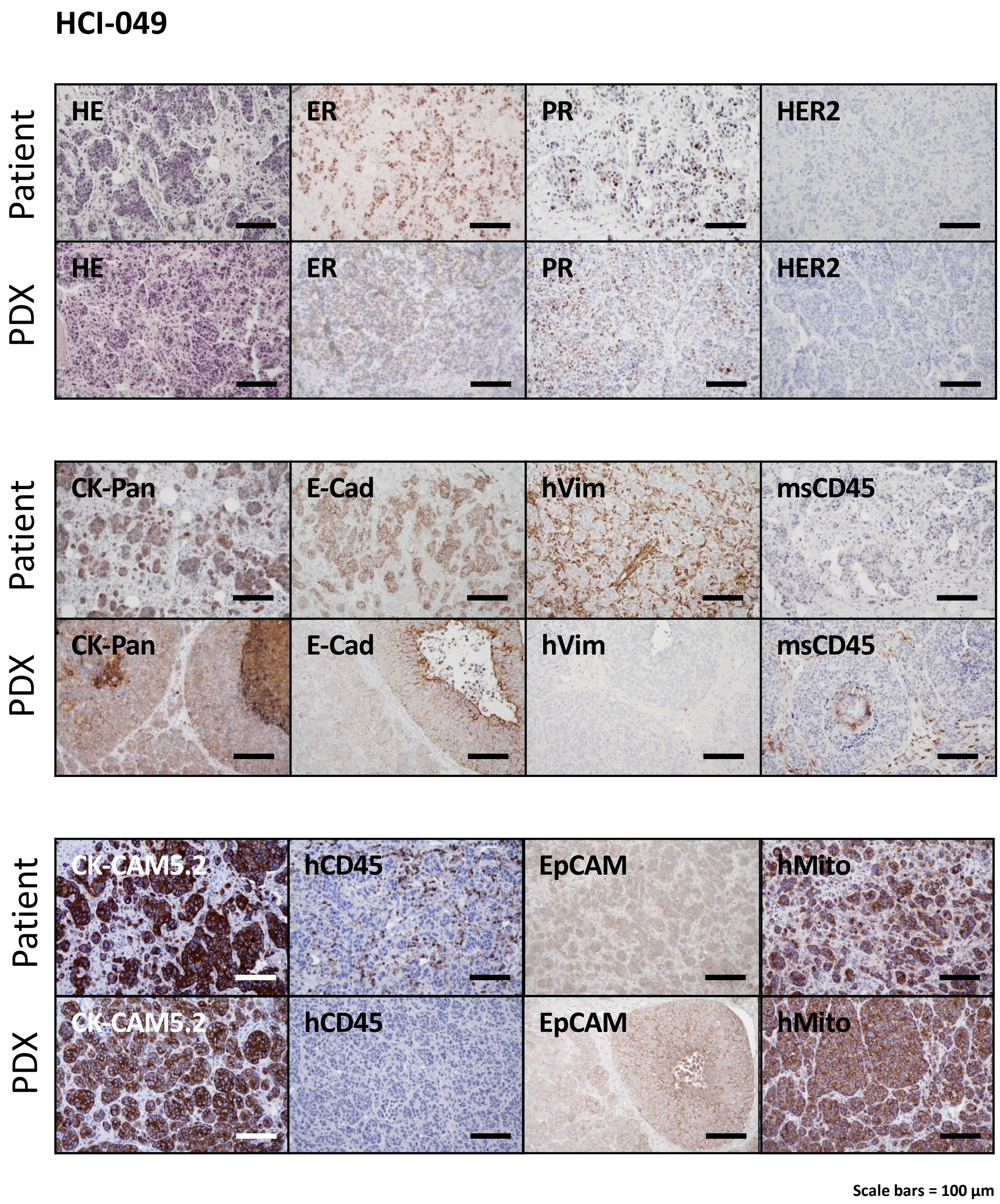

Supplementary Fig. 36

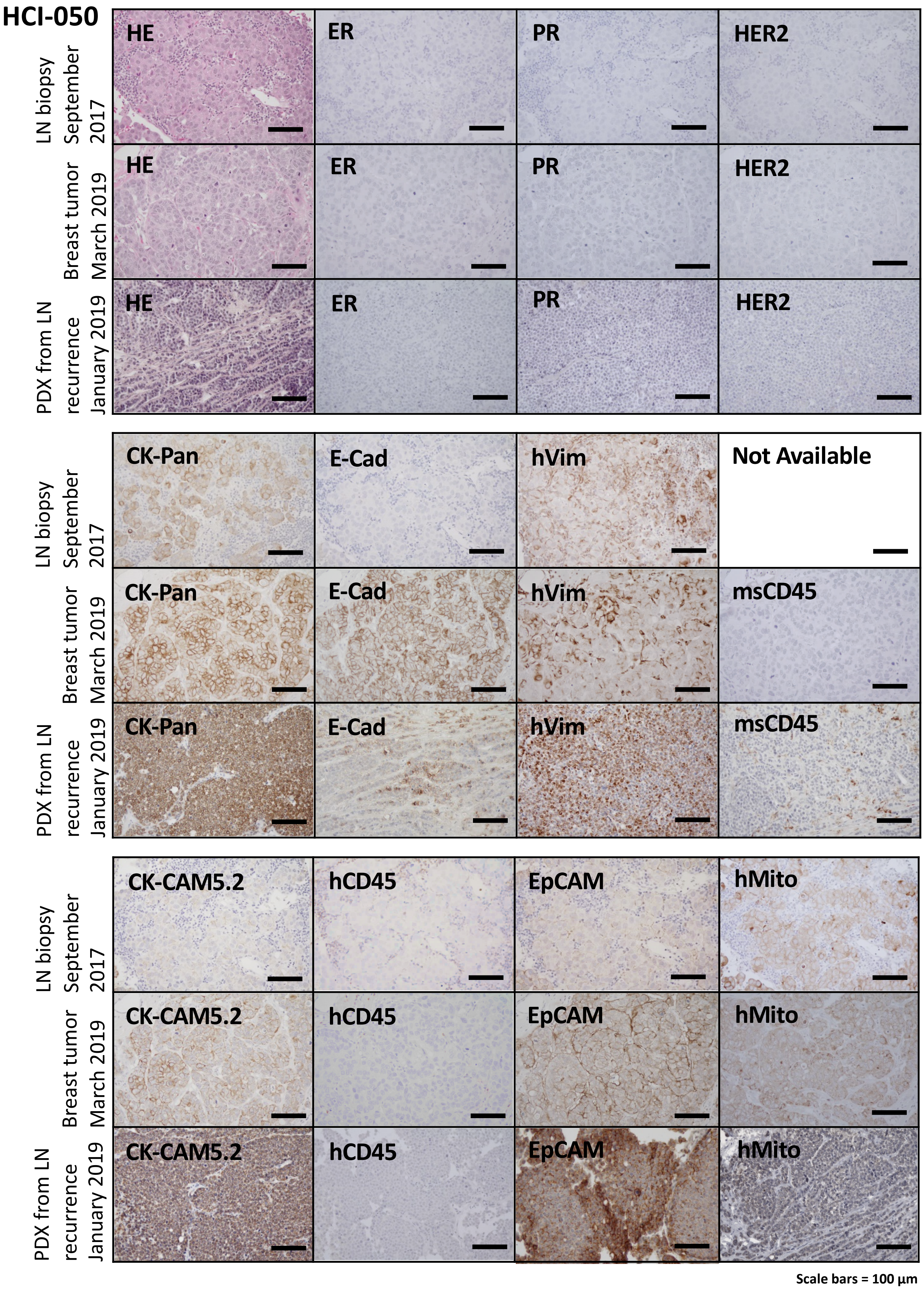

Supplementary Fig. 37

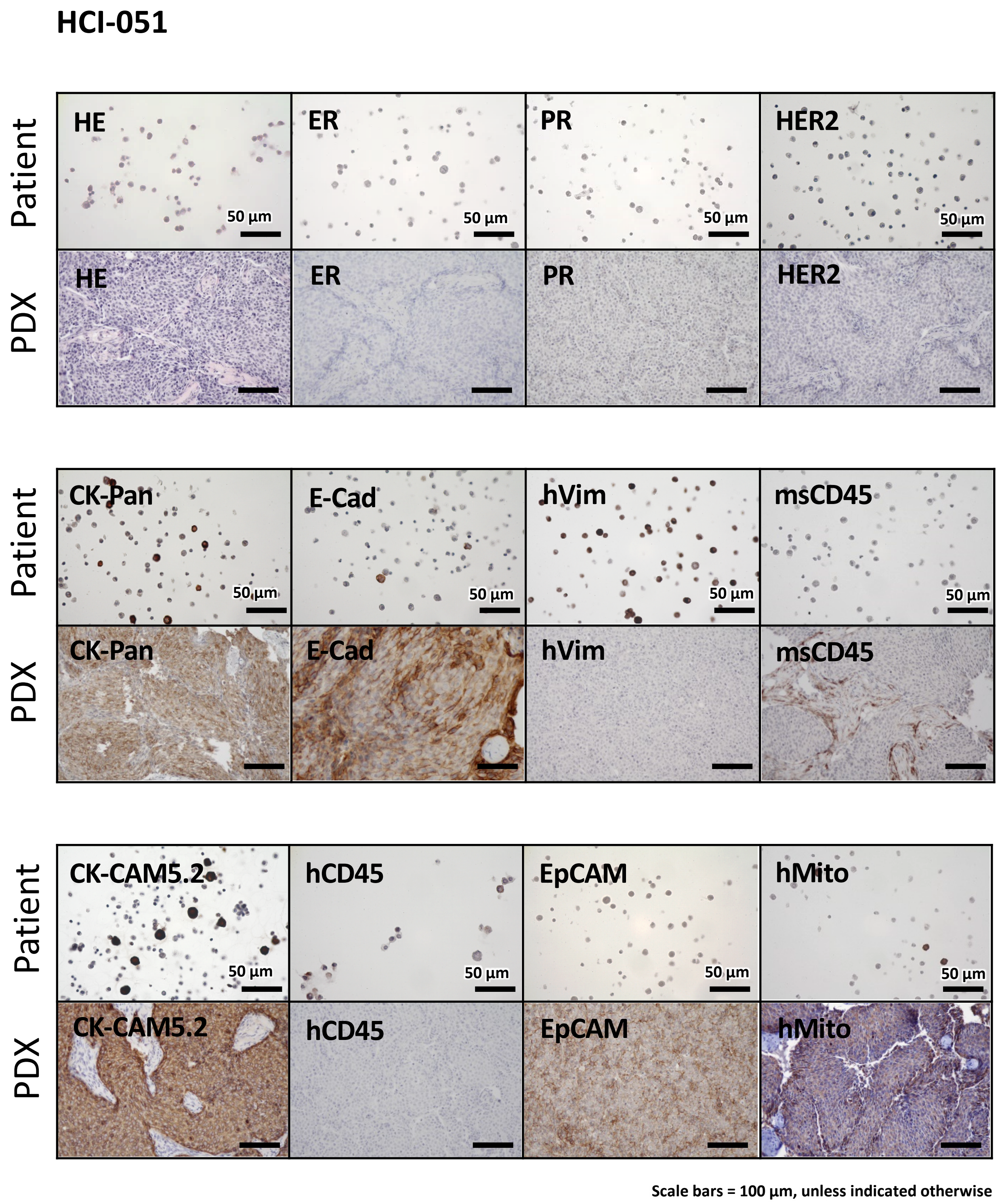

Supplementary Fig. 38

### HCI-052

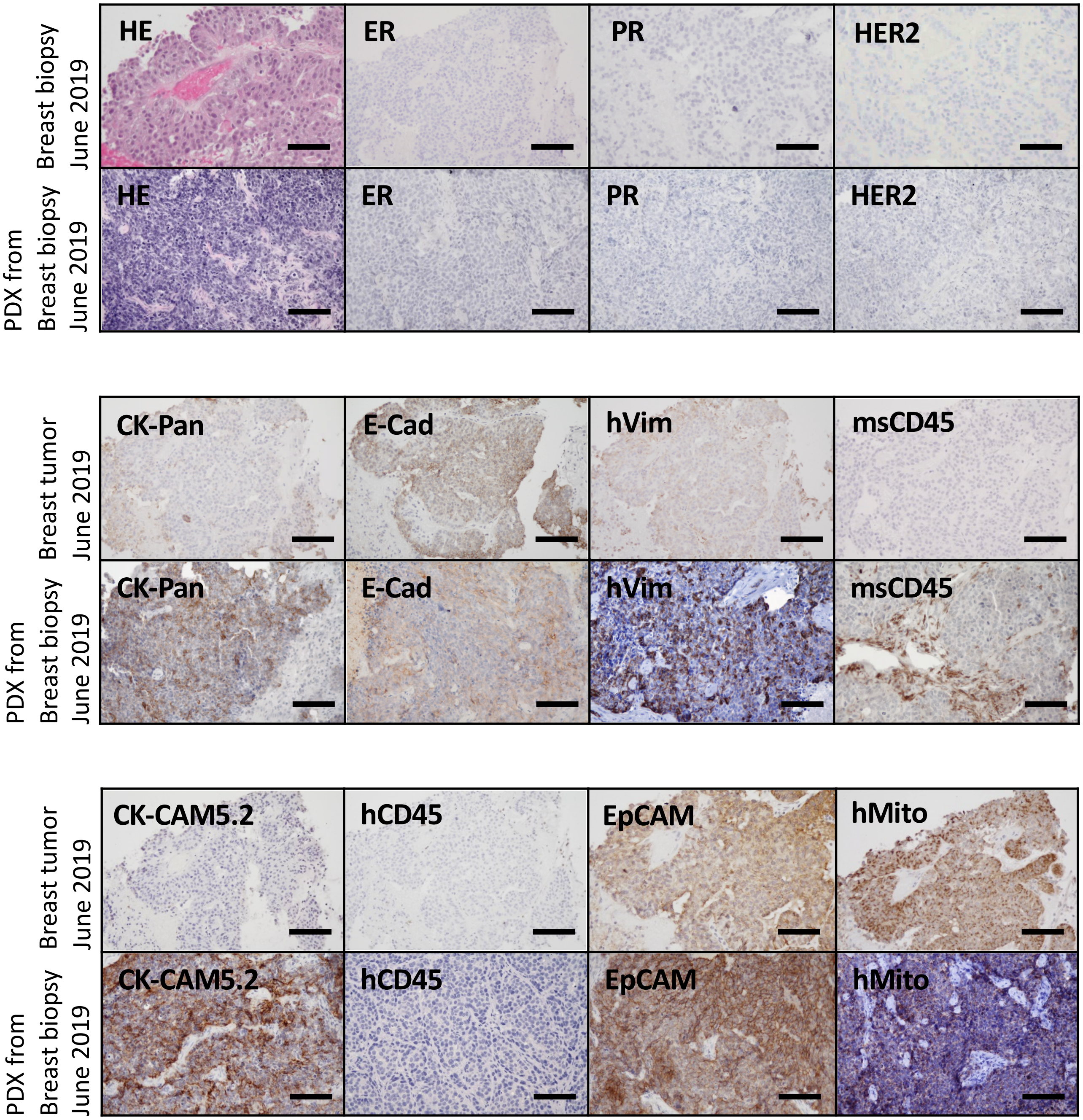


Scale bars = 100 µm

Supplementary Fig. 39

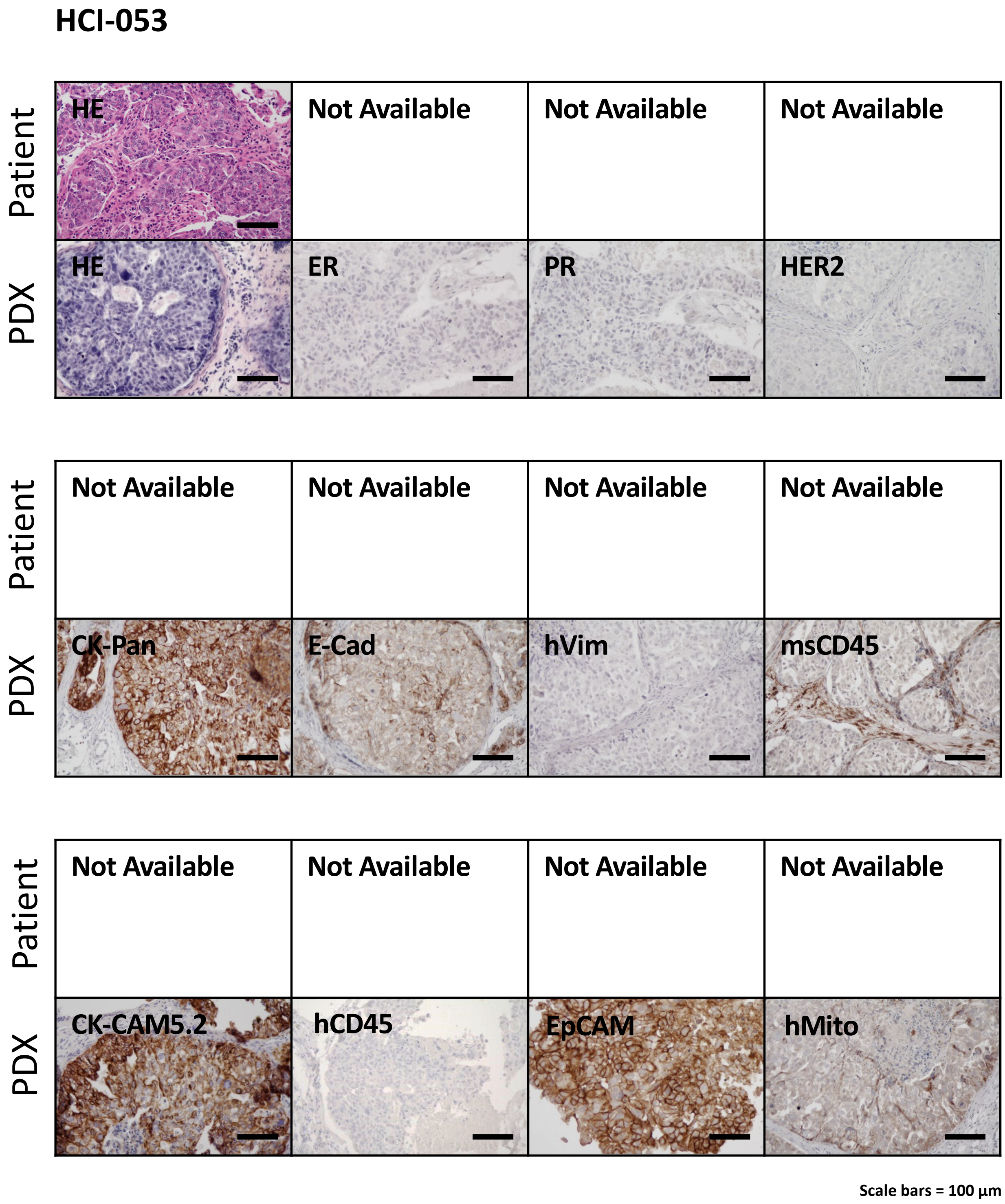

Supplementary Fig. 40

#### HCI-054

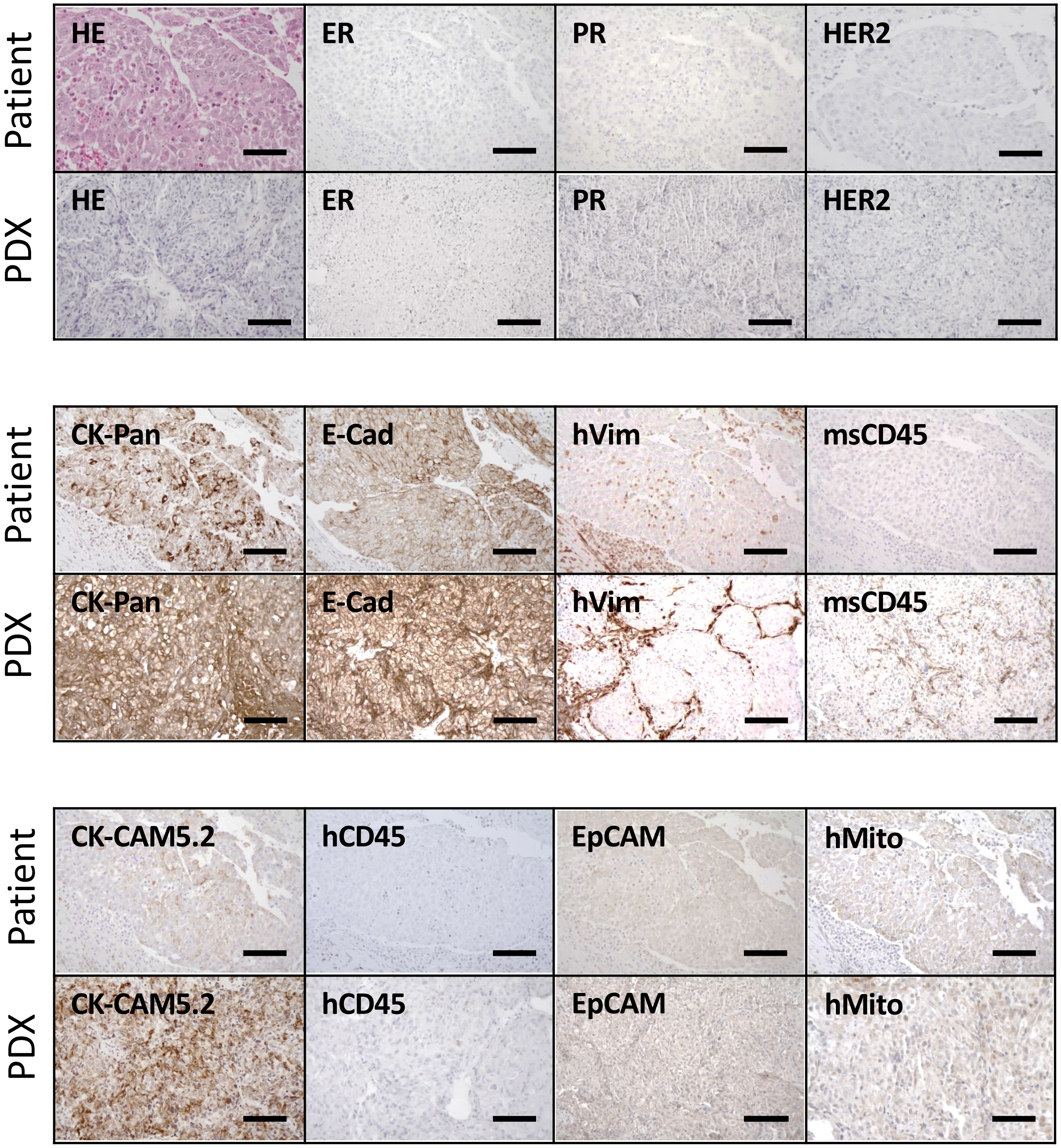


Scale bars = 100 µm

Supplementary Fig. 41

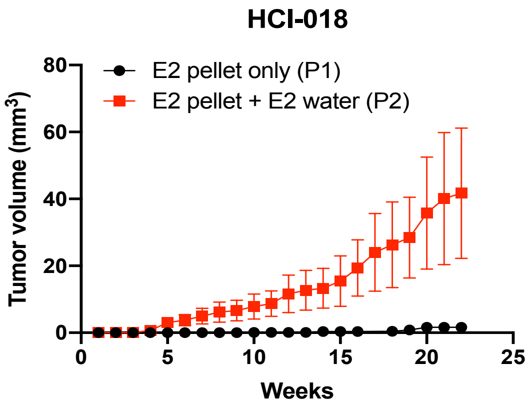

Supplementary Fig. 42

a

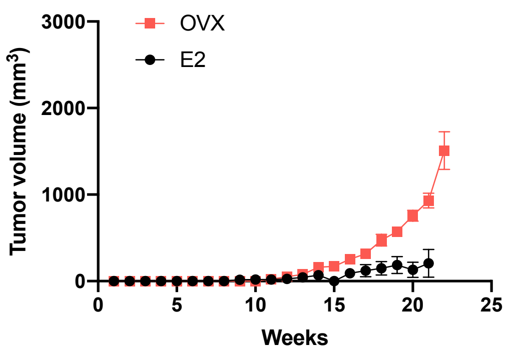


b

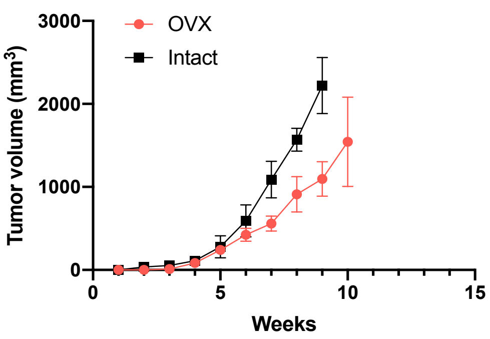


c

HCI-013EI

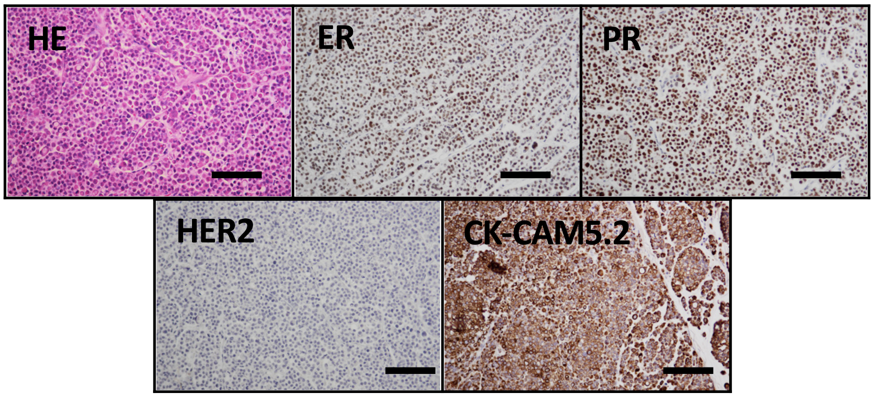


Scale bars = 100 µm

Supplementary Fig. 43

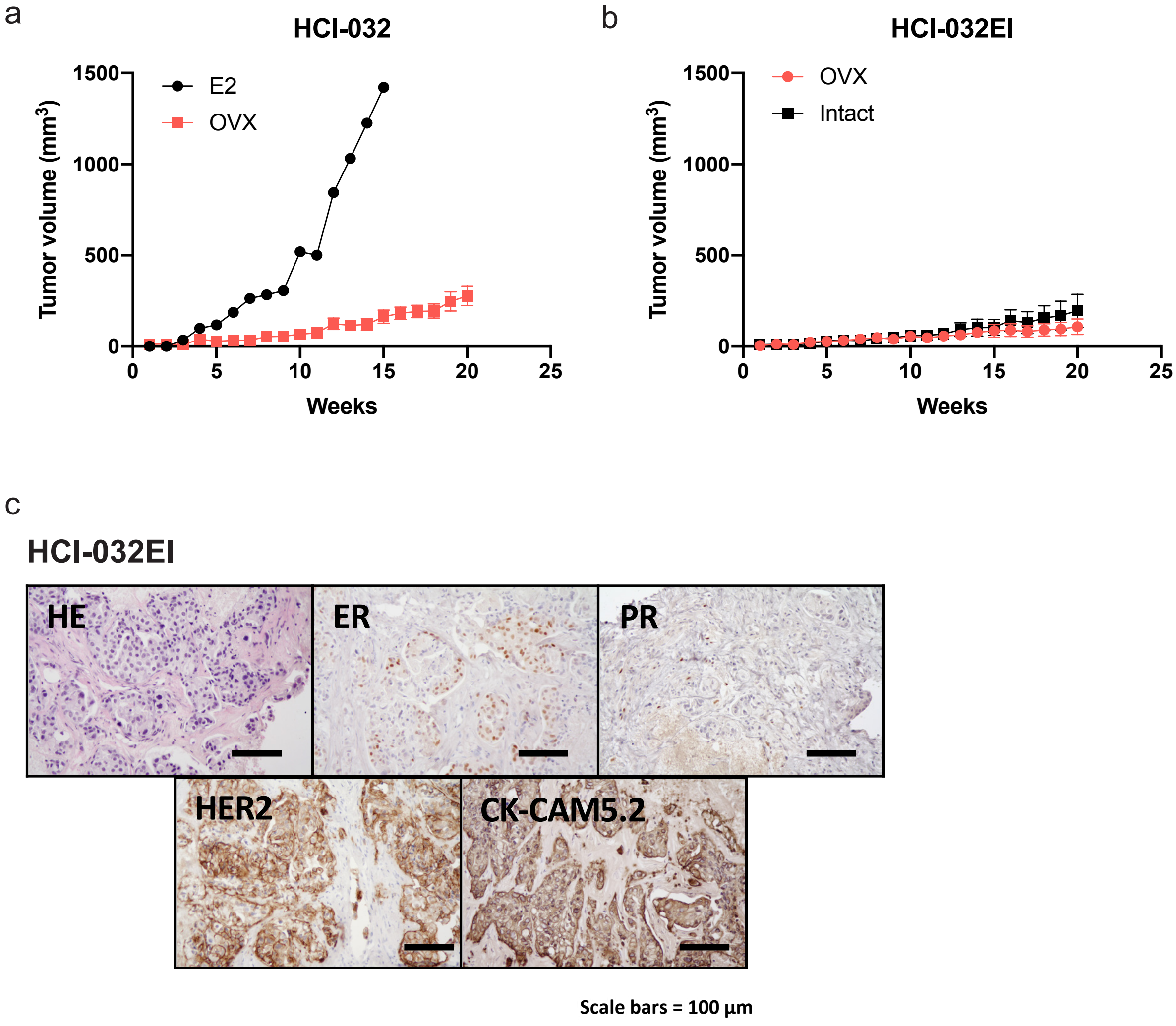

Supplementary Fig. 44

a

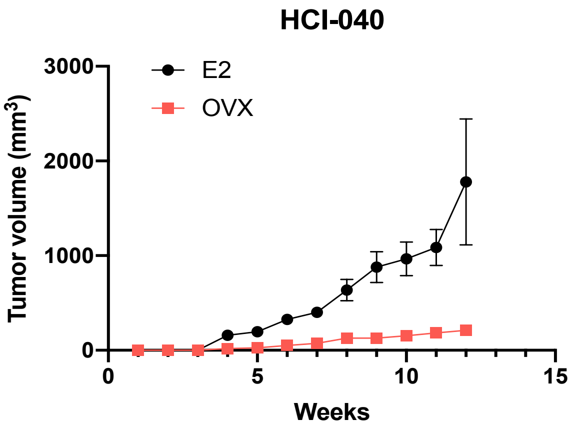


b

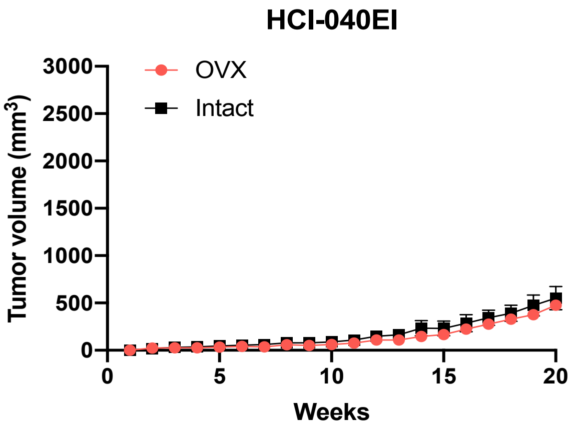


c

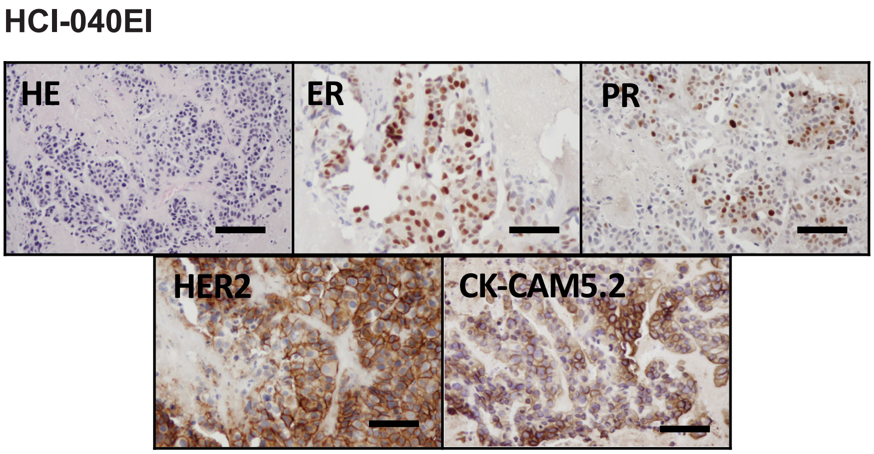


Scale bars = 100 µm

Supplementary Fig. 45

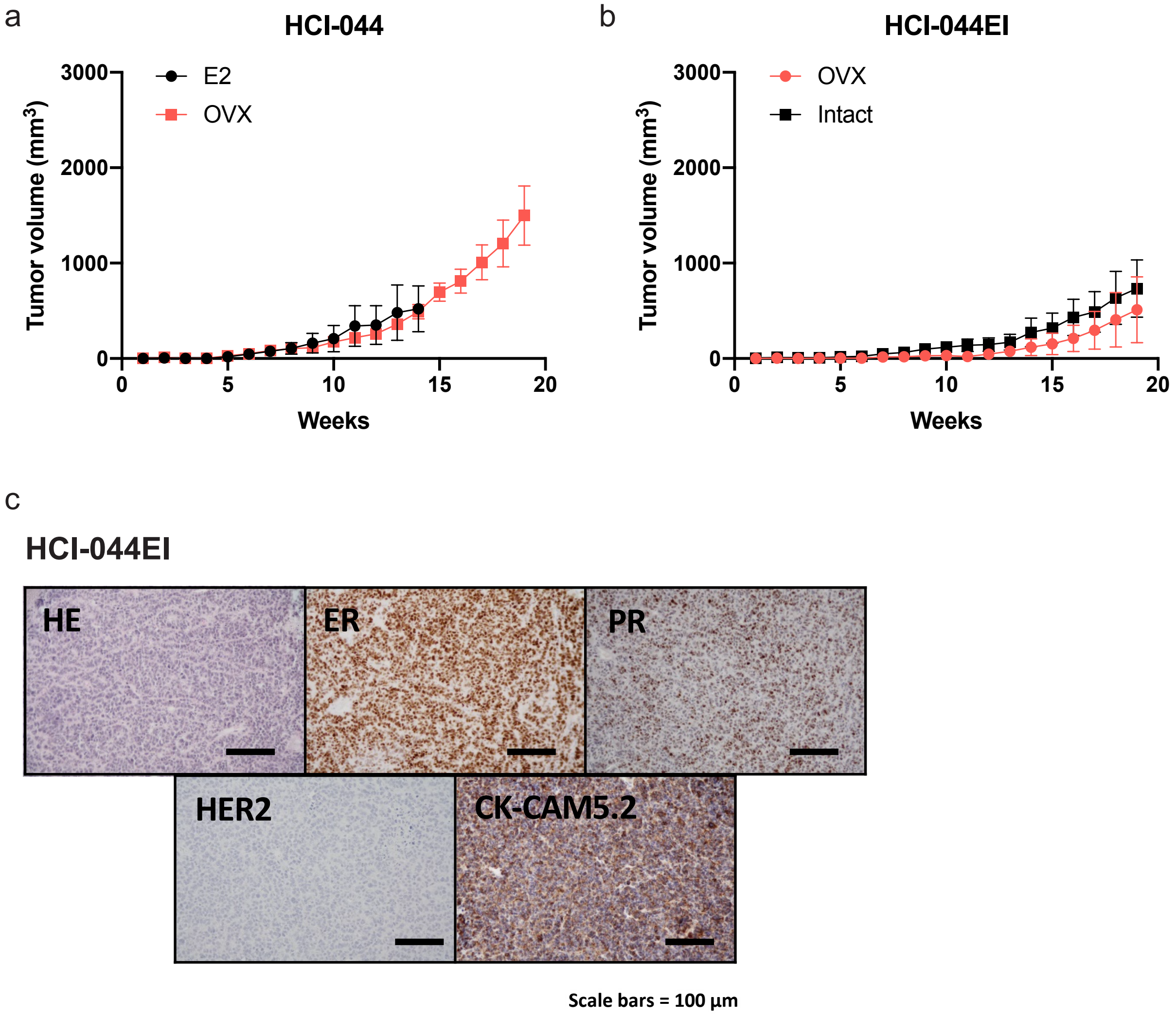

Supplementary Fig. 46

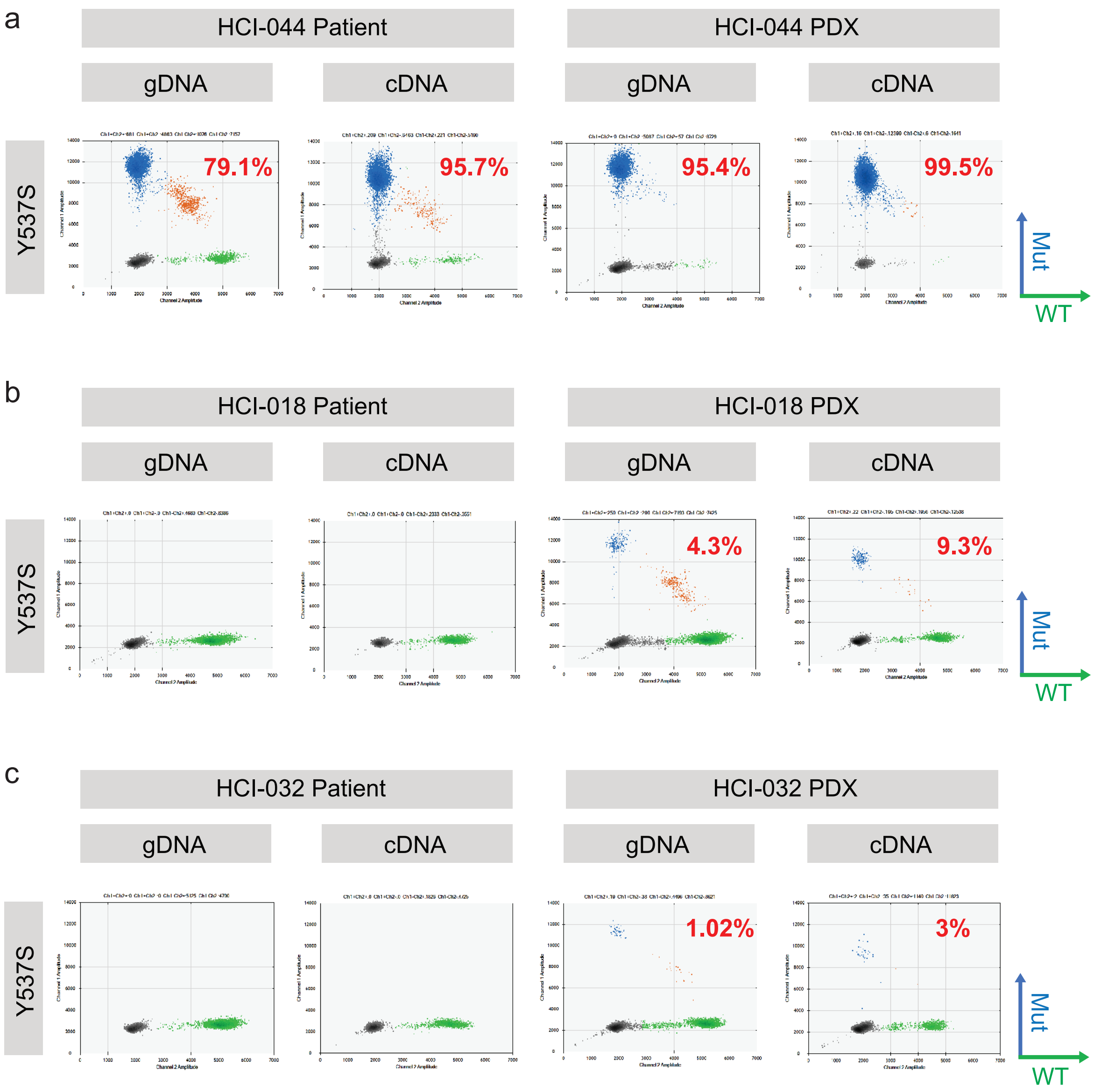

Supplementary Fig. 47

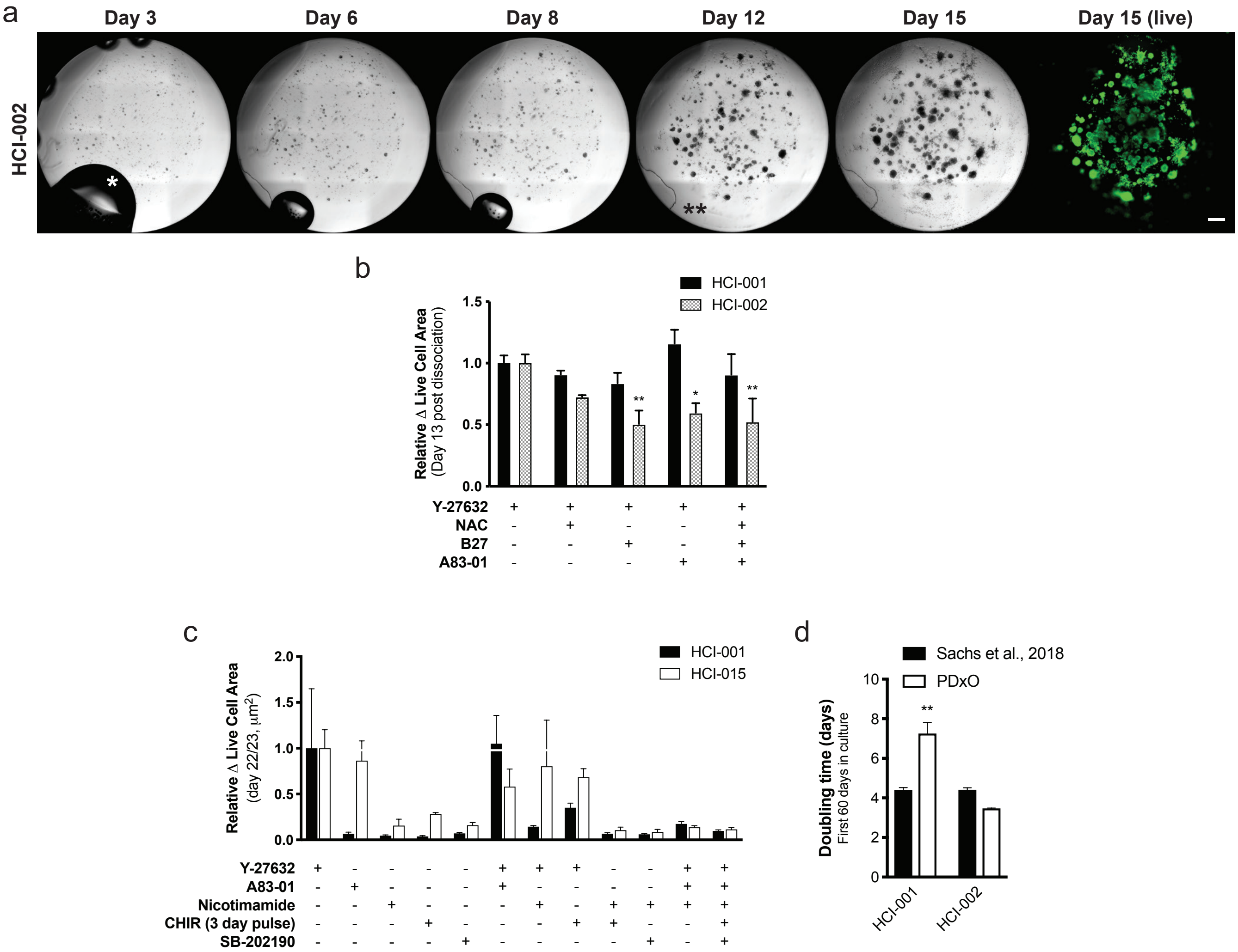

Supplementary Fig. 48

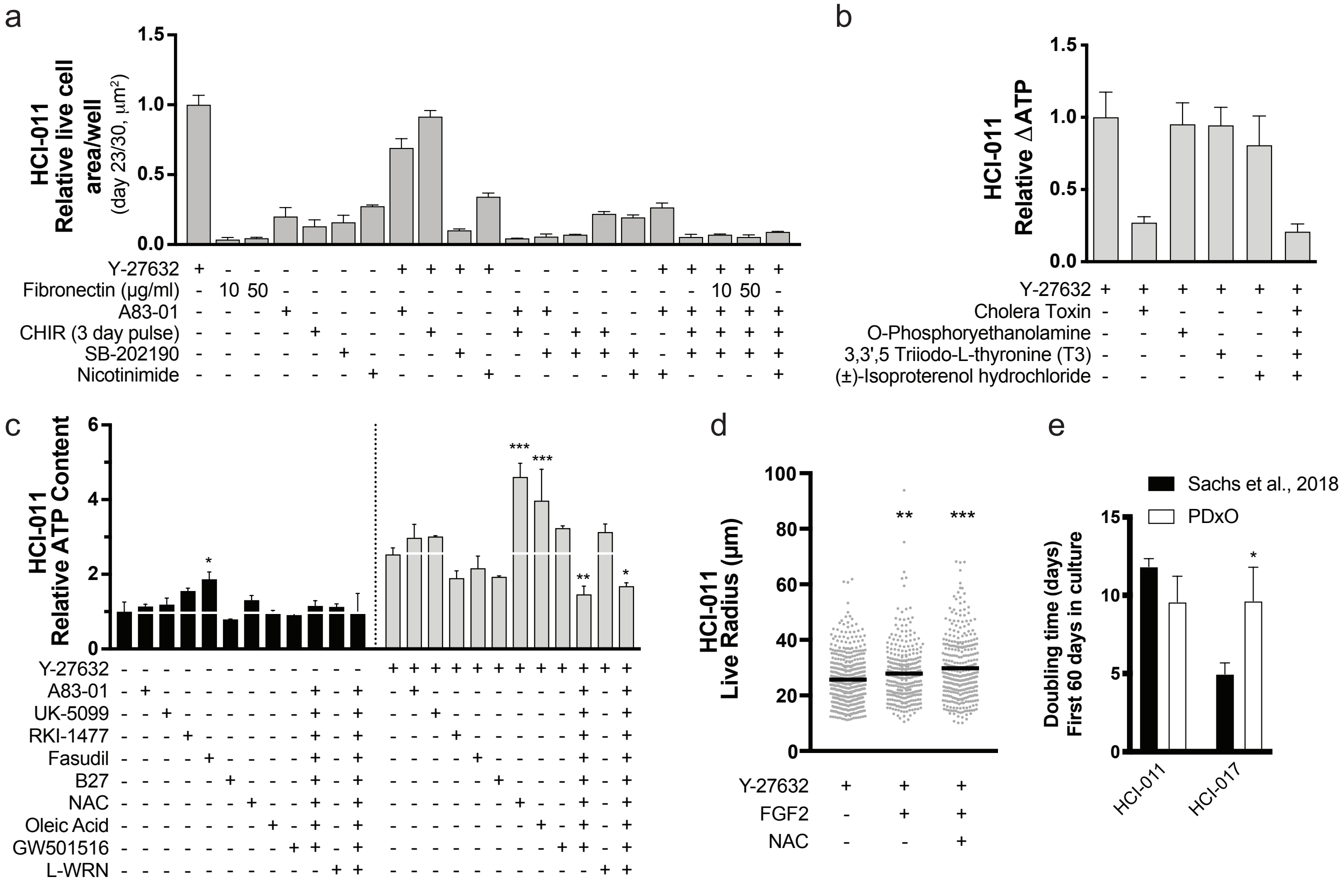

Supplementary Fig. 49

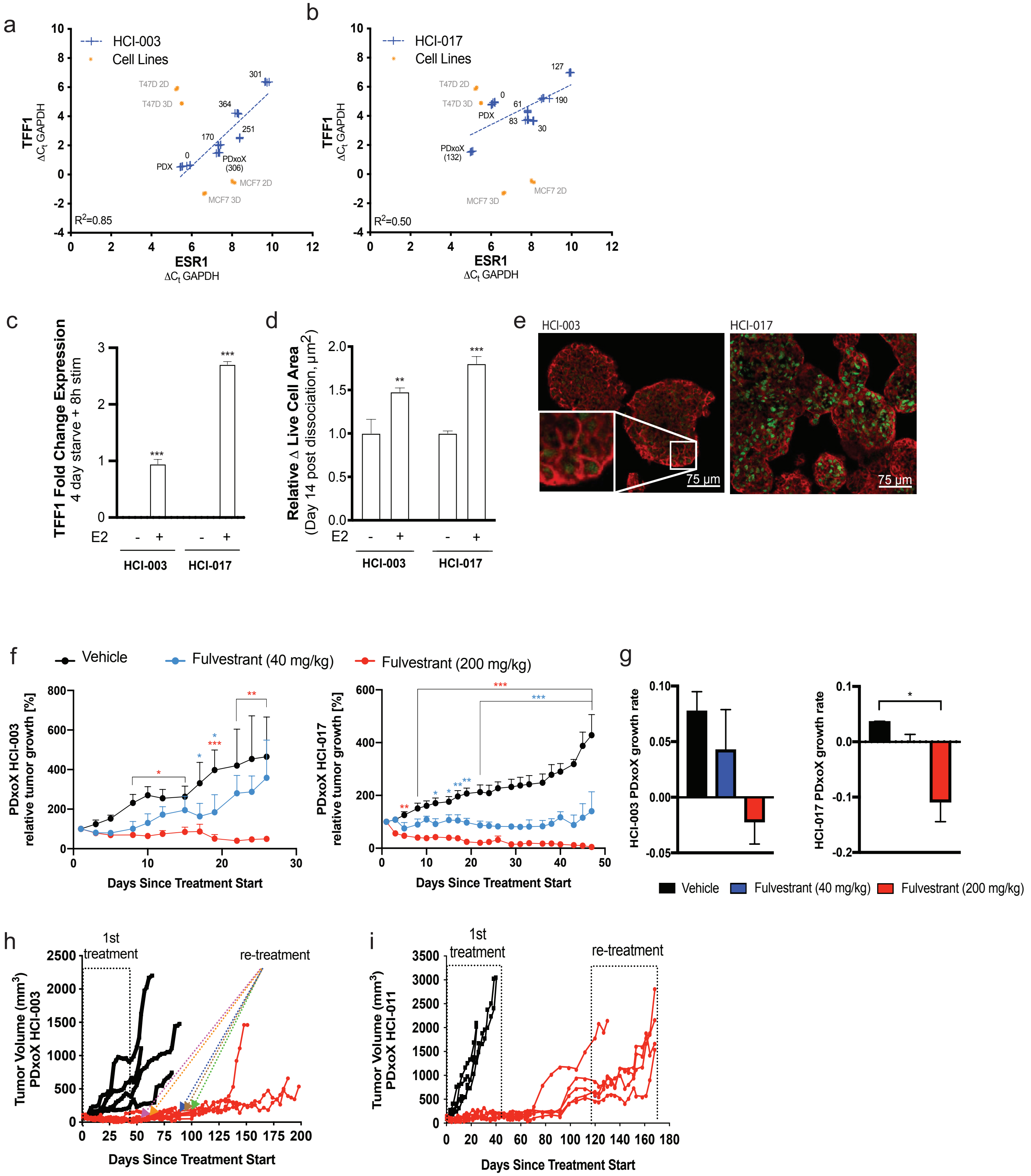

Supplementary Fig. 50

a

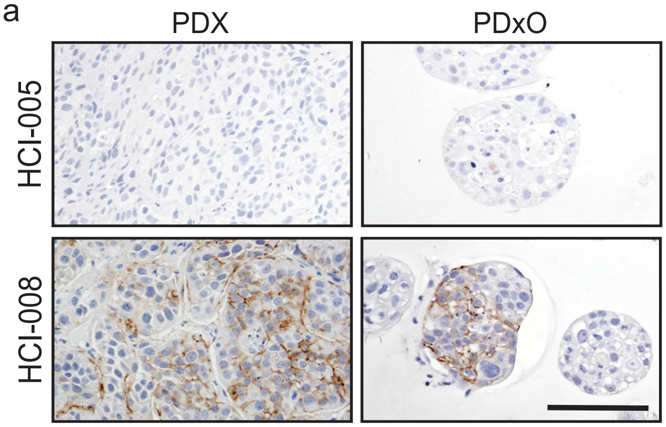


b

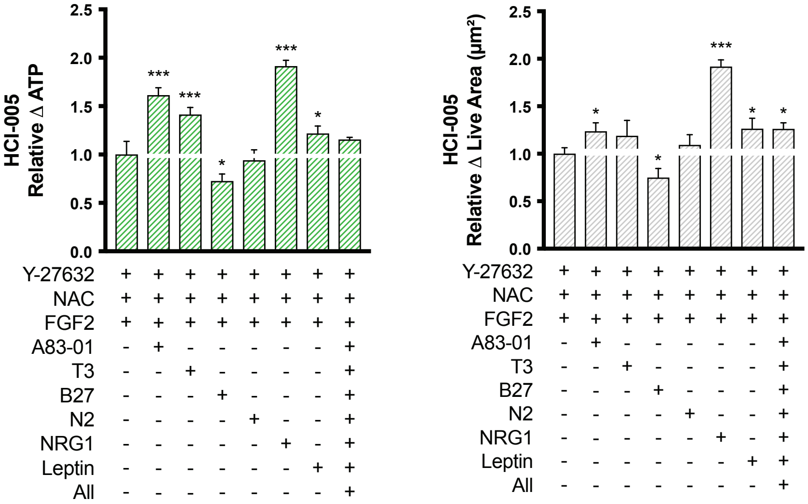

Supplementary Fig. 51

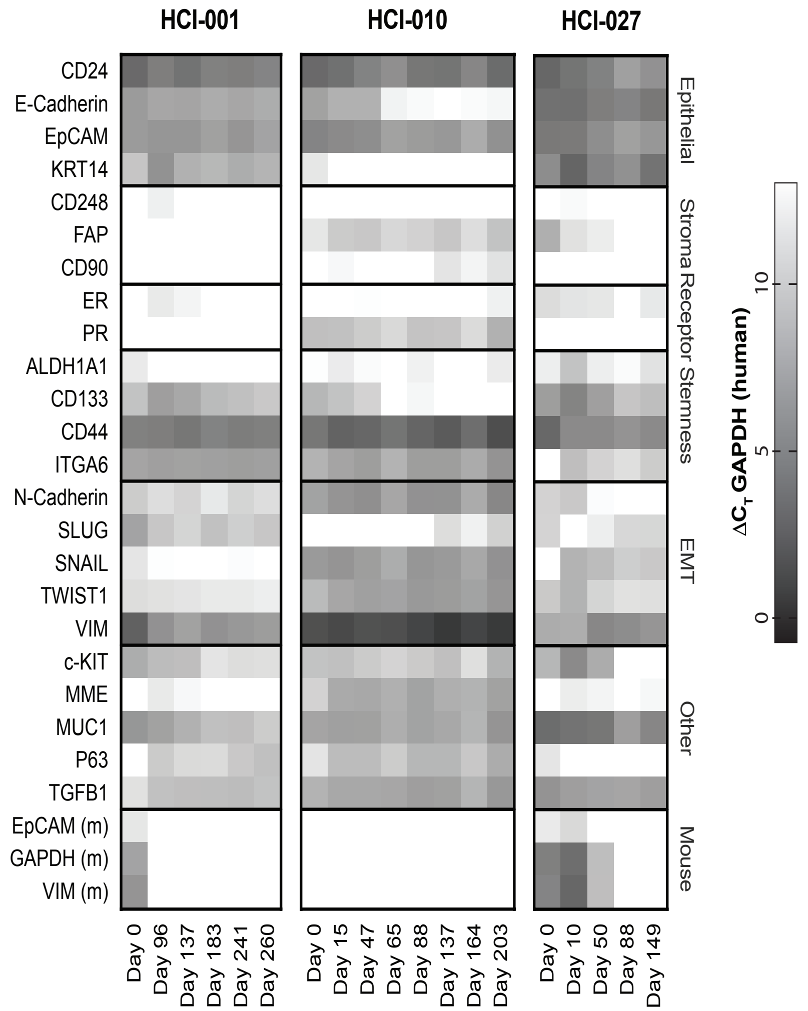

### Supplementary Fig. 52

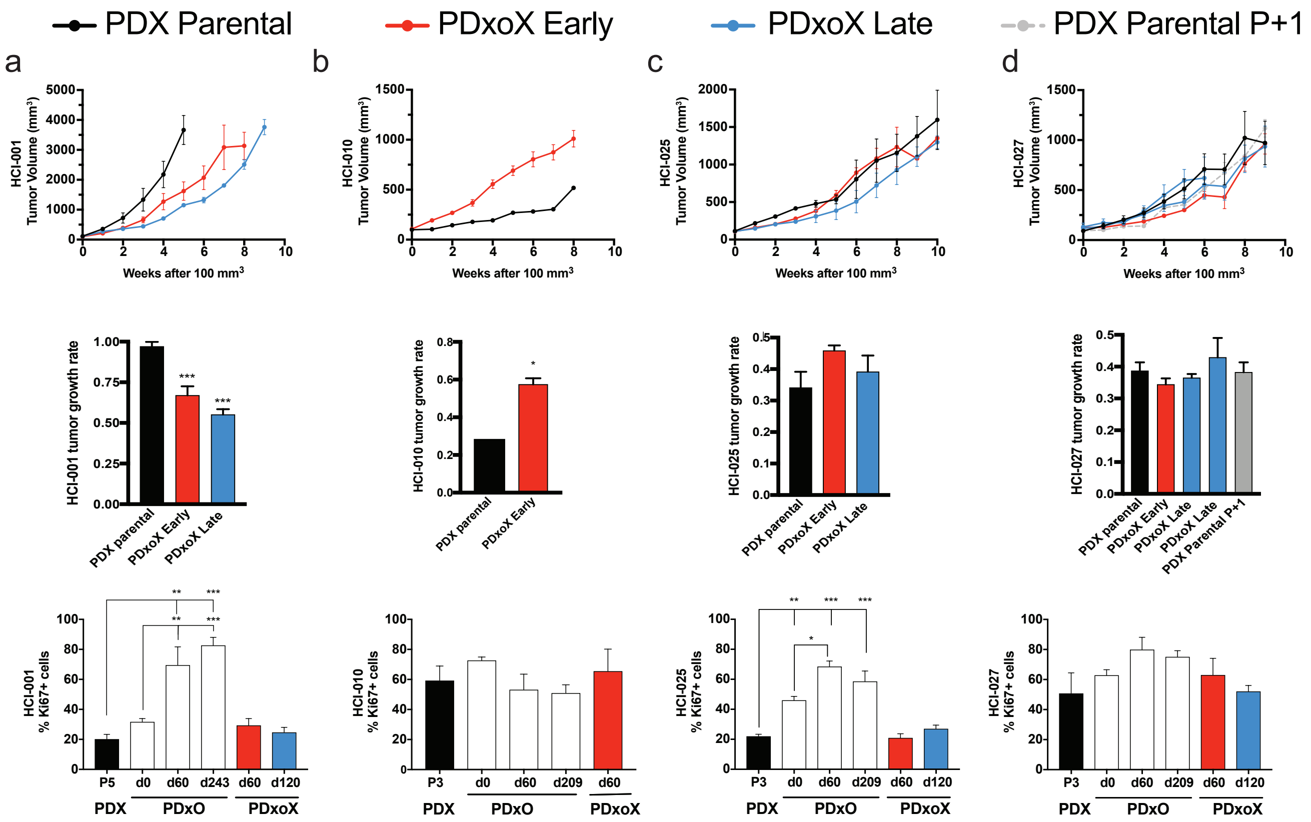

Supplementary Fig. 53

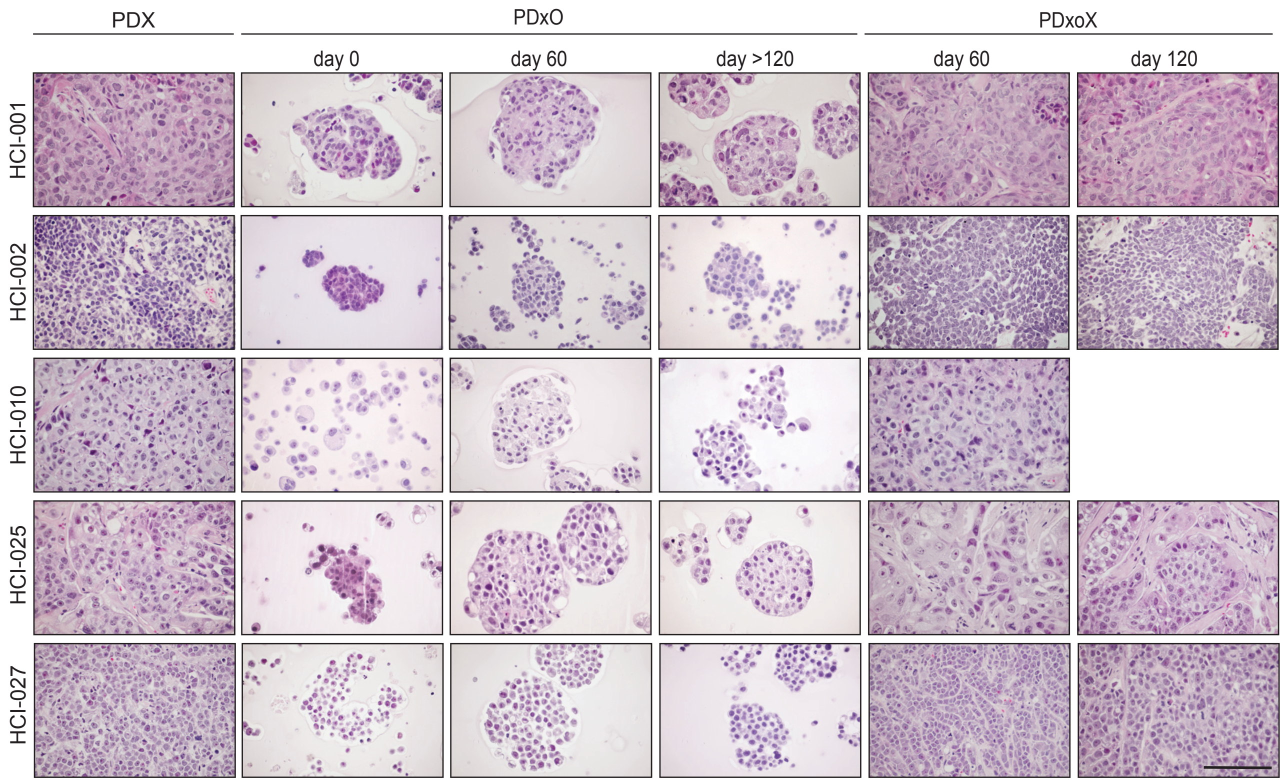

Supplementary Fig. 54

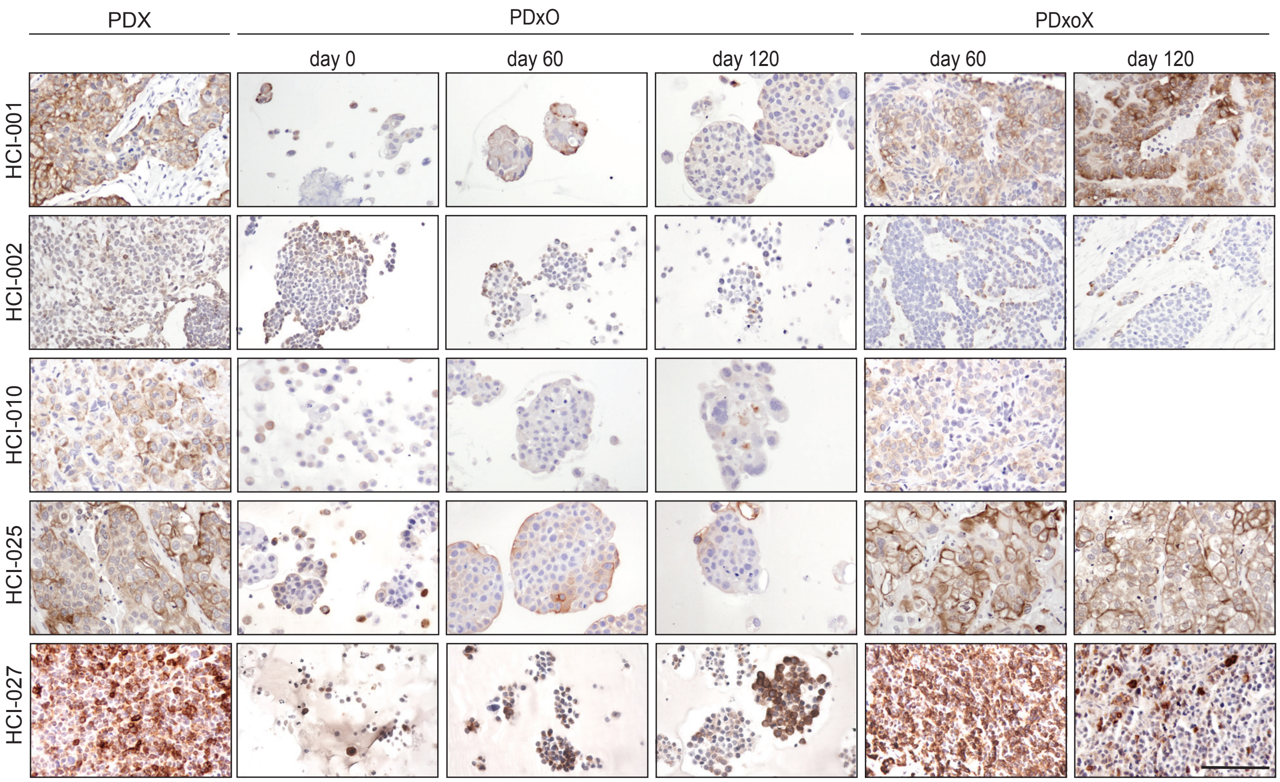

Supplementary Fig. 55

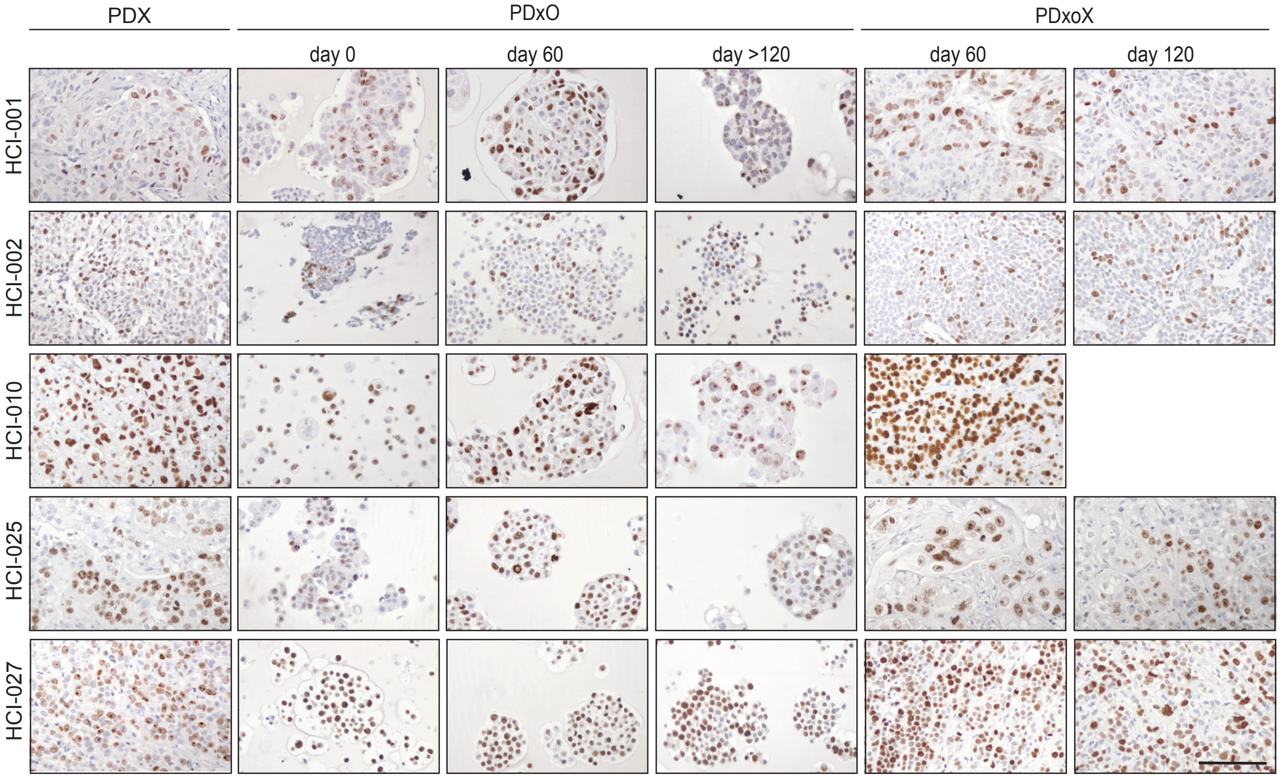

Supplementary Fig. 56

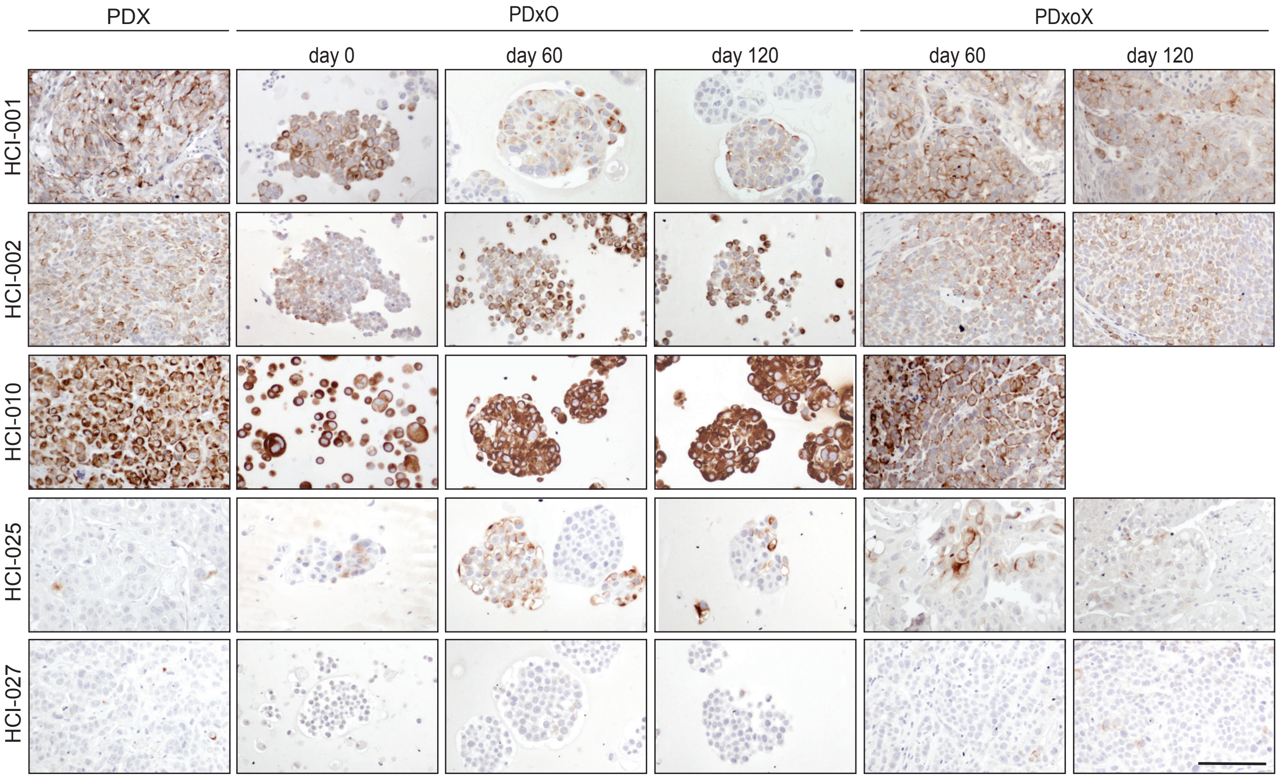

Supplementary Fig. 57

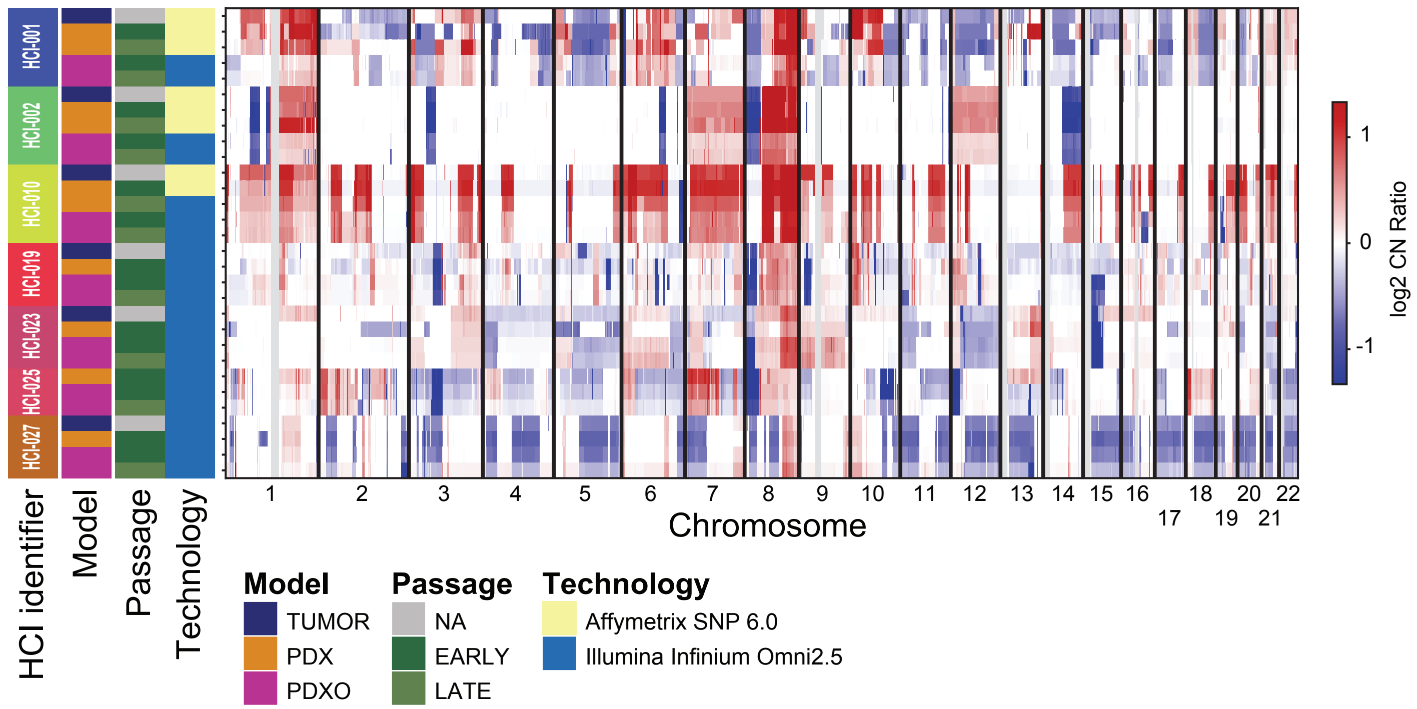


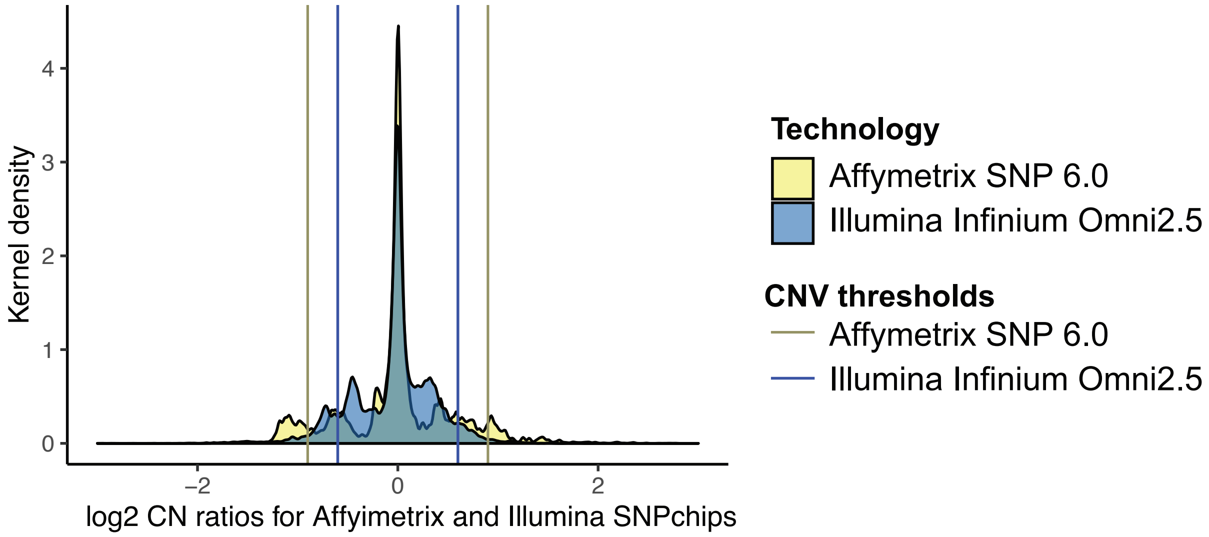

### Supplementary Fig. 58

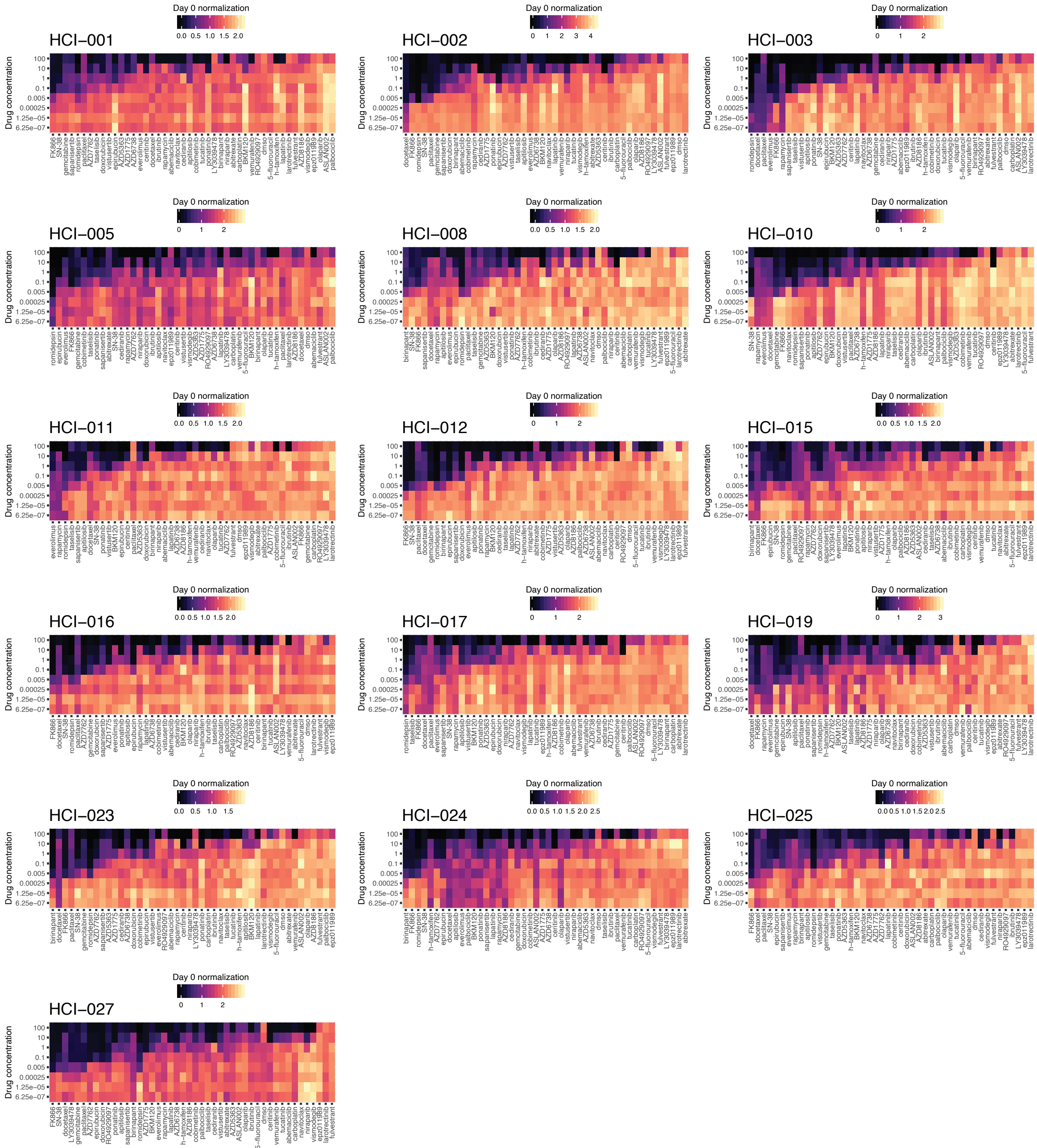

Supplementary Fig. 59

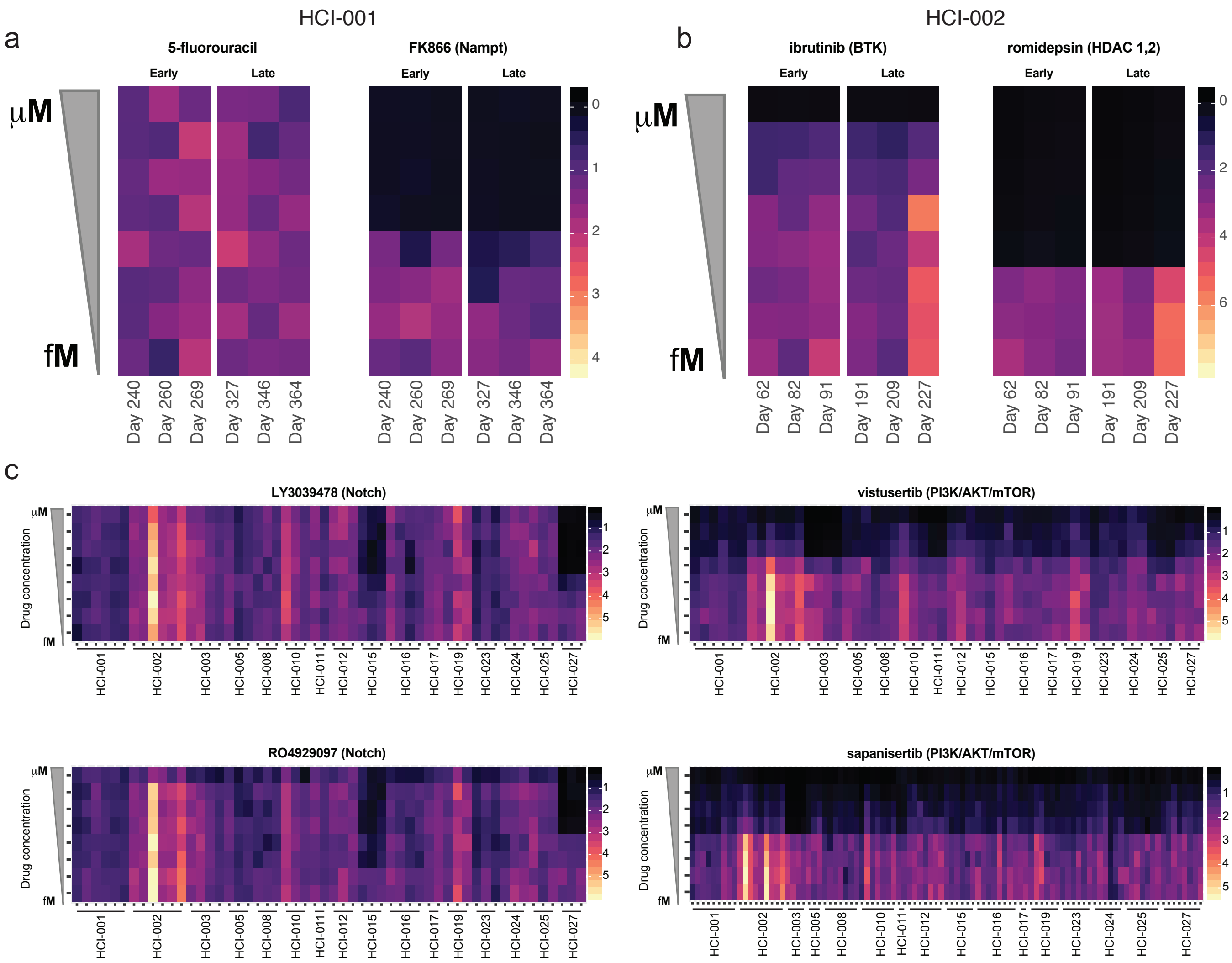

Supplementary Fig. 60

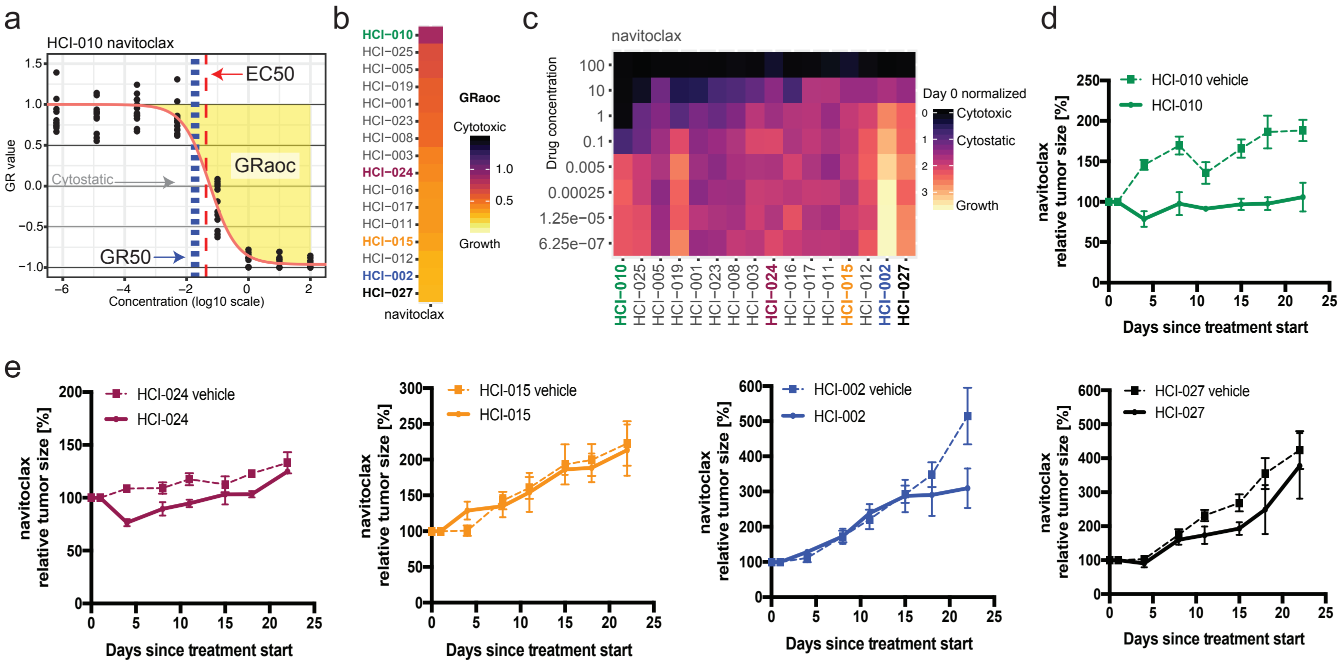

Supplementary Fig. 61

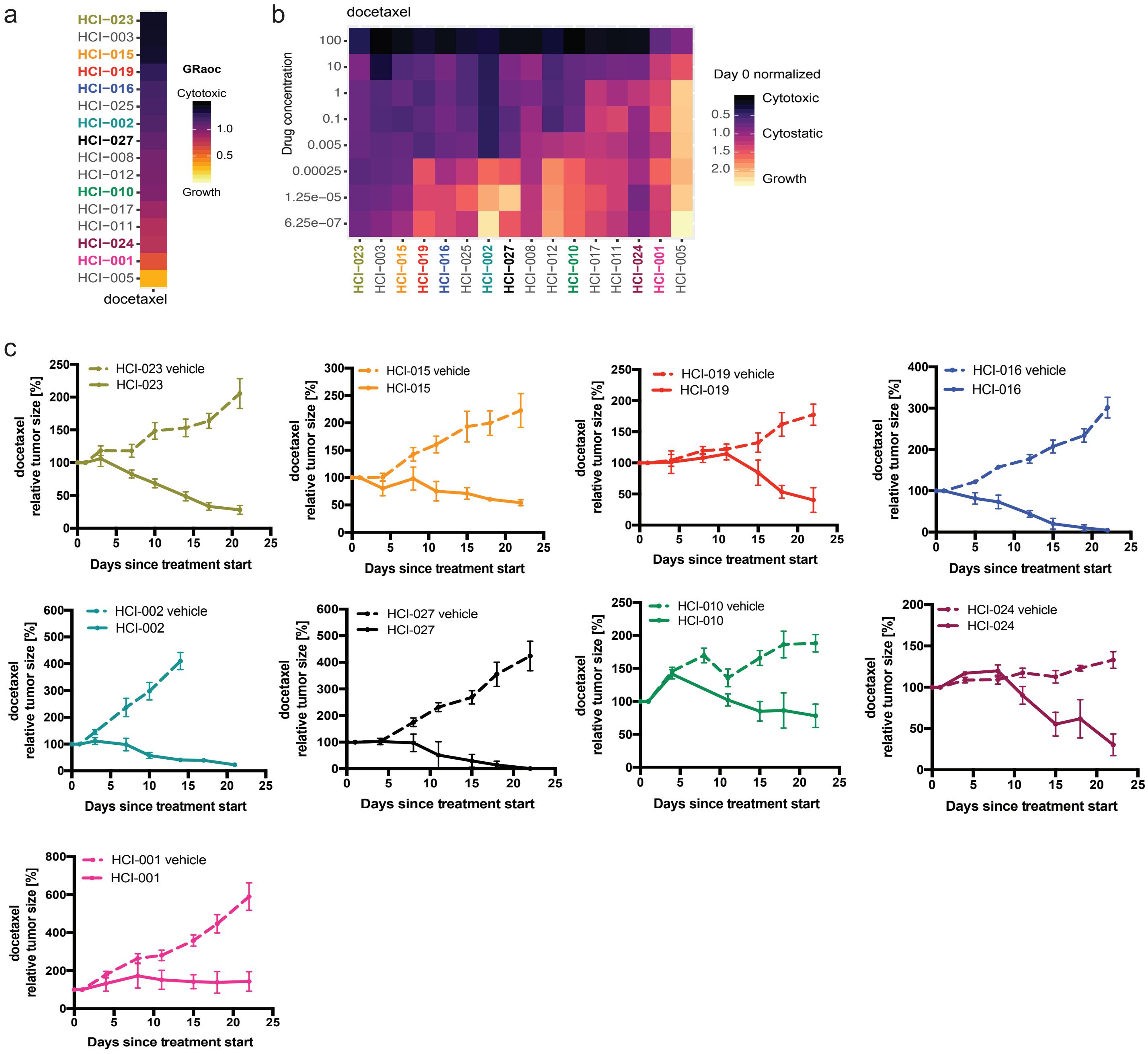

Supplementary Fig. 62

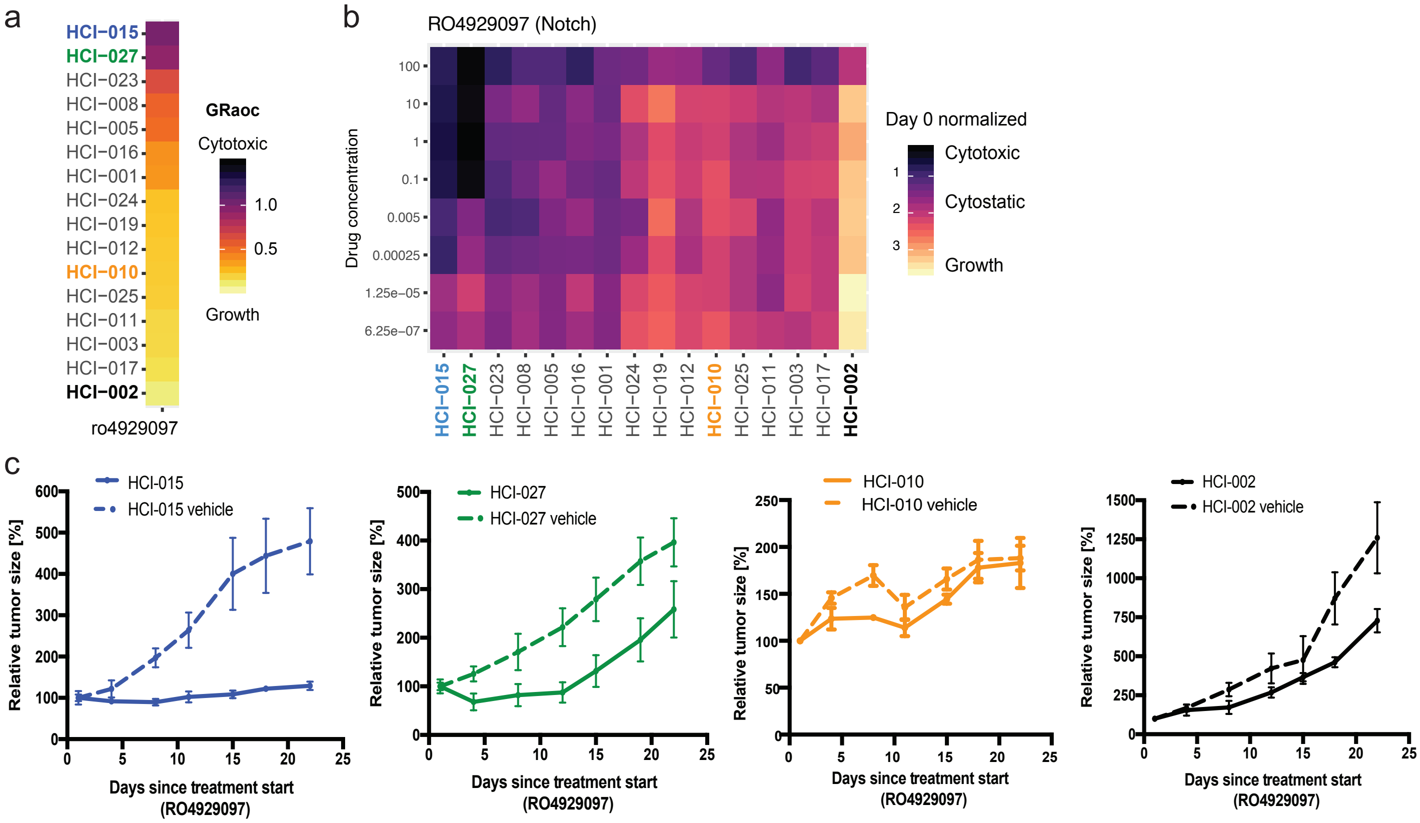

Supplementary Fig. 63

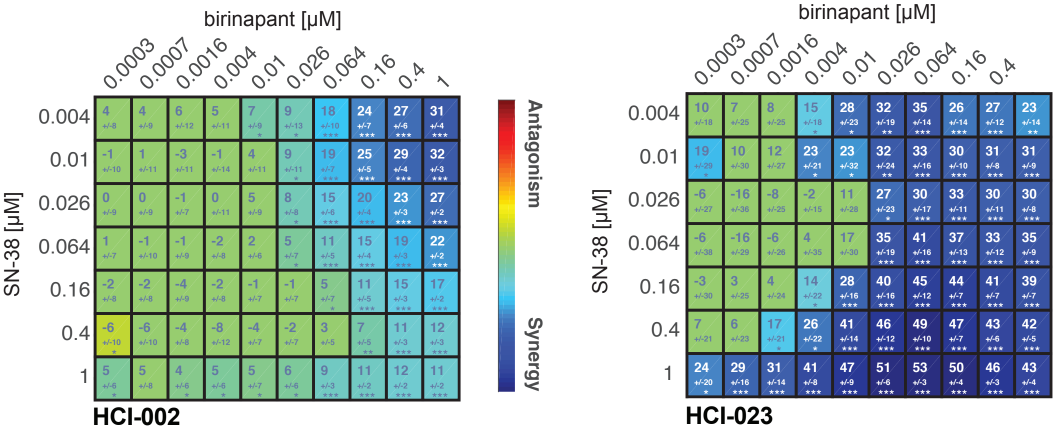

Supplementary Fig. 64

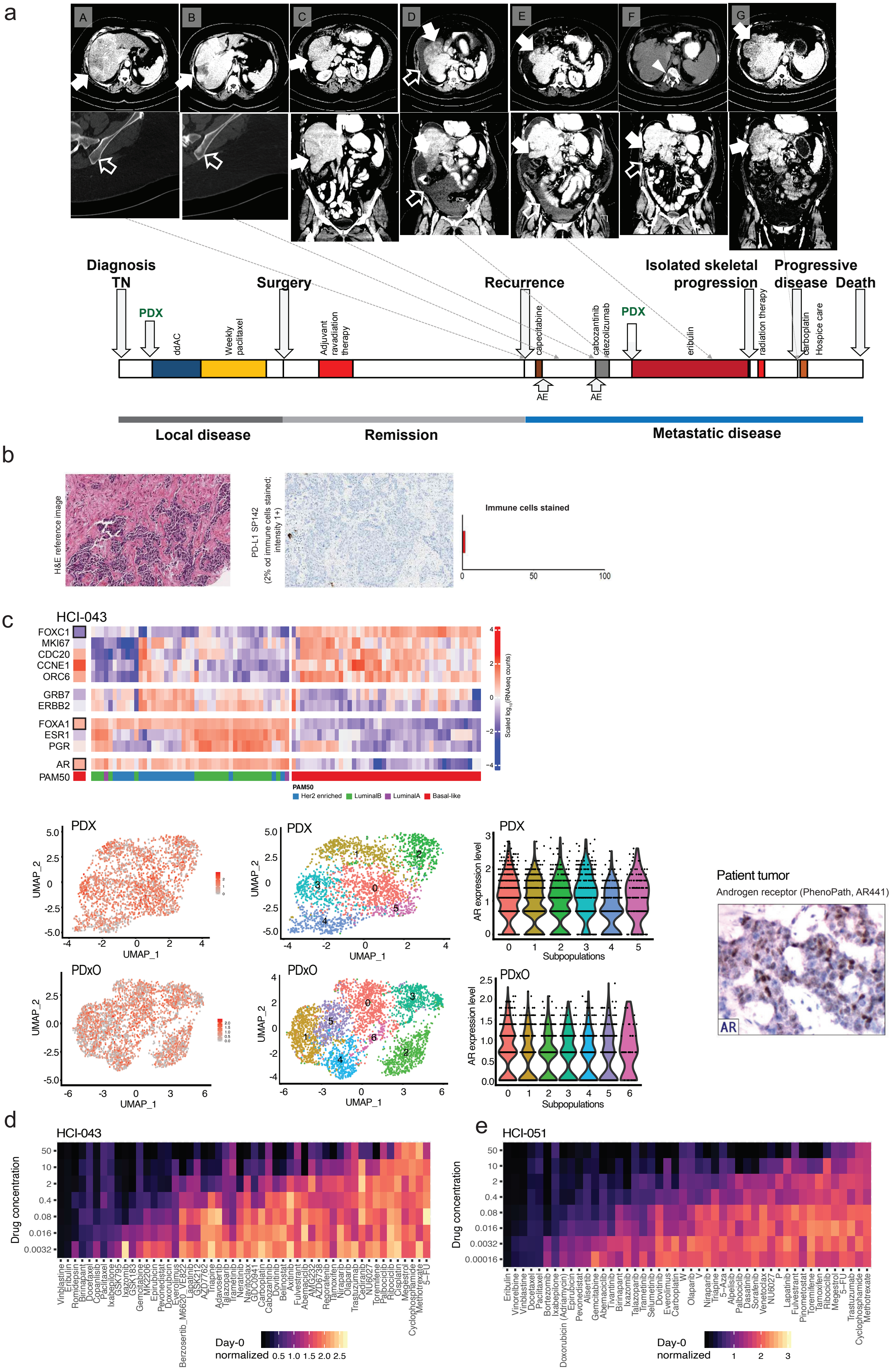
